# Novel misos shape distinct microbial ecologies: opportunities for flavourful sustainable food innovation

**DOI:** 10.1101/2024.02.28.582453

**Authors:** Caroline Isabel Kothe, Christian Carøe, Florent Mazel, David Zilber, Pablo Cruz-Morales, Nacer Mohellibi, Joshua Evans

## Abstract

Fermentation is resurgent around the world as people seek healthier, more sustainable, and tasty food options. This study explores the microbial ecology of miso, a traditional Japanese fermented paste, made with novel regional substrates to develop new plant-based foods. Eight novel miso varieties were developed using different protein-rich substrates: yellow peas, Gotland lentils, and fava beans (each with two treatments: standard and nixtamalisation), as well as rye bread and soybeans. The misos were produced at Noma, a restaurant in Copenhagen, Denmark. Samples were analysed with biological and technical triplicates at the beginning and end of fermentation. We also incorporated in this study six samples of novel misos produced following the same recipe at Inua, a former affiliate restaurant of Noma in Tokyo, Japan. To analyse microbial community structure and diversity, metabarcoding (16S and ITS) and shotgun metagenomic analyses were performed. The misos contain a greater range of microbes than is currently described for miso in the literature. The composition of the novel yellow pea misos was notably similar to the traditional soybean ones, suggesting they are a good alternative, which supports our culinary collaborators’ sensory conclusions. For bacteria, we found that overall substrate had the strongest effect, followed by time, treatment (nixtamalisation), and geography. For fungi, there was a slightly stronger effect of geography and a mild effect of substrate, and no significant effects for treatment or time. Based on an analysis of metagenome-assembled genomes (MAGs), strains of *S. epidermidis* differentiated according to substrate. Carotenoid biosynthesis genes in these MAGs appeared in strains from Japan but not from Denmark, suggesting a possible gene-level geographical effect. The benign and possibly functional presence of *S. epidermidis* in these misos, a species typically associated with the human skin microbiome, suggests possible adaptation to the miso niche, and the flow of microbes between bodies and foods in certain fermentation as more common than is currently recognised. This study improves our understanding of miso ecology, highlights the potential for developing novel misos using diverse local ingredients, and suggests how fermentation innovation can contribute to studies of microbial ecology and evolution.

## 1 Introduction

A fermentation renaissance is afoot. Against a backdrop of growing diet-related disease, insecure food systems, and the proliferation of undernourishing, bland, standardised foods, eaters around the world are rediscovering the ancient craft of fermentation for preservation, nutrition, flavour, and sustainability (Katz, 2016; Redzepi & Zilber, 2018; Steinkraus, 1997). As a way to produce intense, sometimes meaty flavours and unlock nutritional value in plant substrates and byproducts, fermentation has also recently been enlisted to facilitate the green transition (Capozzi *et al*., 2021). For many places in the world, shifting toward more plant-based diets is one of the best ways to mitigate climate change (Willett *et al*., 2019; Xu *et al*., 2021). Making plant-based products taste as satisfying as animal-based products, and in culturally appropriate ways, is therefore one of the main challenges to help reach this urgent goal. While many cultures, such as across Asia and Africa, already have diverse ways of producing rich, savoury plant-based foods through fermentation (Obafemi *et al*., 2022; Steinkraus, 1994; Tamang & Kailasapathy, 2010), there are not as many in Western countries. There is thus great potential in Western food cultures learning from these practices, drawing on this traditional knowledge to produce novel plant-based products using local substrates, yielding diverse flavours that are both familiar and new (Andersen *et al*., 2022; Damsbo-Svendsen *et al*., 2017; Rozin, 1976; Waehrens *et al*., 2023).

Here we experiment with miso, a staple of Japanese cuisine, to illustrate how we might combine existing fermentation knowledge with novel regional substrates to yield new, diverse sources of savoury, plant-based foods. Miso is a fermented paste made by combining cooked soybeans, kōji (the filamentous fungus *Aspergillus oryzae* grown on rice, barley, or soybeans), and salt. There are many styles of miso, which can vary according to the ratio of the ingredients, the fermentation time, and where and by whom it is produced (Shurtleff & Aoyagi, 1981). Miso is used in many Japanese dishes, such as soups, sauces, and marinades, to add richness, depth of flavour, and savouriness, often known as *umami* (Inoue *et al*., 2016). More recently, chefs, home cooks, and fermenters have been experimenting with making misos from other, non-traditional ingredients that can fulfil the same functions, such as chickpeas, nuts, seeds, and other plants (Felder *et al*., 2012a; Kusumoto *et al*., 2021; Redzepi & Zilber, 2018; Reiß, 1993). Some have even been experimenting with introducing additional techniques to the miso process, such as nixtamalisation—an ancient process from Mesoamerica in which maize, and nowadays other substrates, are cooked in an alkaline solution to increase their nutrient bioavailability, culinary functionality, and flavour. We have carried out our experiment at an experimental, industry-leading restaurant in Copenhagen called Noma, in the context of the New Nordic Cuisine, a culinary movement of the past two decades that has sought to rediscover and develop the edible biodiversity of the Nordic region through cooking (Evans & Lorimer, 2021). This context informed the substrates we selected. The aim of this research is to investigate the microbial community structure and diversity of novel misos made with different substrates and produced in different regions, to understand more about fundamental miso ecology and to facilitate the development of new, regionally diverse, and culturally appropriate plant-based foods.

As popular interest in global fermentation techniques has exploded, scientific attention to fermented foods has also grown, as part of a larger research focus on the human microbiome and health, and facilitated by the decreasing cost of Next Generation DNA Sequencing (NGS) technologies (Jahn *et al*., 2023; Lorimer, 2016, 2020; Mardis, 2017; Yong, 2016). While studies on the microbial ecology of many fermented foods are becoming more common (Landis *et al*., 2022; Pasolli *et al*., 2020; Walsh *et al*., 2022; Yap *et al*., 2022), there still exists very little English-language literature on miso ecology. A few older reports exist that use only culture-dependent methods (Hesseltine, 1983), but the few taxa identified *‘do not exclude certain other halophilic yeasts and bacteria from growing in or on the fermenting paste’* (Hesseltine, 1983: 589), as microbial isolation-based taxonomy is biased toward culturable species. Onda *et al*. published a handful of studies on miso bacteria using 16S rRNA sequence analysis for identification (Onda et al., 2002, 2003a; Onda *et al*., 2003b), however these only involved cultivable bacteria. Other authors have studied bacterial and fungal communities in miso and Chinese analogues (Kim *et al*., 2010), as well as in soy sauce (Tanaka *et al*., 2012b), using polymerase chain reaction denaturing gradient gel electrophoresis (PCR-DGGE). NGS methods have been employed to analyse the microbial communities in *doenjang*, Korean fermented soybean pastes (Nam *et al*., 2012), but these products are made differently from miso.

Allwood *et al*. provide a recent review of studies on the fermentation and microbial communities of Japanese kōji and miso (Allwood *et al*., 2021). After reviewing the scant literature in English on miso ecology, they conclude that *‘due to the limitations of these studies, lack of details available regarding the ingredients and fermentation conditions of the purchased miso samples, along with the different methodologies used, it is difficult to draw any conclusions from these studies to identify a typical microbial profile for miso.’* (ibid.: 2200). As our focus here is on novel misos, we are not able to address Allwood *et al*.’s call, however worthwhile, to identify a typical microbial profile for traditional misos. We hope, however, that the NGS methodology we employ could also be used to develop similar knowledge of the microbial community structure and diversity of diverse traditional misos.

In a recent exploratory study, we used metagenomics to identify the microbiota of misos made with substrates other than soybeans (Kothe *et al*., 2023). In this study, while we observed that the microbial composition of each novel miso was shaped differently by its substrate, the absence of replicates limited our ability to say this conclusively and to study potential variability across misos of the same type. Based on these preliminary results, we conducted here an extended study aiming to enhance our understanding of these products, with new substrates, batch replicates, temporal parameters, treatments, and fermentation sites. We hope that this study will complement the previous one and can offer a useful contribution to the literature, facilitating a better understanding of miso ecology and supporting the production of both traditional and innovative misos.

## 2 Methodology

### 2.1 K**ō**ji preparation

Rice (*Oryza sativa*, Kokuho Rose, Nomura & Co., USA) and pearled barley (*Hordeum vulgare*, Lantmännen Cerelia A/S, Vejle, DK) were soaked in water overnight (2 parts water: 1 part dry grains), and steamed at 100°C for 40 min in stainless steel trays using a Rationale Combi oven. The grains were then broken up with gloved hands and cooled at room temperature until they reached 37°C, at which point 0.2% of pure commercial albino white rice kōji spores (from Bio’c, Japan) was used to inoculate each substrate. The grains were mixed with gloved hands to be coated evenly with spores, then were transferred into damp wrung linen cloths. The cloths were folded closed and placed in perforated stainless-steel trays, which were then placed in an incubator fitted with heaters, misters, and temperature and humidity sensors. The incubator was set to 32°C and 70% relative humidity for the first 18 h. The following morning the grains were mixed with gloved hands to redistribute the growing mycelia and dissipate the heat produced by the fungus’ metabolism. Then the grains were returned to non-perforated stainless-steel trays, covered with freshly wrung linen cloths, and returned to the incubator at 28°C, lower this time to prevent overheating, for a further 24 h. By the end of this period, the grains had been bound together with fluffy white mycelia into a kind of cake, resulting in the formation of kōji.

### 2.2 Substrates and treatments

To develop our novel misos, we selected regional Nordic proteinous substrates including yellow peas (*Pisum sativum*, Unifood A/S, DK), Gotland lentils (*Lens culinaris*, Nordisk Råvara; a variety of lentil from the Swedish island of Gotland), fava beans (*Vicia faba*, Unifood A/S, DK), rye bread (Lagkagehuset bakery, DK), and soybeans (*Glycine max*, Unifood A/S, DK) as a control. The legumes were soaked in water overnight (2 parts water: 1 part dry legumes), then drained and simmered in fresh water until their texture was ‘al dente’—that they would yield when bitten but still be firm. The precise temperature and time required to achieve this texture varies depending on the specific quality of the legumes, the hardness of the water, pot and heat source, etc.; the only reliable way to achieve this texture consistently is therefore through the senses. Two treatments were used to prepare the three Nordic legume substrates: standard simmering as describe above and nixtamalisation. For the nixtamalisation process, the soaked legumes were cooked in a solution of 0.1% calcium hydroxide (CaOH). The pots were removed from the heat, covered with a lid, and allowed to sit overnight at room temperature for the alkaline transformations to proceed. The following day, the legumes were drained from the solution, rinsed with cold water, and drained again before being ground (Redzepi & Zilber, 2018). The rye bread was a standard rye bread purchased at Lagkagehuset, a chain of bakeries in Denmark. The kōjis, legumes and rye bread were all ground using a Varimixer Kodiak 20.

### 2.3 Development of novel misos

We developed eight misos using different proteinous substrates using Noma’s standard recipe (Redzepi & Zilber, 2018). Ground pearled barley kōji (or rice kōji for the control with soybeans) and cooked ground proteinous substrates (detailed above) were combined in a 2:3 parts ratio. 4% salt of the combined weight was then added and mixed thoroughly with gloved hands. Each of the eight mixtures was divided into three 2 kg batches, and each batch was packed into a five-litre sterilised glass jar (Utopia Biscotti jars, interior diameter 19 cm, height 24 cm). Each miso was covered on the surface of the mixture with a layer of cling film, weighed down with ceramic weights, and fermented for three months at a constant temperature of 28°C and ambient humidity.

### 2.4 Sampling

Each miso was sampled three times over the three-month fermentation period: at the start, one month in, and at the end. The start samples were taken from the unfermented paste before each mixture was divided into three batches. The final samples were taken by lifting away some of the miso surface with sterile utensils and sampling the middle of the miso paste below. The miso pastes were put into 5 ml Falcon tubes with sterile utensils and stored in the freezer at −20°C. A total of 56 samples were collected: (3 legumes x 2 treatments + 1 rye bread + 1 soybean/rice control) x 3 batches x 2 samplings + 8 start samplings. Of these, only 32 samples (the start and end samples) were sequenced, due to budget constraints.

The pH of the Noma misos was measured at the start and end of fermentation. 5 g of each miso was mixed with 10 ml of water, and the solution was measured with a pH meter (Thermo Fisher, MA, USA). pH measurements were converted to concentration of hydrogen ions in the solution, multiplied by 3 (the dilution factor) to calculate the concentration of hydrogen ions in the miso, corrected for the pH of the water, and converted back to logarithmic scale to yield the pH of the misos.

In addition, six novel misos developed by Inua, a former restaurant in Tokyo affiliated with Noma, were shipped to Copenhagen for analysis (**Table S1**). These samples lack biological replicates. Inua used the same recipe as Noma to produce their misos, except that they mostly used barley kōji spores (also from Bio’c, Japan), while we, like Noma, used rice kōji spores. The overview of the substrates used in each location are shown in **Fig. 1**.

**Fig. 1.**
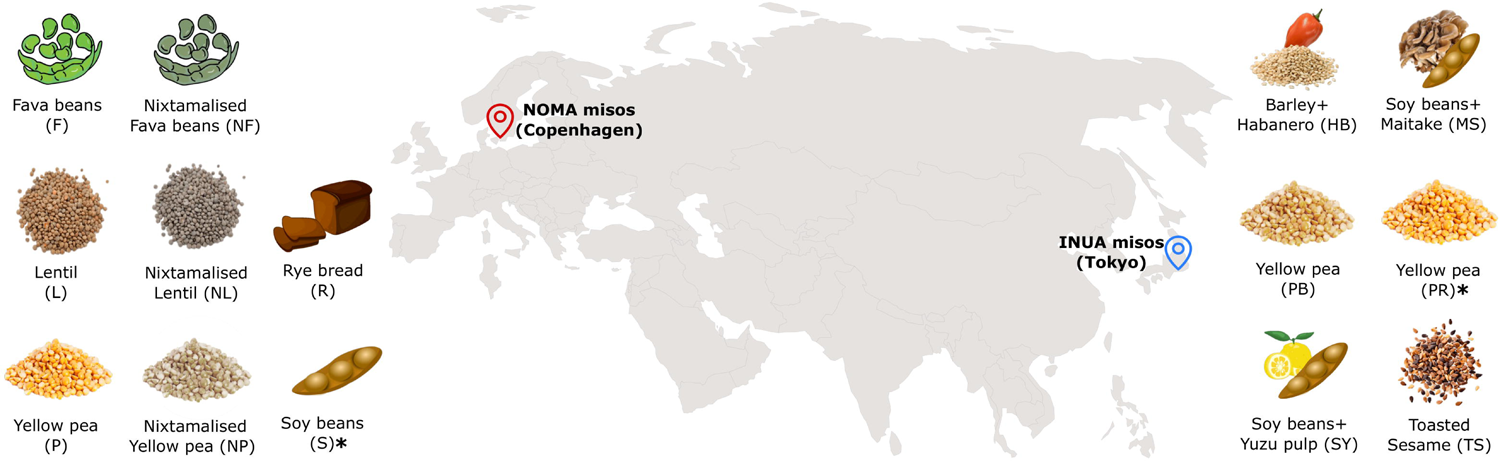
Schema of substrates used to produce misos and places of fermentation. All misos use barley as the kōji substrate, except those marked with an asterisk (*), which use rice.

Samples are named according to the following formula: [S][br].[s][tr], where [S] is the substrate (1- or 2-letter abbreviations), [br] is the biological replicate (numbered 1-3, only for end samples), [s] is the sampling time (1 for start, 3 for end), and [tr] is the technical replicate (lettered a-c).

### 2.5 Metabarcoding analyses

The DNA of the 32 samples (with technical triplicates, 96 samples total) from the miso experiment and the six from Inua (with technical triplicates, 18 samples total) was extracted using the Qiagen DNeasy PowerSoil Kit (∼0.2 g of sample). The DNA was amplified using the 515F and 806R primers that target the variable regions of 16S rRNA, a universal marker for bacterial identification (Apprill *et al*., 2015; Parada *et al*., 2016) and ITS3_KYO2 and ITS4 primers that target the nuclear ribosomal internal transcribed spacer (ITS) region, a common marker for fungal identification (Toju *et al*., 2012; White *et al.,* 1990). The PCR amplifications and subsequent library preparation were set up according to the Tagsteady protocol (Carøe & Bohmann, 2020), and the cycling conditions were 95°C for 5 min, followed by 40 cycles at 95°C for 20 s, 48°C (ITS) or 52°C (16S) for 30 s, and 72°C for 45 s. The final library was sequenced on an Illumina MiSeq instrument on a Version 2 flow cell in paired-end mode running 250 cycles, at the Globe Institute, University of Copenhagen, Denmark.

Raw reads were demultiplexed and barcodes and primers were trimmed using *cutadapt* version 2.9 (with parameters e=0, no-indels, m=100). Reads were then quality filtered using the *filterAndTrim* dada2 v.1.22 R function (with parameters maxEE =6, truncQ=2). The 16S reads were trimmed to 200 base pairs (both forward and reverse reads, truncLen=0 in filterAndTrim function). The ITS region has variable length, so we adapted the default procedure: ITS reads were not trimmed, and we checked that short ITS regions did not mistakenly contain primer reads. Amplicon sequence variants (ASVs) were inferred using the dada function and reads were merged using the *mergePairs* function (with parameter minOverlap=12). Chimeras were removed using the *removeBimeraDenovo* dada2 R function.

We assigned taxonomy for each ASV using the Bayesian RDP classifier, as implemented in dada2 (function assignTaxonomy, parameter minBoot set to 50) with the SILVA database (v.1.3.2) for 16S reads and the UNITE (v.8.2) database for the ITS reads. The ASVs assigned to chloroplasts, mitochondria or without phylum assignation were removed, as well as those present in only one sample or with fewer than 10 reads.

Alpha diversity and beta diversity analyses were performed in R v.3.6.1 using the phyloseq and ggplot2 packages (v.1.30.0) (McMurdie & Holmes, 2013; Poirier *et al*., 2018). For these analyses, we averaged the rarefied reads of the technical replicates. Clusterings were plotted with the Bray-Curtis distance and ward.D2 method and statistics were performed on alpha diversity using Kruskal-Wallis. The Principal Coordinates Analysis (PCoA) was plotted with the ward.D2 method and PERMANOVA tests (Anderson, 2001) were performed to identify microbial compositional variations across four factors: substrate type, substrate treatment (nixtamalisation), sampling time, and geographical location. For substrate type, we exclusively considered end samples from Noma. For substrate treatment, we included fava beans, Gotland lentils, and yellow peas from Noma, as well as their nixtamalised versions. For sampling time, our analysis included both start and end samples collected from Noma. Finally, for geographical location, we integrated end samples obtained from both Noma and Inua. DESeq2 was applied to characterise statistically significant differentially abundant ASVs in the different samples (Love *et al*., 2014).

### 2.6 Shotgun metagenomic analyses

For shotgun metagenomic analysis, we re-extracted DNA from each sample using 3 g of miso. The samples were diluted in 4.4X saline water (0.9% NaCl), placed in stomacher bags (BagPage, Interscience, France) and homogenised in a laboratory blender Stomacher 400 (Seward Co.) at high speed for 2 min. This mixture was filtered (filter porosity of 280 microns) and subsequently centrifuged to concentrate the cells. The pellet was then used for DNA extraction using the Qiagen DNeasy PowerSoil Kit. DNA from the 38 miso samples was sent to BGI Group (Hong Kong, China) for metagenomic sequencing using the DNBSEQ-G400 platform with PE150 chemistry, generating a dataset of at least 16M paired-end reads. Quality control and preprocessing of fastq files were performed with fastp v.0.23.2 (Chen *et al*., 2018).

#### 2.6.1 Microbial diversity

We first estimated microbial community structure by mapping the reads against the catalogue contained in the MetaPhlAn tool v.3.0.4 (Truong *et al*., 2015). Paired-end reads were assembled into contigs using MEGAHIT v.1.2.9, with default settings. We then predicted genes using Prodigal (v.2.6.3) and marker genes were extracted using fetchMG, v.1.0 (Ciccarelli *et al*., 2006; Sunagawa *et al*., 2013). Thereafter, taxonomic assignments were made using the marker genes *ychF* and *leuS*, whose closest homologue was assigned by a BLAST search on all available sequences in the NCBI protein database. Species composition plots were created in R (v.3.6.1) using the package ggplot2, v.3.3.2.

#### 2.6.2 Safety concerns

Since we identified *Staphylococcus* species in the samples, we investigated whether they possessed genes encoding enterotoxins. Using BLAST, we searched for the presence of pyrogenic superantigenic toxins (PTSAgs: *sea-see*, *seg-sevu*, *selv*, *selx*, *sey*, *selz*, *sel26*, *sel27* and *TSST1*) and exfoliative toxins (*eta*, *etb*, *etd*) in the assembled metagenomes. Reliable virulence genes were confirmed if sequence identity and query coverage were both >80% (Zhou *et al*., 2021).

#### 2.6.3 Metagenome-Assembled Genomes (MAGs)

Genome processing and refinement were performed using metaWRAP v.1.3.2, using binning module (-maxbin2-concoct-metabat2 options) and the resulting bins were refined with the bin_refinement module (-c 90 -x 10 options). The quality of the resulting prokaryotic bins was assessed with CheckM (Parks *et al*., 2015). For the eukaryotes, completeness and contamination of resulting bins were assessed with BUSCO (v.3.0.2; Simão *et al*., 2015), using the eukaryota_odb9 set of genes. Bins whose completion was less than 70% were discarded for further analysis.

Taxonomic classification of each MAG was performed using the Automatic Multi-Locus Species Tree web server (https://automlst.ziemertlab.com/; Alanjary *et al*., 2019), which determines closely related genomes based on the core gene alignments of the recovered MAGs. Closest species were inferred based on the percent average nucleotide identity (ANI) calculated using FastANI, v.1.31 (Jain *et al*., 2018). Finally, phylogenomic analyses were performed using Multiple Conserved Genomic Sequence Alignment with Rearrangements (MAUVE v.2.4.0) (Darling *et al*., 2004). Annotation and phylogenomic tree management were performed on iTOL v.5 (http://itol.embl.de).

#### 2.6.4 Phylogenetic and functional analyses

To assess the genomic relatedness between MAGs from different locations and substrates, we performed ANI analyses between recovered genomes of the same species. Next, for the species that showed considerable differences based on the ANI distances, we computed a phylogenetic tree using a concatenated matrix of the set of conserved proteins, representing essential functions shared across different genomes—in this case among our MAGs and other strains from the RefSeq database. Rapid Annotations using Subsystem Technology (RAST; Aziz *et al*., 2008), was used to annotate all MAGs, and protein-coding sequences were extracted for subsequent analysis. We calculated the core and pan-genomes for the 39 selected proteomes to identify a set of conserved orthologs using the Bacterial Pan Genome Analysis tool (BPGA; Chaudhari *et al*., 2016). We identified a set of 133 conserved protein sequences at a minimum sequence identity cutoff of 50%. The set of 133 orthologs was then collected from each genome, the amino acid sequences were aligned, and concatenated to generate a super matrix with 41,381 characters for phylogenetic analysis using IQtree 2 (Minh *et al*., 2020). A substitution model was calculated for each partition in the super matrix and then a phylogenetic tree was calculated. The tree showed excellent support for most nodes. The process was performed automatically using the script available at https://github.com/WeMakeMolecules/Core-to-Tree with the core_seq.txt and DATASET.xls files obtained as outputs from the BPGA run. Then, the unique proteins identified in each group were annotated using the BlastKOALA server (https://www.kegg.jp/blastkoala/).

### 2.7. Data availability

The raw sequences of 16S rRNA and ITS genes and metagenomic reads were deposited on the European Nucleotide Archive (ENA) under the BioProject ID PRJNA992639. The MAGs are available at doi: 10.17632/sjyf9kncs9.1.

## 3 Results

We first employed metabarcoding approaches to analyse the microbial composition and diversity of the novel misos. This analysis allowed us to identify the main genera present in the samples and to detect significant microbial differences across four factors: substrate type, substrate treatment (nixtamalisation), sampling time, and location. We then used shotgun metagenomic sequencing to obtain a more refined understanding of the microbiota in the samples. This approach provided better resolution of the taxonomy to species/strain level, and enabled us to identify the presence of enterotoxin genes and conduct functional analyses.

### 3.1 Revealing the microbiota of the novel misos using metabarcoding

We used the metabarcoding data to analyse the microbial composition and diversity of the novel misos.

#### 3.1.1 Microbial composition of the novel misos

The bacterial and fungal composition of the novel misos was first assessed using amplicon sequencing targeting the 16S rRNA and ITS genes, respectively. The sequences were clustered in 115 bacterial and 55 fungal ASVs, whose taxonomic assignment was possible down to the genus or species level for the majority of the ASVs (**Table S2**). In one replicate of the Gotland lentil miso (L1.3c, ITS gene, **Table S2**), the sequenced reads did not provide sufficient depth, leading to its exclusion from the analysis.

For the bacterial composition, we identified three main groups in the clustering of the misos (**Fig. 2A**). The first was mainly represented by *Pediococcus pentosaceus* and contains the nixtamalised and fava bean misos produced at Noma, as well as the habanero-barley (HB) miso from Inua. The second group was dominated by *Bacillus* spp. and species from the *Enterobacteriaceae* family, and contains samples from the beginning of fermentation and the rye bread misos. The last cluster was heterogeneous and can be divided into three subgroups. The first subgroup was rich in the genus *Staphylococcus*, represented mainly by misos made with soybeans and yellow peas. The second was dominated by *Staphylococcus* and some species of the *Lactobacillaceae* family, including the maitake-soy (MS) miso from Inua and some replicates of the yellow pea and Gotland lentil misos from Noma. The last subgroup comprised misos dominated by either *Enterococcus* spp. or *Pediococcus acidilactici*, represented only by misos from Inua.

**Fig. 2.**
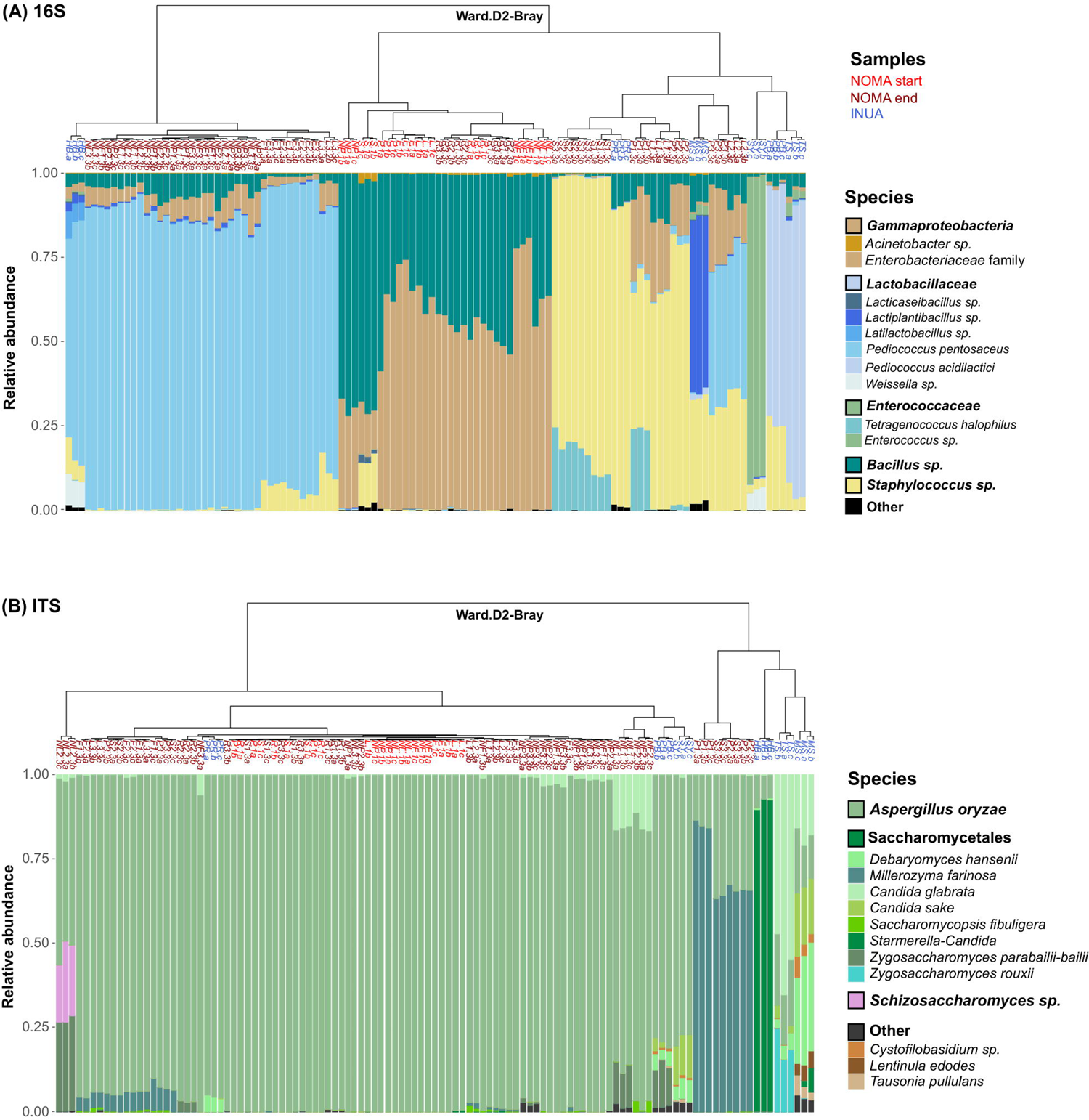
Plot depicting the relative abundance of the main species found in the misos using metabarcoding analyses based on ASV data of 16S rRNA **(A)** and ITS **(B)** gene regions. The samples were clustered using the Bray-Curtis distance matrix based on the ward.D2 method.

For the fungal composition, as expected, most samples were dominated by *A. oryzae* (**Fig. 2B**). However, some replicates of yellow pea and soybean misos made at Noma and some of the Inua misos—habanero-barley (HB), maitake-soy (MS), and toasted sesame (TS)—were co-dominated by other species of the Saccharomycetales family, mainly *Millerozyma farinosa*, *Starmella-Candida* spp.*, Debaryomyces hansenii* and *Candida glabarata*.

#### 3.1.2 Microbial diversity of the novel misos

Alpha and beta diversity analyses and differential abundance tests were performed to identify the main microbial differences among the novel misos between substrates, treatments (standard and nixtamalised), sampling times (start and end), and locations (Noma and Inua) (**Fig. 3**).

**Fig. 3.**
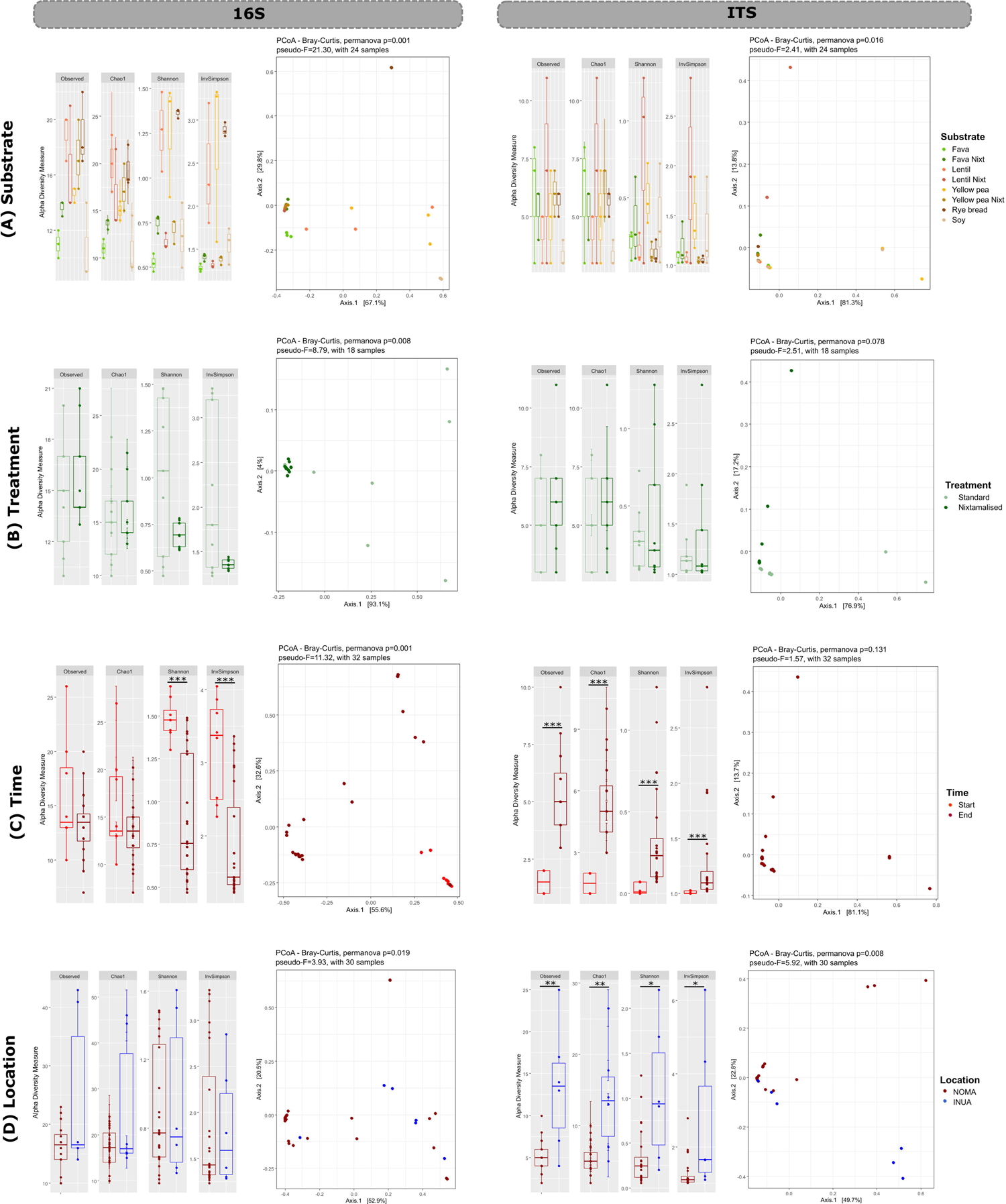
Boxplots of alpha diversity indices and PCoAs among bacterial (16S) and fungal (ITS) communities, analysed according to substrate **(A)**, treatment **(B)**, time **(C)** and location **(D)**. Significance in the alpha diversity analyses is indicated at multiple levels: * p<0.05, ** p<0.01, *** p<0.001. For the beta diversity (PCoAs), a larger pseudo-F value indicates a greater factor of difference in the comparison.

First, we analysed the microbial diversity of the eight substrates used in the Noma misos. Concerning alpha diversity, there was a significant difference between the substrates for bacterial composition (Kruskal-Wallis test, p<0.05 for all indices). Overall, fava bean and soybean misos showed lower ASV richness (Tukey test, Observed and Chao1 indices; **Fig. 3A**, **Table S3**). In the Shannon and InvSimpson indices, which consider species richness as well as evenness, we noticed a higher diversity in rye bread, Gotland lentil and yellow pea misos. The beta diversity analysis showed a strong and significant effect of substrate on the samples (PERMANOVA, pseudo-F=21.30, p=0.001). In the PCoA, some biological replicates of yellow pea and Gotland lentil were clustered nearer to soybean samples for the bacterial composition. Although these samples are abundant in *Staphylococcus spp.*, differential abundance analysis showed that the Gotland lentil and yellow pea samples were more likely to contain *Enterobacteriaceae*, *Bacillus* spp. and *Pediococcus pentosaceus*, when compared to the soybean control miso (DeSeq2, p_adj_<0.001). As mentioned above, regarding bacterial composition, rye bread samples were clustered far apart from the other samples because of the high abundance of *Enterobacteriaceae* and *Bacillus* species. The nixtamalised misos, meanwhile, were closely related, especially due to their abundance of *Pediococcus pentosaceus*. For the fungal composition, no significant differences were detected between the substrates for all indices in alpha diversity analyses (Kruskal-Wallis, p>0.05; **Fig. 3A**, **Table S3**). In fungal beta diversity, there was a small but significant effect of substrate (PERMANOVA, pseudo-F=2.41, p=0.016). Most samples were closely clustered, with *A. oryzae* being dominant. However, some nixtamalised Gotland lentil miso replicates also presented co-dominance of the species *Zygosaccharomyces parabailii-bailii*, *Schizosaccharomyces spp*. and *Candida glabarata* (**Fig. 2B**). Another cluster observed in the fungal PCoA consists of replicates of soybean and yellow pea misos, whose microbiota are richer in *M. farinosa* when compared to the other substrates (DeSeq2, p_adj_<0.001).

We also measured the microbial diversity of the final misos comparing the two treatments—standard and nixtamalisation—used in the production of yellow pea, fava bean and Gotland lentil misos at Noma. In the alpha diversity analysis for bacterial composition, we noticed no significant difference in the ASV richness between these two groups (Kruskal-Wallis test, p>0.05 in all indices; **Fig. 3B**). Beta diversity analysis distinguished the groups as statistically different in bacterial composition (PERMANOVA, pseudo-F=8.79, p=0.008), and differential analyses showed a higher abundance of *Staphylococcus* spp. and *Enterobacteriaceae* in the standard misos. Regarding ITS, no significant differences were observed between standard and nixtamalised misos in the alpha (Kruskal-Wallis test, p>0.05) and beta diversity analyses (PERMANOVA, p>0.05).

We then compared the microbial diversity at the start and end of the fermentation among the Noma samples. Based on the Observed and Chao1 indices, there was no significant difference in bacterial composition between these two groups (Kruskal-Wallis test, p>0.05). Based on the Shannon and InvSimpson indices, however, we noticed a higher richness and evenness of ASVs in the starting samples (Kruskal-Wallis test, p<0.001; **Fig 3C**). In the beta diversity analysis, we observed that the composition and abundance of bacterial ASVs between these two groups have significant differences (PERMANOVA, pseudo-F=11.32, p=0.001). The misos before fermentation are more abundant in *Bacillus* spp. and *Enterobacteriaceae*, in contrast to the fermented samples, which are richer in *Staphylococcus* spp.*, Pediococcus pentosaceus* and *Tetragenococcus halophilus* (Deseq2, p_adj_<0.001; **Table S3**). For fungal composition, we noticed a significant difference in all indices of alpha diversity between start and end Noma samples, where the samples before fermentation have lower richness and evenness of ASVs (p<0.001, **Fig. 3C**). However, the beta diversity for fungal composition showed no differences between the two groups (PERMANOVA, p>0.05).

Finally, we compared the finished misos between Noma and Inua. The alpha diversity analysis based on all indices showed similar observed richness and evenness of bacterial ASVs for Inua misos compared to the Noma fermented samples (p>0.05, **Fig. 3D**). However, the beta diversity of bacterial composition showed differences between the samples of these two restaurants (PERMANOVA, pseudo-F=3.3, p=0.019). These differences were highlighted by differential analyses, where the Inua misos presented greater abundance mainly of *Weissella, Enterococcus sp*. and *Pediococcus acidilactici*, while the Noma samples were richer in *T. halophilus* and *P. pentosaceus* (Deseq2, p_adj_<0.001). Concerning fungal diversity, Inua and Noma presented statistical differences in all alpha diversity indices, with Inua misos being more diverse in richness and evenness (Kruskal-Wallis test, p<0.05). For beta diversity, we also observed significant differences between these two groups (PERMANOVA, pseudo-F=5.92, p=0.008) with the effect for fungi slightly higher than that for bacteria, highlighted by the higher abundance of *M. farinosa* in the Noma misos, and *Candida sake* and *D. hansenii* in the Inua misos.

### 3.2. Genomic resolution of the microbiota using shotgun metagenomics

To confirm the community composition at higher genetic resolution, refine the species assignment, scan for potential enterotoxin genes related to food safety, and perform strain-level and functional analyses, we used shotgun metagenomics.

#### 3.2.1 Community composition and species assignment

Shotgun metagenomics was performed to a depth that would allow the assembly of complete genomes to enable more detailed analyses (**Table S4**, ‘reads_quality’ sheet), such as taxonomic resolution to species and/or strain level. In the previous metabarcoding analysis, some substrates showed variation among the experimental replicates (**Fig. 2**). This within-substrate variation could be due to batch variation and/or because the small sampling weight selected for DNA extraction (∼0.2g) may have captured different parts of the community. Thus, for the shotgun metagenomic analyses, we repeated the DNA extraction, using a protocol involving a larger sample size for each substrate (∼3g; see section 2.6).

To start, we mapped the metagenomic reads against a genomic database (MetaPhlAn tool) and identified 78 bacterial and two fungal species (**Table S4**, ‘metaphlan’ sheet). Overall, biological replicates presented a consistent microbial composition within each substrate (**Fig. 4A**). This finding clears up the question of variability between the batches, which in the metabarcoding could have been due to smaller and therefore less representative sampling. Furthermore, the bacterial species composition obtained from this mapping was similar to the 16S amplicon analysis and resolved most of the ambiguities of the taxonomic assignment of species, especially those of the genus *Staphylococcus*. *S. epidermidis* and *S. pasteuri* were dominant especially in the Noma misos with fava beans, Gotland lentils, yellow peas and soybeans. *S. warneri* was identified in all the misos from Inua. Concerning fungi, only *A. oryzae* and *S. cerevisae* were identified, while the ITS amplicon analysis detected a wider variety of fungal species, likely indicating a lack of available reference genomes in the MetaPhlAn database.

**Fig. 4.**
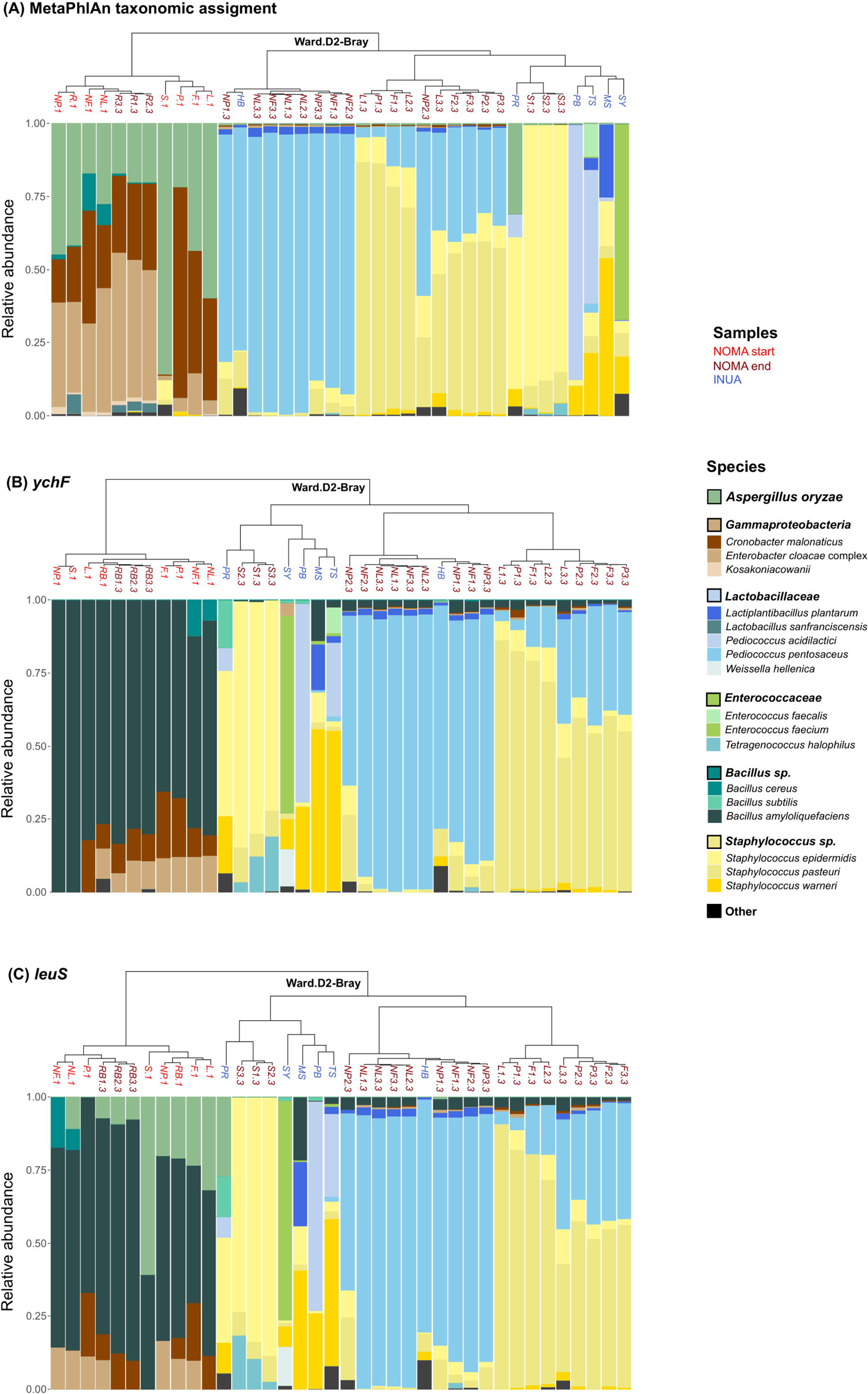
The relative abundance of microbial communities in the misos using shotgun metagenomic data. Values were calculated from the reads using the MetaPhlAn database **(A)** and from the coverage of the *ychF* **(B)** and *leuS* **(C)** marker genes assembled from the metagenomes. The samples were clustered using the Bray-Curtis distance matrix based on ward.D2 method.

To overcome this limitation by further characterising the microbiota independently of the set references, we applied an analysis using two marker genes—*ychF* and *leuS*—from the assembled metagenomes. The marker gene *ychF* yielded 30 bacterial and four fungal species, and the *leuS* gene yielded 23 bacterial and five fungal species (**Table S4**, ‘ychF’ and ‘leuS’ sheets). Although the *ychF* gene detected some fungal species such as *D. hansenii, Z. bailii, C. glabarata* and *S. cerevisae*, it was unable to detect *A. oryzae,* the main kōji mold present in miso. Therefore, we chose another marker gene (*leuS*), which detected *A. oryzae*, as well other fungal species such as *Cyberlindnera fabianii* and *Millerozyma farinosa*. Overall, the marker-gene analyses were consistent with the previous analysis of mapping to the MetaPhlAn database (**Fig. 4**), but some differences—besides the presence of other fungal species—are important to highlight. In particular, we note the detection of *Bacillus amyloliquefaciens*, a species probably missing in the MetaPhlAn database, which is dominant in the start samples as well as in the rye bread miso.

#### 3.2.2 Food safety concerns

Although the dominant *Staphylococcus* species identified in the misos were coagulase-negative, they may carry enterotoxin genes. In the search for pyrogenic toxin superantigens and exfoliating toxins in the metagenomes, we detected *selx* and *sel26* genes in the habanero-barley miso (HB) and in one sample of the Gotland lentil miso (L3.3). The coverage of these genes was the same as that for *S. aureus*, which was detected at a relative abundance of around 2.6-3.2% in both samples (**Table S4**, ‘ychF’ and ‘leuS’ sheets).

#### 3.2.3 Strain-level and functional analyses

From the 38 metagenomic samples, 90 prokaryotic and 10 *A. oryzae* metagenome-assembled genomes (MAGs) of high quality were recovered (**Table S5**). The majority of the bacterial MAGs obtained corresponded to *B. amyloliquefaciens,* followed by the species *P. pentosaceus, L. plantarum, S. pasteuri* and *S. epidermidis* (**Fig. 5A**). Additional comparative analyses were conducted of MAGs within the same species (**Table S6**), which aimed to assess the degree of relatedness or divergence among strains emerging in distinct substrates and geographical locations. Most of the strains within each species group exhibited high similarity, except for those of *S. epidermidis*. ANI and core phylogenetic analyses highlighted that the MAGs from this species were divided into two distinct subclusters (**Fig. 5B**), corresponding to the substrate used. The first group included one strain recovered from the maitake-soy (MS) miso at Inua, and the second cluster was composed of three strains from the yellow pea misos made at each location. This ‘yellow pea group’ shared ANI values ranging from 98.6 to 99.4% within its group and from 96.5% to 97.3% with the *S. epidermidis* strain recovered from the MS miso (**Table S6)**. In a pangenomic analysis performed on these two groups of strains, 1,726 of the core genes were shared between the yellow pea group and maitake-soy MAGs, with 63 and 445 genes unique to each group, respectively. Among the annotated genes, the unique genes in the yellow pea group were mainly associated with biofilm formation (*icaA, icaB, icaD, icaR*).

**Fig. 5.**
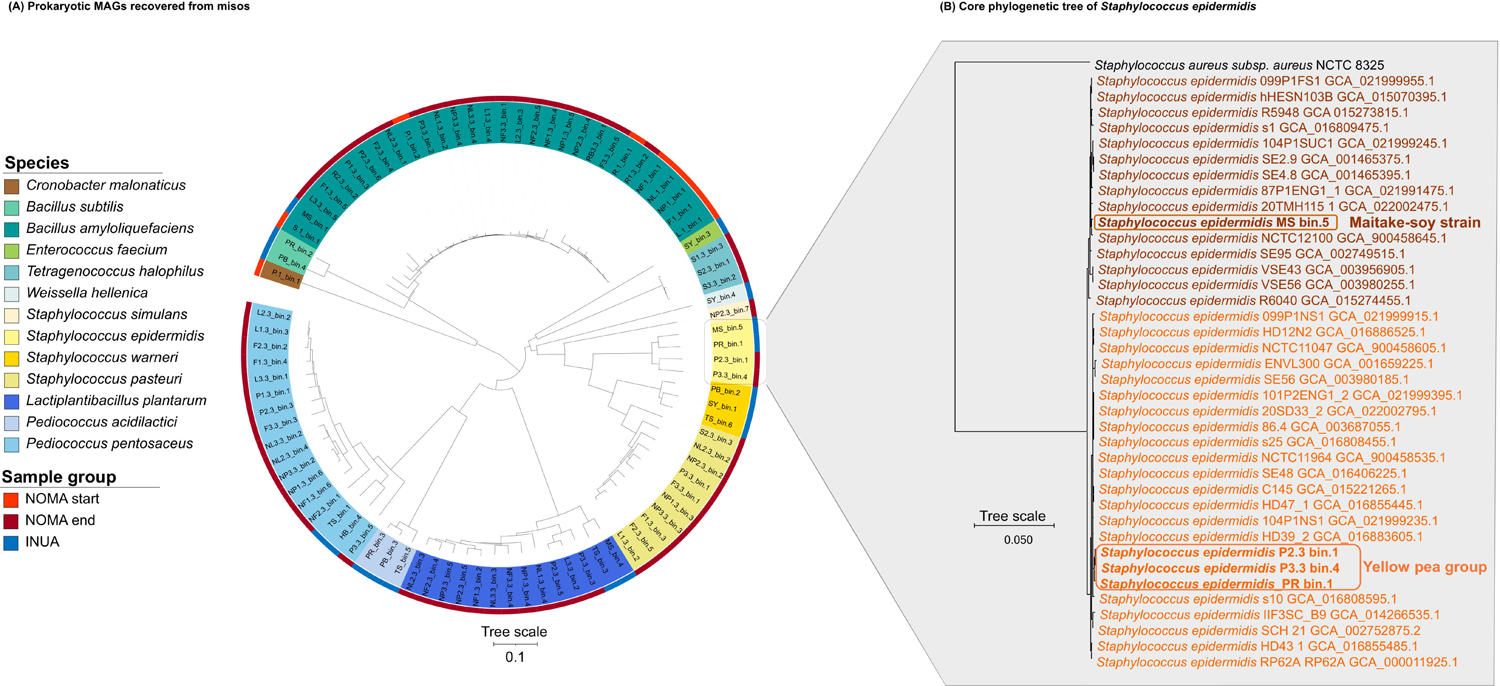
Phylogenomic tree of the MAGs recovered from the 38 samples **(A)** and core phylogenetic tree of *S. epidermidis* MAGs **(B)**.

For the MS strain, there were many unique genes, mainly associated with amino acid and energy metabolism, cofactors and vitamins, enzymes, and carotenoid biosynthesis, as well as several genes poorly characterised (**Table S7**). Several unique genes belonged to the category “Genetic information processing”, such as genes related to DNA replication and repair, translation, transcription and regulation. We also identified genes related to the prokaryotic defence system, such as CRISPR-associated proteins, as well as ABC transporters and two-component systems, important mechanisms that enable bacteria to respond to environmental changes and optimize their growth and survival.

We focused our investigation on the carotenoid biosynthesis genes found in the MS *S. epidermidis* strain (*crtM, crtN, crtO, crtP* and *crtQ*), due to the various functions, many especially food-relevant, that carotenoids play in bacteria, including pigmentation and essential biological processes such as antioxidant activity. Hypothesising horizontal gene transfer (HGT), we investigated where the MS strain had acquired these genes from. We first studied the maitake, as the main difference between the MS miso and the yellow pea group, searching for carotenoid biosynthesis genes in maitake strains available in the public database NCBI (https://www.ncbi.nlm.nih.gov/genome; *Grifola frondosa* 9006-1 and MG88), but found none. We then examined the MAGs of *S. warneri* recovered from the misos in this study, as HGT is both more common among bacteria, more likely between species of the same genus, and, among the other *Staphylococcus* spp. present in the MS miso (*S. warneri* and *S. pasteuri*), *S. warneri* is much more abundant. While no MAGs for *S. warneri* were recovered from the MS miso (instead from PB, SY, and TS), it is not unlikely that the *S. warneri* in the MS miso is the same or a similar strain. Analysis of these *S. warneri* MAGs revealed two genes linked to carotenoid biosynthesis: *crtM* and *crtN*. Noting the relatively high abundance of *S. warneri* in the yellow pea miso produced at Inua (PR) compared with the yellow pea misos produced at Noma (P2.3, P3.3), we conducted a more detailed analysis of the *S. epidermidis* MAG from this miso as well, in which we also identified carotenoid genes identical to those in the MS miso (*crtM, crtN, crtO, crtP* and *crtQ*).

## 4 Discussion

There is limited research characterising miso ecology, making it difficult to compare these novel misos to traditional ones, as even the traditional ones have yet to be sufficiently characterised. Based on the partial list of culturable miso taxa compiled by Allwood *et al*. (2021), the Noma and Inua misos share many of the microorganisms commonly found in miso, as well as some that are unique to each sample. The distinctive features of these misos lies in their overall microbial composition, which can include differences at the species and strain level. In this study as well as in our previous exploratory analysis (Kothe *et al*., 2023), we identified *A. oryzae* and *B. amyloliquefaciens* in almost all the miso samples analysed. The first is the main species in the kōji starter culture; the second may also be present in this starter culture (Woldemariam *et al*., 2020). Both our studies reveal that the microbial ecology of miso is shaped by its proteinous substrate. While our first study lacked replicates and thus offered this finding tentatively, the present study can offer it more certainty. Furthermore, the novel misos we have investigated present a higher microbial diversity than currently described in the miso literature (Allwood *et al*., 2021). These findings suggest that there are probably many microorganisms involved in miso production, including ones not previously found in misos as well as ones new to science, which may contribute to their flavours, texture, and nutrients. For instance, we previously discovered a potential new species of *Exiguobacterium*, a genus for which no species had ever been found in miso, and with only one other documented instance of being found in a fermented food (Kothe *et al*., 2023).

In the subsequent sections, we discuss the main dimensions of difference among the misos—the substrates used, nixtamalisation effect, time, and place of fermentation—and provide an overview of the potential roles of the microorganisms identified in the misos. We then explore gene-level insights such as considerations of food safety and the adaptability of strains to different substrates.

### 4.1 Dimensions of Difference

#### 4.1.1 Substrate effect

In our study, we observe that substrates significantly impact the structure of miso microbiota. The novel misos made at Noma vary in microbial composition and diversity depending on the substrate, with some substrates showing clearer differences than others, particularly in bacterial composition (indicated by the high pseudo-F and R^2^ values; **Fig. 3A**, **Table S3**). This substrate effect may have arisen from the autochthonous microbiota associated with the proteinous ingredients, which can exhibit distinct behaviour during fermentation. An additional, non-mutually-exclusive explanation is that since different substrates consist of varying nutrient and energy sources, they represent distinct microbial niches that differentially favour or inhibit the growth of different microbial species.

Excluding the non-fermented start samples, we identified three main clusters of the misos based on their microbial composition (**Fig. 4**). The first group, consisting of the nixtamalised samples, is dominated by *Pediococcus pentosaceus,* a species commonly found in plant-based fermentations (Qi *et al*., 2021), as well as in raw substrates like beans and cereals (Ucak *et al*., 2022; Verni *et al*., 2017). In the second cluster are the standard misos of Gotland lentil, yellow pea and fava bean, and we identified co-dominance of the species *P. pentosaceus* and *S. pasteuri*. Although *S. pasteuri* is less reported in food compared to the other species detected in the misos, it has been documented in *douchi*, a Chinese product of fermented black soybeans (Li *et al*., 2018). The third group consists of the soybean control samples, where *S. epidermidis* was detected as the predominant species. This species is also not frequently reported in food and is more commonly associated with the human skin microbiome (Brown & Horswill, 2020). While we also detected this species in misos from our previous study (Kothe *et al.,* 2023), other *Staphylococcus* species are described in the miso literature, such as *S. gallinarum* and *S. kloosii* (Allwood *et al*., 2021). The *S. Epidermidis* in our study may have originated from the producers’ hands and adapted to the soybean and/or rice kōji substrate used in this traditional miso.

The rye bread samples clustered with the non-fermented start samples. At first, we thought this meant that the rye bread misos did not ferment; however, pH measurements suggest microbial activity, as the pH dropped in all three biological replicates (start pH: 4.52; end pH 3.99±0.23). Based on these data, we suggest the rye bread misos did ferment but with the microorganisms present initially, rather than the community shifting over time.

#### 4.1.2 Nixtamalisation effect

We also observed a clear effect of nixtamalisation on the miso microbial communities: the nixtamalised misos were significantly more similar in ASV composition and diversity to each other than to their non-nixtamalised counterparts, specifically for bacteria. The nixtamalised samples exhibit a high abundance of *P. pentosaceus* and *L. plantarum*, while the standard misos are predominantly composed of *S. pasteuri*. A study of the microbiota of fermented nixtamalised corn also identified *Pediococcus* and *Lactobacillus* (Sefa-Dedeh *et al*., 2004), suggesting that the alkaline environment created through nixtamalisation favours the development of these genera. This hypothesis is also supported by our previous study (Kothe *et al*., 2023), in which we also detected *P. penstosaceus* in a nixatamlised yellow pea miso, different from the samples used in this study but made with the same recipe.

The chemistry of nixtamalisation suggests a potential mechanism for this hypothesis. The alkaline treatment employed during nixtamalisation converts the hemicellulose components of the cell walls of grains and legumes into soluble gums, which improves the release and accessibility of nutrients such as vitamins, minerals and other bioactive compounds (Kamau *et al*., 2020). The resulting simpler carbon sources and the compounds created by the treatment can promote the growth of *Pediococcus*, a bacterium known for its ability to thrive at various pH levels, including alkaline conditions (Dey *et al*., 2019; Nakagawa & Kitahara, 1959). In contrast, the standard misos show a higher abundance of *Staphylococcus*. This may be attributed to the fact that species of this genus can secrete enzymes to break down complex sugars and starches, thus able to utilise a wider range of carbon sources than other taxa (Lakshmi *et al*., 2013), which may help it outcompete species such as *Pediococcus* in the miso environment.

It is worth noting that *Pediococcus* and *Lactobacillus* are commonly autochthonous to legumes (Sáez *et al*., 2017), and are frequently employed as starter cultures in plant-based fermentation to develop desired flavours in the final products (Ferri *et al*., 2016). These lactic acid bacteria play an important role in improving resistance to pathogens, since they lower the pH by producing organic acids. Their presence in these fermented plant-based products is consistent with these general patterns.

#### 4.1.3 Time effect

Perhaps not surprisingly, but still important to note, is that we observed an effect of the fermentation process itself on the microbial communities of the misos, specifically for bacteria. We identified that our start samples, before fermentation, have a higher abundance of *Enterobacteriaceae*, which are typically associated with gut microbiomes. After fermentation, the microbiota shifted to primarily consist of lactic acid bacteria and the *Staphylocococaceae* family, with variations based on the specific substrate used. Other studies focusing on fermented plant-based substrates, such as coffee and sourdough, have also consistently demonstrated a similar finding, that these raw materials, prior to fermentation, have a higher abundance of *Enterobacteriaceae*, and that the fermentation process produces a more selective microbiota (Costa *et al*., 2022; Pothakos *et al*., 2020).

Meanwhile, we observed the opposite tendency with the fungi, with diversity increasing by the end of the fermentation, as clearly shown by the alpha diversity graphs. This notable pattern of opposing trajectories for bacterial and fungal diversity over the course of the fermentation can be explained by the process by which the misos were produced. At the start of the fermentation, the miso mixture was fungally dominated by the *A. oryzae* inoculated from the kōji, while bacterially exhibited a range of taxa potentially introduced from the ingredients, equipment, fermenters’ bodies and surrounding environment. As fermentation progressed, the growth of lactic acid bacteria made fewer bacterial taxa dominant, while the dropping pH, combined with the salt, selected against *A. oryzae* and allowed other salt- and acid-tolerant yeasts to grow. This increase in non-*Aspergillus* fungi would have lowered the relative abundance of *A. oryzae* DNA in the fermented samples even if the amount of *A. oryzae* DNA remained constant, and especially if some of it became degraded over the fermentation which is likely. Nonetheless, by the end of the fermentation, the *A. oryzae* DNA remained dominant, explaining the lack of significant effect of time in fungal beta-diversity.

Along with the *A. oryzae*, we also observed a higher abundance of *B. amyloliquefaciens* in all the start samples. This finding suggests that the kōji spores used in miso production may harbour other species with functional relevance in addition to the traditional filamentous fungus *A. oryzae*. Overall, the fermented samples exhibit a diverse array of identified species which also differs from their starting microbiota, emphasising the significant impact of fermentation on the composition of the final miso microbiota.

#### 4.1.4 Geographical effect

The effect of fermentation location on the miso microbial communities is not fully clear and seems to differ across genus, species, strain, and gene levels. It must be noted that this lack of clarity may be due to the different levels of robustness between the Noma and Inua miso datasets. While all misos were made with the same basic recipe, the Noma recipes were made with biological replicates, while the Inua ones were not. Thus, while there is notable variation in microbial composition among the Inua misos, it is difficult to determine whether this variation is related to substrate or just batch variation, as there are no batch replicates within substrate type. Overall, the samples collected at Inua showed a higher fungal species diversity than those obtained at Noma (**Fig. 3D**). This difference in fungal diversity could be attributed to variations in handling environments.

In our previous study of novel misos made at Noma (Kothe *et al*., 2023), we also observed a higher level of species diversity compared with the misos made at Noma in the present study. This difference could be attributed to the different circumstances under which each batch of misos was made. The misos from the first study and those in this study were made in two different locations, at different times, under the creative direction of two different people, at different moments in the restaurant’s stylistic evolution. In the first study, the misos were made in 2017, in Noma’s original fermentation lab which at the time was housed in a few shipping containers behind the restaurant, more open to the environment. This was under the direction of a chef with a more tinkering approach, at a moment in the restaurant’s history when slightly more variability was tolerable. The misos in the current study, meanwhile, were made in 2018, after the restaurant had moved to a new location in a brand-new building, in a purpose-built fermentation lab with a more controlled environment. This was under the direction of a different chef, with a more controlled, scientific approach, and at a moment in the restaurant when consistency was becoming ever more important. It is likely that these different micro-geographic circumstances and approaches could have shaped levels of diversity in the misos accordingly. A similar difference may have been at play between Inua’s higher diversity than Noma 2.0’s.

Another factor that could contribute to the variation observed between the Inua and Noma samples is the type of spores used, as Inua primarily used barley kōji spores while Noma mainly used rice kōji spores (**Table S1**). The different strains present in each starter culture may have influenced the microbial ecologies of the misos differently (we reached out to the producer Bio’c to ask about the microbiological differences between the different starter cultures, but they were not able to share this information). Additionally, while broadly similar, misos made with the same substrate in the two locations were not exactly identical because of different sourcing which could contribute to distinct microbial compositions, as could the additional seasonings like yuzu or habanero in some of the Inua misos.

At the genus level of bacterial composition, it appears that factors such as substrate, nixtamalisation and time have more impact on sample composition than geographical location, as indicated by the higher Pseudo-F and R^2^ values for these factors (**Fig. 3D**). The geographical effect for fungal composition is slightly higher than for bacteria, and higher than the substrate effect for fungi (not surprising given the dominance of *A. oryzae* in the misos). This observation of a present but relatively weak geographical effect is further supported by the clustering of some of the samples produced in Tokyo alongside some from Copenhagen. For example, the Inua yellow pea sample (PR) closely resemble the soybean miso samples from Noma (**Fig. 4**), both made using rice kōji and dominant in *S. epidermidis*. This microbiological similarity supports the chefs’ culinary conclusion that yellow peas serve as an effective alternative to the traditional soybean for miso, indeed regardless of the production location.

This finding aligns with a large-scale study conducted by Wolfe *et al*. (2014) on the microbial ecology of cheese rinds that reaches similar conclusions about the concept of ‘microbial terroir’. The study found that, contrary to a previous hypothesis that different geographic locations might have unique local microbiomes shaping fermentations in distinct ways (Felder *et al*., 2012), reproducible community types developed mainly by the treatment they received, regardless of the specific geographical location of production. They also acknowledge that a significant portion of the diversity observed may exist at the strain level, since they only analysed their samples to the genus level. Our study with miso, meanwhile, suggests a similar pattern of treatment over geography might extend to the species and strain level.

### 4.2 Gene-level insights

#### 4.2.1 Food safety

The presence of *Staphylococcus* species in food production often raises concerns regarding food safety. Our study did not detect enterotoxin genes in the samples containing coagulase-negative *Staphylococcus* species. However, in the habanero-soy (HB) miso from Inua and one of the Gotland lentil misos (L3.3) made at Noma, we identified the presence of *S. aureus*, a pathogenic species known for its toxin production (Otto, 2014). The relative abundance of *S. aureus* bases on metagenomic read coverage was 2.6% in HB and 2.8% in L3.3, indicating a small but significant proportion of this species (This species is absent in the figures because of its low abundance, which falls below the inclusion threshold of 3%; see instead **Table S4**). In these samples, we detected genes encoding two enterotoxins (*selx* and *sel26*).

Although the presence of these genes is a valuable initial step to assessing potential risk, it is important to note it does not necessarily mean that the misos contain active toxins. Enterotoxin expression is favoured by environmental conditions such as optimal temperature growth for *S. aureus* (37°C) and neutral pH (between 6-7.5). Misos are typically fermented at a lower temperature (often room temperature or cooler in traditional production; these misos were kept at 28°C to expedite the process), and they typically begin at a pH lower than this range, often starting around pH 6 and dropping to around pH 5 (Allwood *et al*., 2023). Our misos exhibited a lower pH range, starting between pH 4.52 and pH 5.64 (mean=5.32, SD=0.38) and ending between pH 3.32 and pH 5.02 (mean=4.02, SD=0.45; **Table S1**). All but three ended at a pH below 4.6, the general threshold for safety recognised by the FDA (2007). Even the highest was at the lower range of typical misos (Allwood *et al*., 2023). While salt is less of a deterrent for production of these toxins (Schelin *et al*., 2011), which could still be produced at the 5% concentration of these misos, their low pH makes it unlikely.

Microbial guidelines for food indicate an acceptable level of *S. aureus* in ready-to-eat foods of less than 10^3^ CFU/g of food (Centre for Food Safety, 2014). Here we did not use culture dependent analysis, and consequently did not measure the CFU/g of *S. aureus* in these two misos. According to Rezac *et al*. (2018), aerobic bacteria counts of miso range from 10^2^ to 10^7^ CFU/g. If we consider that these misos had more than 10^6^ CFU/g in total, only 1% of *S. aureus*, 10^4^ CFU/g, would already be outside the permissible limit.

We emphasise that all misos produced at Noma and Inua for service, as with all their fermented products, are made according to carefully prepared and followed Hazard Analysis and Critical Control Point (HACCP) plans, and that these misos were experimental ones using the same recipe but outside of their typical production. So we do not believe there are any concerns about the safety of their misos used for service, or that this minor finding in our experiment should raise them. Indeed, in a previous study conducted by our team of Noma’s misos actually produced for service, we found no issues with *S. aureus* or any enterotoxins (Kothe *et al*., 2023). It should also be noted that only two of the 38 samples exhibited *S. aureus* or these enterotoxin genes, and that only one of the three Gotland lentil misos, all made at the same time with the same starting materials and protocol, exhibited these. We therefore conclude that this finding is most likely to be an exceptional contingency rather than symptomatic of any systemic problem.

Nonetheless, this result from our experiment emphasises that best practices for production and handling of miso should still be implemented by any producers, including conscientious manufacturing practices, proper sanitation procedures, regular monitoring of microbial contamination, and HACCP systems. Analyses for the presence of *S. aureus* could also be incorporated into food safety standards and guidelines for miso production to ensure the quality and safety of miso products and protect consumer health.

#### 4.2.2 Strain differentiation according to substrate

To bring our inquiry deeper to the strain level and investigate whether specific strains were associated with each substrate or production site, we conducted an analysis of the high-quality MAGs. We noticed that the substrate may have a greater impact on strain genetic diversity than the production site. Strains of *S. epidermidi*s identified in misos made at the different production sites but with the same substrate (yellow pea) were closely related, while another strain of *S. epidermidis* identified in the maitake-soy (MS) miso at Inua showed greater genetic differences compared with those found in the yellow pea misos at both sites (**Fig. 5B**).

Other studies have demonstrated that strains can adapt genetically when exposed to selective pressure from their environment. This adaptation can occur in response to various factors, such as the composition of nutrients, iron restriction, and/or high salt concentration, as in the case of cheese rinds (Bodinaku *et al*., 2019; Monnet *et al*., 2010). These findings support the possibility that the observed genetic differences among *S. epidermidis* strains associated with different substrates may be a consequence of the specific selective pressures exerted by these substrates.

We focused on elucidating the differences among the *S. epidermidis* strains from the yellow pea group and the maitake-soy miso (**Fig 5B**). Our analysis revealed the presence of unique genes in the MS strain, particularly those related to transcription, translation, and gene regulation processes, as well as genes associated with defence systems allowing prokaryotes to survive and adapt to different environmental conditions through the acquisition of new genetic material. The presence of these genes in the MS strain of *S. epidermidis* further supports the possibility of adaptative gene acquisition in this specific strain.

We also identified unique genes associated with carotenoid biosynthesis in the *S. epidermidis* MS strain compared to the ‘core sequences’ of the yellow pea group. Carotenoid biosynthesis genes are known to be widespread in various taxonomic groups of fungi (Sandmann, 2022) and bacteria (Ram *et al*., 2020). There are at least three hypotheses to explain how these genes appear in the MS strain and not in the yellow pea group.

The first hypothesis suggests the occurrence of horizontal gene transfer (HGT) from the maitake (*Grifola frondosa*) to the MS *S. epidermidis* strain. While our investigation for carotenoid biosynthesis genes in the maitake genomes available on NCBI yielded no positive results, these genes could be present in other maitake strains. Further studies could investigate this possibility by including other maitake strains, particularly those used in the production of the MS miso.

The second hypothesis proposes HGT between bacterial species in the misos. Considering that *S. warneri* carries carotenoid biosynthesis genes and is highly abundant in the Japanese misos, the transfer of genes from this species to *S. epidermidis* is plausible. This proposal aligns with well-established mechanisms of gene exchange observed between bacterial species in fermented foods (Bonham *et al*., 2017; Wang *et al*., 2023), most prevalent among closely related species (Bonham *et al*., 2017; Ravenhall *et al*., 2015).

The third hypothesis posits the pre-existence of the carotenoid genes in some *S. epidermidis* strains. Although *S. epidermidis* are not normally pigmented, Ogo (1985) noted the presence of a carotenoid glucoside similar to staphyloxanthin in some strains of this species. This finding suggests that the five genes coding for enzymes involved in carotenoid biosynthesis which we identified in *S. epidermidis* from the PR and MS misos could have existed in the strains already, rather than being taken up by them during the fermentation. It may not be a coincidence that our samples and Ogo’s study are from Japan. Though Ogo (1985) does not specify the strain used or its origin, as the study was conducted in Japan and published in Japanese we might reasonably infer (though cannot affirm) that the strain in question was also found in Japan. If so, our finding of these carotenoid genes, paired with Ogo’s identification of the carotenoid glucoside, suggests the possible emergence of carotenoid biosynthesis genes in geographically distinct populations of *S. epidermidis* strains, which would support, at the level of individual genes, a moderate effect of ‘microbial terroir’.

Based on the current data, the second or third hypotheses appear more likely than the first. Further studies could investigate the specific mechanism for the observed strain differentiation at the genetic level. Whether through pre-existing geographically distinct strain populations, adaptive horizontal gene transfer among bacterial species in the misos, cross-kingdom uptake from maitake, or a combination, this genetic analysis shows a clear effect of strain differentiation correlated with substrate, and, at the genetic level, a possible geographical effect.

## 5 Conclusion

This study provides valuable insights into the microbial ecology of miso, particularly novel misos, enhancing our understanding of this fermented product and its potential to be made using novel substrates. Our findings confirm the presence of important microorganisms, such as *A. oryzae* and *B. amyloliquefaciens*, in almost all miso samples, underscoring their significance as part of the kōji starter culture. We also demonstrate that the microbial diversity of miso extends beyond the taxonomic range currently described. And while all substrates showed successful fermentations, the microbiota of some substrates, like the yellow pea, exhibited marked similarities to that of the traditional soybean miso. This finding suggests that, at least from a microbial perspective, yellow pea is a particularly good alternative for soybean— confirming microbiologically what our chef collaborators had concluded through their culinary experimentation and sensory knowledge.

It is difficult to make any conclusive claims about the diversity of these novel misos relative to their traditional counterparts without having comparable NGS data for the latter. That being said, one factor that could account for this apparent greater diversity compared to the literature is the kind of space they are made in—a restaurant kitchen where many kinds of fermentation are produced alongside each other. Compared with many traditional producers who often specialise in making one or a few specific kinds of the same fermented product, these kitchen spaces are noteworthy in that the variety of microorganisms present from concurrent fermentation processes could be influencing the microbial composition of the various foods produced therein. This variety could be one reason why these novel misos, despite being made under different circumstances and with different compositions, all exhibit such relatively high microbial diversity. Investigation of different traditional misos using similar methods would be necessary to test this hypothesis.

When it comes to differences between the misos, substrate type, treatment such as nixtamalisation, the fermentation process, and geographical location all shape the bacterial composition to different degrees, with a weaker effect for fungi. The effect of geography seems to vary according to the level of analysis. In the overall statistical analysis based on the metabarcoding data to the genus and/or species level, a small geographical effect is present but is the weakest of the four factors studied for bacteria; for fungi it is the strongest, but at a comparable low level. When it comes to species variation within the same genus, there sometimes appears some degree of geographical effect: among the *Staphylococcus* spp., for example, the predominant species in the Noma misos (aside from the soybean ones) is *S. pasteuri*, while in the Inua misos it is *S. warneri*; among the *Pediococcus* spp., the predominant species are *P. pentosaceus* and *P. acidilactici* respectively. The pangenomic analysis of *S. epidermidis* strains, meanwhile, shows a clear clustering of MAGs based on substrate over geography. Yet focussing in on the carotenoid genes in these MAGs revealed a surprising possible effect of geography, in which the Inua strains across substrates contain carotenoid biosynthesis genes while the Noma strains do not—a pattern supported by the literature.

Concerning the *S. epidermidis*, it is notable that this species is usually associated with the human skin microbiome rather than fermented food. Its presence in and evident adaptation to the miso niche is therefore an illuminating example of the benign, even generative microbial traffic between humans and fermented foods, with consequences for microbial biodiversity, ecology and evolution. Such biological novelty emerging within holobiotic fermentation microbiomes (Theis *et al*., 2016; Dunn *et al*., 2020) likely also has functional impacts on nutrition and flavour—one promising avenue for further research.

Many other directions for further research emerge from this study’s limitations. While our study sheds some further light on the complexities of the microbial geography of fermentation, investigating geographic variation was not its main aim. Our ability to make stronger claims about geographical effect and ‘microbial terroir’ is limited by the lack of replicates for the Inua misos, having only one kind of miso from both Inua and Noma, and having only one species with MAGs that span both locations in the same substrate (and only four of these MAGs). Investigating these dynamics of microbial terroir more directly and robustly is one fruitful direction for further work. Similarly, we were not able here to comment in much detail on the development of the microbial communities over the course of the fermentation, as we were only able to sequence the start and end samples. Further insights here might be gained by sequencing the intermediate samples we took after one month, as well as designing future studies with more sampling throughout for greater resolution of the change in microbial communities over time. And while we made use of our biological replicates to conduct statistical analyses for effects of substrate, time, treatment, and location, we only had space here to include and discuss these analyses for the 16S and ITS data. Future research could do the same using the metagenomics data, to see what might be refined and perhaps what might change. While the current study offers insight into microbial diversity and some of the effects that shape it, it does not offer any mechanistic conclusions— for example in elucidating the mechanisms for the genetic differentiation observed among *S. epidermidis* strains. Further research might investigate specific mechanisms underlying dynamics of microbial composition and genetics in miso fermentation and identify the functional roles of individual microorganisms and microbial interactions in shaping the sensory and nutritional properties of the final product. Finally, it might also, as described in the introduction, investigate all these questions for traditional misos as well, so that we have something to compare the more novel products to. By exploring these questions further, future studies would promote the development and enjoyment of miso and other fermented plant-based foods, facilitating product quality, sustainability, and food diversity, while deepening our knowledge of the microbiology of food fermentation, both traditional and novel.

## Supplementary materials

**Table S1.** Metadata of the 38 samples used in this study with information about the proteinous and kōji substrates to produce the miso, the location of fermentation, the time of sampling (start or end), and the indication of the biological replicate.

**Table S2.** Raw metabarcoding reads detecting bacterial (16S) and fungal (ITS) ASVs in the samples. The colour code indicates abundance of each species, ranging from red (low abundance) to green (high abundance).

**Table S3.** Statistical analyses of the metabarcoding data evaluating the effects of substrate, nixtamalisation, time, and location. The Kruskal-Wallis and Tukey methods were applied for measures of alpha diversity. PERMANOVA and Deseq were used for beta diversity analyses.

**Table S4.** The table is divided in four sheets: (i) ‘<u>reads_quality’</u>, reporting the quality control of the metagenomic shotgun sequences; (ii) ‘metaphan’, the relative abundance of microbial communities in the 38 miso metagenomes using MetaPhlAn taxonomic assignment; (iii) ‘ychF’, abundance table using the *ychF* gene as a marker; and (iv) ‘<u>leuS’</u>, abundance table using the *leuS* gene as a marker. The colour code indicates relative abundance, ranging from red (low abundance) to green (high abundance).

**Table S5**. Quality of prokaryotic and *A. oryzae* MAGs, and their ANIs with the closest reference genome. The colour code indicates the MAGs’ completeness, ranging from low (red) to high (green), and contamination, ranging from low (red) to high (blue).

**Table S6**. ANI analyses for the recovered MAGs of *S. epidermidis*, *P. acidilacti*, *L. plantarum*, *P. pentosaceus* and *S. warneri*. The colour code indicates the ANI values ranging from low (red) to high (green).

**Table S7**. Unique genes annotated with BlastKoala KEGG from the recovered MAGs of *S. epidermidis* from the yellow pea group and maitake-soy strain.

## Acknowledgements

First and foremost, we would like to acknowledge the countless generations of miso makers, domestic and professional, within and outside of Japan, without whose knowledge and craft this study would not be possible and to which it attempts to contribute.

We would like to thank Jason White, at the time the deputy head of the Fermentation Lab at Restaurant Noma under DZ, for being involved in designing the experiment with JE and for helping him with its planning and execution. We would also like to thank Lars Williams, founding head of the Noma Fermentation Lab, for originally developing Noma’s miso recipes and production; René Redzepi, chef and owner of Restaurant Noma, for allowing JE to carry out research there; Thomas Frebel, then head chef of Restaurant Inua in Tokyo, for supplying us with samples of misos; and Risa Kamio for transporting these samples to Copenhagen. Thanks also to Taeko Hamada for locating and translating the Ogo paper (Ogo 1985) important for the discussion of strain differentiation.

This research has been funded by the Mortimer May DPhil Scholarship in Human Geography at Hertford College, University of Oxford, which funded JE’s PhD research; M Thomas P Gilbert’s grant from the Danish National Research Foundation, grant number DNRF143, that covered the metabarcoding sequencing; The Novo Nordisk Foundation, grant number NNF20CC0035580, that covered the metagenomics sequencing and the salaries of CIK, PCM, and JE; and a Banting postdoctoral fellowship from the Canadian government as well as the Swiss National Science Foundation and University of Lausanne that funded FM.

## Author Contributions

JE designed and carried out the experiment. DZ helped design and facilitate the experiment. CC supervised JE in the lab and taught him how to conduct the lab work for the metabarcoding. FM conducted the preliminary metabarcoding analyses. CIK generated the metagenomics data, redid all analyses, and prepared the figures. PCM and NM helped CIK with analyses of the metagenomic and genomic data. JE funded the metagenomics data generation. CIK wrote the first draft of the manuscript. JE and CIK elaborated the introduction, discussion, and conclusion sections together and revised the manuscript. All authors had the opportunity to read, comment on, and approve the manuscript before submission.

