## Supplementary material for "Novel misos shape distinct microbial ecologies: opportunities for flavourful sustainable food innovation": Supplemetary materials

**Table S1.** Metadata of the 38 samples used in this study with information about the proteinous and kōji substrates to produce the miso, the location of fermentation, the time of sampling (start or end), and the indication of the biological replicate.

| Sample | Biological replicate | Proteinous substrate | Spores | Kōji substrate | Made by | Country | Fermentation | pH |
|---|---|---|---|---|---|---|---|---|
| F.1 | 1 | Fava bean | Rice kōji | Pearled barley | NOMA | Denmark | Start | 5.53 |
| NF.1 | 1 | Nixtamalised fava bean | Rice kōji | Pearled barley | NOMA | Denmark | Start | 5.53 |
| L.1 | 1 | Gotland lentil | Rice kōji | Pearled barley | NOMA | Denmark | Start | 5.13 |
| NL.1 | 1 | Nixtamalised Gotland lentil | Rice kōji | Pearled barley | NOMA | Denmark | Start | 5.53 |
| P.1 | 1 | Yellow pea | Rice kōji | Pearled barley | NOMA | Denmark | Start | 5.13 |
| NP.1 | 1 | Nixtamalised yellow pea | Rice kōji | Pearled barley | NOMA | Denmark | Start | 5.53 |
| R.1 | 1 | Rye bread | Rice kōji | Pearled barley | NOMA | Denmark | Start | 4.52 |
| S.1 | 1 | Soybean | Rice kōji | Pearled barley | NOMA | Denmark | Start | 5.64 |
| F1.3 | 1 | Fava bean | Rice kōji | Pearled barley | NOMA | Denmark | End | 4.42 |
| F2.3 | 2 | Fava bean | Rice kōji | Pearled barley | NOMA | Denmark | End | 4.62 |
| F3.3 | 3 | Fava bean | Rice kōji | Pearled barley | NOMA | Denmark | End | 3.72 |
| NF1.3 | 1 | Nixtamalised fava bean | Rice kōji | Pearled barley | NOMA | Denmark | End | 3.62 |
| NF2.3 | 2 | Nixtamalised fava bean | Rice kōji | Pearled barley | NOMA | Denmark | End | 4.02 |
| NF3.3 | 3 | Nixtamalised fava bean | Rice kōji | Pearled barley | NOMA | Denmark | End | 3.22 |
| L1.3 | 1 | Gotland lentil | Rice kōji | Pearled barley | NOMA | Denmark | End | 4.32 |
| L2.3 | 2 | Gotland lentil | Rice kōji | Pearled barley | NOMA | Denmark | End | 5.02 |
| L3.3 | 3 | Gotland lentil | Rice kōji | Pearled barley | NOMA | Denmark | End | 3.42 |
| NL1.3 | 1 | Nixtamalised Gotland lentil | Rice kōji | Pearled barley | NOMA | Denmark | End | 3.82 |
| NL2.3 | 2 | Nixtamalised Gotland lentil | Rice kōji | Pearled barley | NOMA | Denmark | End | 4.12 |
| NL3.3 | 3 | Nixtamalised Gotland lentil | Rice kōji | Pearled barley | NOMA | Denmark | End | 3.32 |
| P1.3 | 1 | Yellow pea | Rice kōji | Pearled barley | NOMA | Denmark | End | 4.12 |
| P2.3 | 2 | Yellow pea | Rice kōji | Pearled barley | NOMA | Denmark | End | 4.72 |
| P3.3 | 3 | Yellow pea | Rice kōji | Pearled barley | NOMA | Denmark | End | 4.02 |
| NP1.3 | 1 | Nixtamalised yellow pea | Rice kōji | Pearled barley | NOMA | Denmark | End | 4.02 |
| NP2.3 | 2 | Nixtamalised yellow pea | Rice kōji | Pearled barley | NOMA | Denmark | End | 4.12 |
| NP3.3 | 3 | Nixtamalised yellow pea | Rice kōji | Pearled barley | NOMA | Denmark | End | 3.42 |
| R1.3 | 1 | Rye bread | Rice kōji | Pearled barley | NOMA | Denmark | End | 4.12 |
| R2.3 | 2 | Rye bread | Rice kōji | Pearled barley | NOMA | Denmark | End | 4.12 |
| R3.3 | 3 | Rye bread | Rice kōji | Pearled barley | NOMA | Denmark | End | 3.72 |
| S1.3 | 1 | Soybean | Rice kōji | Rice | NOMA | Denmark | End | 4.32 |
| S2.3 | 2 | Soybean | Rice kōji | Rice | NOMA | Denmark | End | 4.32 |
| S3.3 | 3 | Soybean | Rice kōji | Rice | NOMA | Denmark | End | 3.72 |
| HB | 1 | Habanero-barley | Barley kōji | Pearled barley | INUA | Japan | End | - |
| MS | 1 | Maitake-soy | Barley kōji | Pearled barley | INUA | Japan | End | - |
| PB | 1 | Yellow pea | Barley kōji | Pearled barley | INUA | Japan | End | - |
| PR | 1 | Yellow pea | Barley kōji | Rice | INUA | Japan | End | - |
| SY | 1 | Soy-yuzu pulp | Barley kōji | Pearled barley | INUA | Japan | End | - |
| TS | 1 | Toasted sesame | Barley kōji | Pearled barley | INUA | Japan | End | - |

| | start | end |
|---|---|---|
| **mean pH** | 5.3175 | 4.02 |
| **SD** | 0.376250297 | 0.449907 |

**Table S2**. Raw metabarcoding reads detecting bacterial (16S) and fungal (ITS) ASVs in the samples. The colour code indicates abundance of each species, ranging from red (low abundance) to green (high abundance).

| | | | | | | | | | | Fermentation | start | start | start |
|---|---|---|---|---|---|---|---|---|---|---|---|---|---|
| | | | | | | | | | | Technical replicate | a | b | c |
| | | | | | | | | | | | fava start | fava start | fava start |
| Kingdom | Phylum | Class | Order | Family | Genus | Species | Description sequence | ASV | | sum | F.1a | F.1b | F.1c |
| Bacteria | Firmicutes | Bacilli | Lactobacillales | Lactobacillaceae | Pediococcus_1 | Pediococcus pentosaceus | TACGTAG | ASV | 1 | 1895684 | 9 | 0 | 0 |
| Bacteria | Firmicutes | Bacilli | Bacillales | Staphylococcaceae | Staphylococcus | Staphylococcus sp. | TACGTAG | ASV | 2 | 920589 | 9 | 0 | 20 |
| Bacteria | Proteobacteria | Gammaproteol | Enterobacteriales | Enterobacteriaceae | Enterobacteriaceae far | Enterobacteriaceae family | TACGGAG | ASV | 3 | 565458 | 1801 | 3047 | 3265 |
| Bacteria | Firmicutes | Bacilli | Bacillales | Bacillaceae | Bacillus | Bacillus sp. | TACGTAG | ASV | 4 | 450746 | 1703 | 2015 | 2260 |
| Bacteria | Firmicutes | Bacilli | Lactobacillales | Lactobacillaceae | Pediococcus_2 | Pediococcus acidilactici | TACGTAG | ASV | 5 | 348297 | 0 | 0 | 0 |
| Bacteria | Proteobacteria | Gammaproteol | Enterobacteriales | Enterobacteriaceae | Enterobacteriaceae far | Enterobacteriaceae family | TACGGAG | ASV | 6 | 154047 | 1813 | 3114 | 3429 |
| Bacteria | Firmicutes | Bacilli | Lactobacillales | Lactobacillaceae | Lactiplantibacillus | Lactiplantibacillus sp. | TACGTAG | ASV | 8 | 120779 | 0 | 0 | 0 |
| Bacteria | Firmicutes | Bacilli | Bacillales | Bacillaceae | Bacillus | Bacillus sp. | TACGTAG | ASV | 10 | 110337 | 15 | 14 | 10 |
| Bacteria | Firmicutes | Bacilli | Lactobacillales | Lactobacillaceae | Lacticaseibacillus | Lacticaseibacillus sp. | TACGTAG | ASV | 11 | 1399 | 13 | 9 | 16 |
| Bacteria | Firmicutes | Bacilli | Lactobacillales | Enterococcaceae | Tetragenococcus | Tetragenococcus halophilus | TACGTAG | ASV | 13 | 106657 | 0 | 0 | 0 |
| Bacteria | Firmicutes | Bacilli | Lactobacillales | Enterococcaceae | Enterococcus | Enterococcus sp. | TACGTAG | ASV | 14 | 62993 | 0 | 0 | 0 |
| Bacteria | Firmicutes | Bacilli | Bacillales | Bacillaceae | Bacillus | Bacillus sp. | TACGTAG | ASV | 16 | 51096 | 150 | 192 | 214 |
| Bacteria | Firmicutes | Bacilli | Bacillales | Bacillaceae | Bacillus | Bacillus sp. | TACGTAG | ASV | 17 | 49994 | 177 | 212 | 256 |
| Bacteria | Firmicutes | Bacilli | Lactobacillales | Leuconostocaceae | Weissella | Weissella sp. | TACGTAT | ASV | 19 | 8051 | 0 | 0 | 0 |
| Bacteria | Firmicutes | Bacilli | Lactobacillales | Lactobacillaceae | Pediococcus | Pediococcus pentosaceus | TACGTAG | ASV | 22 | 26714 | 0 | 0 | 0 |
| Bacteria | Proteobacteria | Alphaproteoba | Acetobacterales | Acetobacteraceae | Komagataeibacter | Komagataeibacter rhaeticus | TACGAAG | ASV | 23 | 30 | 0 | 0 | 0 |
| Bacteria | Proteobacteria | Alphaproteoba | Acetobacterales | Acetobacteraceae | Acetobacter | Acetobacter sp. | TACGAAG | ASV | 24 | 30 | 0 | 4 | 0 |
| Bacteria | Proteobacteria | Gammaproteol | Enterobacteriales | Enterobacteriaceae | Enterobacteriaceae far | Enterobacteriaceae family | TACGGAG | ASV | 26 | 18880 | 307 | 437 | 489 |
| Bacteria | Proteobacteria | Gammaproteol | Enterobacteriales | Enterobacteriaceae | Enterobacteriaceae far | Enterobacteriaceae family | TACGGAG | ASV | 28 | 15415 | 0 | 54 | 53 |
| Bacteria | Firmicutes | Bacilli | Lactobacillales | Lactobacillaceae | Latilactobacillus | Latilactobacillus sp. | TACGTAG | ASV | 29 | 13851 | 0 | 0 | 0 |
| Bacteria | Proteobacteria | Alphaproteoba | Acetobacterales | Acetobacteraceae | Gluconobacter | Gluconobacter sp. | TACGAAG | ASV | 30 | 6 | 0 | 0 | 0 |
| Bacteria | Firmicutes | Bacilli | Lactobacillales | Streptococcaceae | Lactococcus | Lactococcus sp. | TACGTAG | ASV | 31 | 406 | 0 | 3 | 0 |
| Bacteria | Proteobacteria | Gammaproteol | Betaproteobacteri | Neisseriaceae | Neisseria | Neisseria sp. | TACGTAG | ASV | 32 | 6 | 0 | 0 | 0 |
| Bacteria | Firmicutes | Bacilli | Bacillales | Staphylococcaceae | Staphylococcus | Staphylococcus sp. | TACGTAG | ASV | 33 | 9401 | 0 | 0 | 0 |
| Bacteria | Firmicutes | Bacilli | Lactobacillales | Lactobacillaceae | Pediococcus | Pediococcus pentosaceus gro | TACGTAG | ASV | 34 | 9630 | 0 | 0 | 0 |
| Bacteria | Firmicutes | Bacilli | Lactobacillales | Streptococcaceae | Streptococcus | Streptococcus thermophilus- | TACGTAG | ASV | 36 | 157 | 0 | 3 | 0 |
| Bacteria | Firmicutes | Bacilli | Bacillales | Staphylococcaceae | Staphylococcus | Staphylococcus sp. | TACGTAG | ASV | 37 | 7569 | 0 | 0 | 0 |
| Bacteria | Firmicutes | Bacilli | Lactobacillales | Leuconostocaceae | Weissella | Weissella sp. | TACGTAT | ASV | 38 | 6355 | 0 | 0 | 0 |
| Bacteria | Actinobacteria | Actinobacteria | Micrococcales | Micrococcaceae | Micrococcus | Micrococcus sp. | TACGTAG | ASV | 39 | 66 | 0 | 0 | 0 |
| Bacteria | Firmicutes | Bacilli | Lactobacillales | Leuconostocaceae | Weissella | Weissella sp. | TACGTAT | ASV | 40 | 5596 | 0 | 0 | 0 |
| Bacteria | Firmicutes | Bacilli | Lactobacillales | Lactobacillaceae | Pediococcus | Pediococcus pentosaceus gro | TACGTAG | ASV | 41 | 5587 | 0 | 0 | 0 |
| Bacteria | Actinobacteria | Actinobacteria | Corynebacteriales | Corynebacteriaceae | Lawsonella | Lawsonella clevelandensis | TACGTAG | ASV | 43 | 18 | 0 | 0 | 0 |
| Bacteria | Proteobacteria | Gammaproteol | Enterobacteriales | Enterobacteriaceae | Enterobacteriaceae far | Enterobacteriaceae family | TACGGAG | ASV | 44 | 4640 | 0 | 0 | 0 |
| Bacteria | Proteobacteria | Gammaproteol | Betaproteobacteri | Neisseriaceae | Neisseria | Neisseria sp. | TACGTAG | ASV | 46 | 2 | 0 | 0 | 0 |
| Bacteria | Firmicutes | Bacilli | Bacillales | Staphylococcaceae | Staphylococcus | Staphylococcus sp. | TACGTAG | ASV | 47 | 4382 | 0 | 0 | 0 |
| Bacteria | Firmicutes | Bacilli | Bacillales | Staphylococcaceae | Staphylococcus | Staphylococcus sp. | TACGTAG | ASV | 48 | 3830 | 0 | 0 | 0 |
| Bacteria | Firmicutes | Bacilli | Lactobacillales | Lactobacillaceae | Lacticaseibacillus | Lacticaseibacillus sp. | TACGTAG | ASV | 52 | 88 | 0 | 0 | 0 |
| Bacteria | Firmicutes | Bacilli | Lactobacillales | Lactobacillaceae | Lacticaseibacillus | Lacticaseibacillus sp. | TACGTAG | ASV | 53 | 38 | 0 | 0 | 0 |
| Bacteria | Proteobacteria | Gammaproteol | Pseudomonadales | Pseudomonadaceae | Pseudomonas | Pseudomonas sp. | TACAGAG | ASV | 56 | 37 | 0 | 0 | 0 |
| Bacteria | Firmicutes | Bacilli | Bacillales | Staphylococcaceae | Staphylococcus | Staphylococcus sp. | TACGTAG | ASV | 57 | 2672 | 0 | 0 | 0 |
| Bacteria | Firmicutes | Bacilli | Bacillales | Bacillaceae | Bacillus | Bacillus sp. | TACGTAG | ASV | 58 | 2442 | 0 | 0 | 0 |
| Bacteria | Proteobacteria | Gammaproteol | Enterobacteriales | Enterobacteriaceae | Enterobacteriaceae far | Enterobacteriaceae family | TACGGAG | ASV | 62 | 2307 | 0 | 0 | 0 |
| Bacteria | Proteobacteria | Gammaproteol | Pseudomonadales | Moraxellaceae | Acinetobacter | Acinetobacter sp. | TACAGAG | ASV | 64 | 2079 | 0 | 0 | 0 |
| Bacteria | Proteobacteria | Gammaproteol | Pseudomonadales | Moraxellaceae | Acinetobacter | Acinetobacter sp. | TACAGAG | ASV | 66 | 1987 | 0 | 0 | 0 |
| Bacteria | Proteobacteria | Gammaproteol | Enterobacteriales | Enterobacteriaceae | Enterobacteriaceae far | Enterobacteriaceae family | TACGGAG | ASV | 67 | 1957 | 0 | 0 | 0 |
| Bacteria | Firmicutes | Bacilli | Bacillales | Bacillaceae | Bacillus | Bacillus sp. | TACGTAG | ASV | 68 | 1749 | 0 | 0 | 0 |
| Bacteria | Firmicutes | Bacilli | Bacillales | Bacillaceae | Bacillus | Bacillus sp. | TACGTAG | ASV | 71 | 1570 | 0 | 0 | 0 |
| Bacteria | Firmicutes | Bacilli | Bacillales | Staphylococcaceae | Staphylococcus | Staphylococcus sp. | TACGTAG | ASV | 72 | 1478 | 0 | 0 | 0 |
| Bacteria | Proteobacteria | Gammaproteol | Enterobacteriales | Enterobacteriaceae | Enterobacteriaceae far | Enterobacteriaceae family | TACGGAG | ASV | 73 | 1360 | 0 | 0 | 0 |
| Bacteria | Firmicutes | Bacilli | Lactobacillales | Lactobacillaceae | Agrilactobacillus | Agrilactobacillus composti | TACGTAG | ASV | 75 | 1375 | 0 | 0 | 0 |
| Bacteria | Proteobacteria | Gammaproteol | Pseudomonadales | Moraxellaceae | Psychrobacter | Psychrobacter immobilis grou | TACAGAG | ASV | 78 | 1200 | 0 | 0 | 0 |
| Bacteria | Firmicutes | Bacilli | Bacillales | Bacillaceae | Bacillus | Bacillus sp. | TACGTAG | ASV | 80 | 1174 | 0 | 0 | 0 |
| Bacteria | Firmicutes | Bacilli | Lactobacillales | Leuconostocaceae | Weissella | unknown species | TACGTAT | ASV | 84 | 1038 | 0 | 0 | 0 |
| Bacteria | Firmicutes | Bacilli | Bacillales | Bacillaceae | Bacillus | Bacillus sp. | TACGTAG | ASV | 85 | 1047 | 0 | 0 | 0 |
| Bacteria | Firmicutes | Bacilli | Bacillales | Bacillaceae | Virgibacillus | Virgibacillus sp. | TACGTAG | ASV | 86 | 1005 | 0 | 0 | 0 |
| Bacteria | Firmicutes | Bacilli | Lactobacillales | Lactobacillaceae | Pediococcus | unknown species | TACGTAG | ASV | 87 | 982 | 0 | 0 | 0 |
| Bacteria | Firmicutes | Bacilli | Bacillales | Staphylococcaceae | Staphylococcus | Staphylococcus sp. | TACGTAG | ASV | 88 | 988 | 0 | 0 | 0 |
| Bacteria | Firmicutes | Bacilli | Lactobacillales | Leuconostocaceae | Weissella | unknown species | TACGTAT | ASV | 90 | 907 | 0 | 0 | 0 |
| Bacteria | Firmicutes | Bacilli | Bacillales | Staphylococcaceae | Staphylococcus | Staphylococcus sp. | TACGTAG | ASV | 91 | 856 | 0 | 0 | 0 |
| Bacteria | Proteobacteria | Gammaproteol | Pseudomonadales | Moraxellaceae | Moraxella | Moraxella osloensis | TACAGAG | ASV | 92 | 194 | 0 | 0 | 0 |
| Bacteria | Firmicutes | Bacilli | Lactobacillales | Enterococcaceae | Enterococcus | Enterococcus sp. | TACGTAG | ASV | 94 | 738 | 0 | 0 | 0 |
| Bacteria | Proteobacteria | Gammaproteol | Xanthomonadales | Xanthomonadaceae | Stenotrophomonas | Stenotrophomonas sp. | TACGAAG | ASV | 97 | 32 | 0 | 0 | 0 |
| Bacteria | Firmicutes | Bacilli | Bacillales | Bacillaceae | Bacillus | Bacillus sp. | TACGTAG | ASV | 98 | 629 | 0 | 0 | 0 |
| Bacteria | Firmicutes | Bacilli | Lactobacillales | Lactobacillaceae | Levilactobacillus | Levilactobacillus sp. | TACGTAG | ASV | 99 | 547 | 0 | 0 | 0 |
| Bacteria | Proteobacteria | Gammaproteol | Pseudomonadales | Moraxellaceae | Acinetobacter | Acinetobacter sp. | TACAGAG | ASV | 100 | 579 | 0 | 0 | 0 |
| Bacteria | Proteobacteria | Gammaproteol | Pseudomonadales | Pseudomonadaceae | Pseudomonas | Pseudomonas sp. | TACAGAG | ASV | 101 | 603 | 0 | 0 | 0 |
| Bacteria | Firmicutes | Bacilli | Bacillales | Bacillaceae | Bacillus | Bacillus sp. | TACGTAG | ASV | 104 | 516 | 0 | 0 | 0 |
| Bacteria | Firmicutes | Bacilli | Lactobacillales | Lactobacillaceae | Lactiplantibacillus | Lactiplantibacillus sp. | TACGTAG | ASV | 107 | 471 | 0 | 0 | 0 |
| Bacteria | Proteobacteria | Gammaproteol | Pseudomonadales | Moraxellaceae | Acinetobacter | Acinetobacter sp. | TACAGAG | ASV | 108 | 455 | 9 | 0 | 10 |
| Bacteria | Firmicutes | Bacilli | Bacillales | Staphylococcaceae | Mammaliicoccus | Mammaliicoccus sp. | TACGTAG | ASV | 109 | 436 | 0 | 0 | 0 |
| Bacteria | Proteobacteria | Gammaproteol | Xanthomonadales | Xanthomonadaceae | Stenotrophomonas-Ps | unknown species | TACGAAG | ASV | 111 | 438 | 0 | 0 | 0 |
| Bacteria | Firmicutes | Bacilli | Lactobacillales | Streptococcaceae | Streptococcus | Streptococcus sp. | TACGTAG | ASV | 112 | 171 | 0 | 0 | 0 |
| Bacteria | Firmicutes | Bacilli | Lactobacillales | Leuconostocaceae | Leuconostoc | Leuconostoc sp. | TACGTAT | ASV | 113 | 425 | 0 | 0 | 0 |
| Bacteria | Proteobacteria | Gammaproteol | Pseudomonadales | Pseudomonadaceae | Pseudomonas | Pseudomonas sp. | TACAGAG | ASV | 116 | 390 | 0 | 0 | 0 |
| Bacteria | Firmicutes | Bacilli | Bacillales | Bacillaceae | Niallia | Niallia sp. | TACGTAG | ASV | 117 | 386 | 0 | 0 | 0 |
| Bacteria | Firmicutes | Bacilli | Lactobacillales | Lactobacillaceae | Lacticaseibacillus | Lacticaseibacillus saniviri | TACGTAG | ASV | 118 | 383 | 0 | 0 | 0 |
| Bacteria | Firmicutes | Bacilli | Lactobacillales | Carnobacteriaceae | Carnobacterium | Carnobacterium sp. | TACGTAG | ASV | 119 | 381 | 0 | 0 | 0 |
| Bacteria | Firmicutes | Bacilli | Bacillales | Staphylococcaceae | Staphylococcus | Staphylococcus sp. | TACGTAG | ASV | 120 | 376 | 0 | 0 | 0 |
| Bacteria | Proteobacteria | Gammaproteol | Pseudomonadales | Moraxellaceae | Acinetobacter | Acinetobacter sp. | TACAGAG | ASV | 122 | 366 | 0 | 0 | 0 |
| Bacteria | Firmicutes | Bacilli | Bacillales | Bacillaceae | Priestia-Bacillus | Priestia-Bacillus genera | TACGTAG | ASV | 123 | 360 | 0 | 0 | 0 |
| Bacteria | Proteobacteria | Gammaproteol | Enterobacteriales | Enterobacteriaceae | Enterobacteriaceae far | Enterobacteriaceae family | TACGGAG | ASV | 124 | 348 | 0 | 0 | 0 |
| Bacteria | Proteobacteria | Gammaproteol | Pseudomonadales | Moraxellaceae | Acinetobacter | Acinetobacter sp. | TACAGAG | ASV | 130 | 303 | 0 | 0 | 0 |
| Bacteria | Firmicutes | Bacilli | Bacillales | Listeriaceae | Brochothrix | Brochothrix thermosphacta | TACGTAG | ASV | 132 | 300 | 0 | 0 | 0 |
| Bacteria | Proteobacteria | Gammaproteol | Enterobacteriales | Enterobacteriaceae | Enterobacteriaceae far | Enterobacteriaceae family | TACGGAG | ASV | 136 | 248 | 0 | 0 | 0 |
| Bacteria | Actinobacteria | Actinobacteria | Micrococcales | Intrasporangiaceae | Janibacter | Janibacter sp. | TACGTAG | ASV | 138 | 43 | 0 | 0 | 0 |
| Bacteria | Firmicutes | Bacilli | Lactobacillales | Leuconostocaceae | Weissella | Weisella sp. | TACGTAT | ASV | 140 | 229 | 0 | 0 | 0 |
| Bacteria | Proteobacteria | Gammaproteol | Enterobacteriales | Enterobacteriaceae | Enterobacteriaceae far | Enterobacteriaceae family | TACGGAG | ASV | 142 | 227 | 0 | 77 | 0 |
| Bacteria | Proteobacteria | Gammaproteol | Enterobacteriales | Enterobacteriaceae | Enterobacteriaceae far | Enterobacteriaceae family | TACGGAG | ASV | 145 | 202 | 0 | 0 | 0 |
| Bacteria | Bacteroidetes | Bacteroidia | Flavobacteriales | Weeksellaceae | Chryseobacterium | Chryseobacterium sp. | TACGGAG | ASV | 146 | 194 | 0 | 0 | 0 |
| Bacteria | Firmicutes | Bacilli | Bacillales | Staphylococcaceae | Staphylococcus | Staphylococcus sp. | TACGTAG | ASV | 152 | 177 | 0 | 0 | 0 |
| Bacteria | Actinobacteria | Actinobacteria | Micrococcales | Micrococcaceae | Arthrobacter | Arthrobacter sp. | TACGTAG | ASV | 155 | 171 | 0 | 0 | 0 |
| Bacteria | Proteobacteria | Gammaproteol | Enterobacteriales | Enterobacteriaceae | Enterobacteriaceae far | Enterobacteriaceae family | TACGGAG | ASV | 160 | 160 | 0 | 0 | 0 |
| Bacteria | Firmicutes | Bacilli | Bacillales | Staphylococcaceae | Staphylococcus | Staphylococcus sp. | TACGTAG | ASV | 170 | 116 | 0 | 0 | 0 |
| Bacteria | Actinobacteria | Actinobacteria | Corynebacteriales | Corynebacteriaceae | Corynebacterium | Corynebacterium lipophilofla | TACGTAG | ASV | 172 | 105 | 0 | 0 | 0 |
| Bacteria | Firmicutes | Bacilli | Bacillales | Bacillaceae | Bacillus | Bacillus sp. | TACGTAG | ASV | 173 | 105 | 0 | 0 | 0 |
| Bacteria | Firmicutes | Bacilli | Bacillales | Bacillaceae | Oceanobacillus | Oceanobacillus sp. | TACGTAG | ASV | 178 | 92 | 0 | 0 | 0 |
| Bacteria | Proteobacteria | Gammaproteol | Aeromonadales | Aeromonadaceae | Aeromonas | Aeromonas sp. | TACGGAG | ASV | 180 | 89 | 0 | 0 | 0 |
| Bacteria | Proteobacteria | Gammaproteol | Betaproteobacteri | Hydrogenophilaceae | Tepidiphilus | Tepidiphilus succinatimander | TACGTAG | ASV | 181 | 88 | 0 | 0 | 0 |
| Bacteria | Proteobacteria | Gammaproteol | Pseudomonadales | Moraxellaceae | Psychrobacter | Psychrobacter sp. | TACAGAG | ASV | 187 | 71 | 0 | 0 | 0 |
| Bacteria | Bacteroidetes | Bacteroidia | Sphingobacteriales | Sphingobacteriaceae | Sphingobacterium | Sphingobacterium sp. | TACGGAG | ASV | 189 | 70 | 0 | 0 | 0 |
| Bacteria | Actinobacteria | Actinobacteria | Micrococcales | Dermabacteraceae | Brachybacterium | unknown species | TACGTAG | ASV | 197 | 60 | 0 | 0 | 0 |
| Bacteria | Firmicutes | Bacilli | Lactobacillales | Leuconostocaceae | Leuconostoc | Leuconostoc sp. | TACGTAT | ASV | 206 | 51 | 0 | 0 | 0 |
| Bacteria | Proteobacteria | Gammaproteol | Enterobacteriales | Enterobacteriaceae | Enterobacteriaceae far | Enterobacteriaceae family | TACGGAG | ASV | 207 | 50 | 0 | 0 | 0 |
| Bacteria | Bacteroidetes | Bacteroidia | Bacteroidales | Prevotellaceae | Ihuprevotella | Ihuprevotella massiliensis | TACGGAA | ASV | 209 | 45 | 0 | 0 | 0 |
| Bacteria | Firmicutes | Bacilli | Bacillales | Paenibacillaceae | Brevibacillus | Brevibacillus sp. | TACGTAG | ASV | 210 | 45 | 0 | 0 | 0 |
| Bacteria | Proteobacteria | Gammaproteol | Xanthomonadales | Xanthomonadaceae | Stenotrophomonas | unknown species | TACGAAG | ASV | 216 | 40 | 0 | 0 | 0 |
| Bacteria | Actinobacteria | Actinobacteria | Micrococcales | Sanguibacteraceae | Sanguibacter | Sanguibacter inulinus-keddie | TACGTAG | ASV | 217 | 29 | 0 | 0 | 0 |
| Bacteria | Firmicutes | Bacilli | Bacillales | Staphylococcaceae | Mammaliicoccus | Mammaliicoccus sp. | TACGTAG | ASV | 222 | 34 | 0 | 0 | 0 |
| Bacteria | Proteobacteria | Gammaproteol | Enterobacteriales | Enterobacteriaceae | Enterobacteriaceae far | Enterobacteriaceae family | TACGGAG | ASV | 228 | 30 | 0 | 0 | 0 |
| Bacteria | Proteobacteria | Gammaproteol | Betaproteobacteri | Burkholderiaceae | Schlegelella | Schlegelella aquatica-thermo | TACGTAG | ASV | 235 | 26 | 0 | 0 | 0 |
| Bacteria | Proteobacteria | Gammaproteol | Pseudomonadales | Pseudomonadaceae | Pseudomonas | Pseudomonas sp. | TACGAAG | ASV | 238 | 25 | 0 | 0 | 0 |
| Bacteria | Actinobacteria | Actinobacteria | Micrococcales | Micrococcaceae | Rothia | Rothia kristinae | TACGTAG | ASV | 244 | 22 | 0 | 0 | 0 |
| Bacteria | Proteobacteria | Gammaproteol | Pseudomonadales | Pseudomonadaceae | Pseudomonas | Pseudomonas sp. | TACAGAG | ASV | 249 | 18 | 0 | 0 | 0 |
| Bacteria | Firmicutes | Bacilli | Lactobacillales | Leuconostocaceae | Weissella | Weissella sp. | TACGTAT | ASV | 253 | 8 | 0 | 0 | 0 |
| Bacteria | Firmicutes | Bacilli | Bacillales | Paenibacillaceae | Brevibacillus | Brevibacillus sp. | TACGTAG | ASV | 274 | 11 | 0 | 0 | 0 |
| | | | | | | | | | | | 6006 | 9181 | 10022 |

**Table S2.** Raw metabarcoding reads detecting bacterial (16S) and fungal (ITS) ASVs in the samples. The colour code indicates abundance of each species, ranging from red (low abundance) to green (high abundance).

| Kingdo | Phylum | Class | Order | Family | Genus | Species | Description sequence | ASV |
|---|---|---|---|---|---|---|---|---|
| Fungi | Ascomycota | Eurotiomycetes | Eurotiales | Aspergillaceae | Aspergillus | Aspergillus oryzae | ATGCGATAACT | ASV_1 |
| Fungi | Ascomycota | Saccharomycetes | Saccharomycetale | Debaryomycetace | Millerozyma | Millerozyma farinosa | ATGCGATAAGT | ASV_2 |
| Fungi | Ascomycota | Saccharomycetes | Saccharomycetale | Saccharomycetac | Starmerella-Cand | Starmerella-Candida genera | GCGCGATAGGT | ASV_4 |
| Fungi | Ascomycota | Saccharomycetes | Saccharomycetale | Saccharomycetac | Zygosaccharomyc | Zygosaccharomyces parabail | ATGCGATACGT | ASV_12 |
| Fungi | Ascomycota | Saccharomycetes | Saccharomycetale | Saccharomycetac | Nakaseomyces | [Candida] glabrata | ATGCGATACGT | ASV_13 |
| Fungi | Ascomycota | Saccharomycetes | Saccharomycetale | Saccharomycetac | Nakaseomyces | [Candida] glabrata | ATGCGATACGT | ASV_14 |
| Fungi | Ascomycota | Eurotiomycetes | Eurotiales | Aspergillaceae | Aspergillus | Aspergillus oryzae-flavus | ATGCGATAACT | ASV_15 |
| Fungi | Ascomycota | Saccharomycetes | Saccharomycetale | Debaryomycetace | Debaryomyces | Debaryomyces hansenii | ATGCGATAAGT | ASV_16 |
| Fungi | Ascomycota | Saccharomycetes | Saccharomycetale | Saccharomycetac | Nakaseomyces | [Candida] glabrata | ATGCGATACGT | ASV_17 |
| Fungi | Ascomycota | Saccharomycetes | Saccharomycetale | Saccharomycetac | Zygosaccharomyc | Zygosaccharomyces rouxii | CTGCGATACGT | ASV_18 |
| Fungi | Ascomycota | Saccharomycetes | Saccharomycetale | Phaffomycetacea | Candida | Candida sake | ATGCGATACGT | ASV_20 |
| Fungi | Ascomycota | Schizosaccharom | Schizosaccharom | Schizosaccharom | Schizosaccharom | Schizosaccharomyces pombe | ATGCGATACGT | ASV_21 |
| Fungi | Ascomycota | Schizosaccharom | Schizosaccharom | Schizosaccharom | Schizosaccharom | Schizosaccharomyces pombe | ATGCGATACGT | ASV_22 |
| Fungi | Ascomycota | Saccharomycetes | Saccharomycetale | Saccharomycetac | Zygosaccharomyc | Zygosaccharomyces bisporu | ATGCGATACGT | ASV_23 |
| Fungi | Ascomycota | Saccharomycetes | Saccharomycetale | Saccharomycetac | Zygosaccharomyc | Zygosaccharomyces bisporu | ATGCGATACGT | ASV_25 |
| Fungi | Ascomycota | Saccharomycetes | Saccharomycetale | Saccharomycetac | Zygosaccharomyc | Zygosaccharomyces parabail | ATGCGATACGT | ASV_26 |
| Fungi | Ascomycota | Saccharomycetes | Saccharomycetale | Debaryomycetace | Debaryomyces | Debaryomyces hansenii | ATGCGATAAGT | ASV_27 |
| Fungi | Ascomycota | Saccharomycetes | Saccharomycetale | Saccharomycopsi | Saccharomycopsi | Saccharomycopsis fibuligera | TTGCGATAAGT | ASV_28 |
| Fungi | Ascomycota | Saccharomycetes | Saccharomycetale | Debaryomycetace | Millerozyma | Millerozyma farinosa | ATGCGATAAGT | ASV_29 |
| Fungi | Basidiomyco | Tremellomycetes | Cystofilobasidiale | Cyfstofilobasidiac | Tausonia | Tausonia pullulans | ATGCGATAAGT | ASV_30 |
| Fungi | Basidiomyco | Agaricomycetes | Agaricales | Omphalotaceae | Lentinula | Lentinula edodes | ATGCGATAAGT | ASV_31 |
| Fungi | Ascomycota | Saccharomycetes | Saccharomycetale | Saccharomycetac | Nakaseomyces | [Candida] glabrata | ATGCGATACGT | ASV_34 |
| Fungi | Ascomycota | Saccharomycetes | Saccharomycetale | Saccharomycetac | Nakaseomyces | [Candida] glabrata | ATGCGATACGT | ASV_35 |
| Fungi | Ascomycota | Eurotiomycetes | Eurotiales | Aspergillaceae | Aspergillus | Aspergillus sp. | ATGCGATAATT | ASV_37 |
| Fungi | Ascomycota | Saccharomycetes | Saccharomycetale | Saccharomycetac | Nakaseomyces | [Candida] glabrata | GATACGTAATG | ASV_39 |
| Fungi | Basidiomyco | Tremellomycetes | Cystofilobasidiale | Cystofilobasidiace | Cystofilobasidiu | Cystofilobasidium capitatum | TTGCGATAAGT | ASV_40 |
| Fungi | Ascomycota | Eurotiomycetes | Eurotiales | Aspergillaceae | Aspergillus | Aspergillus oryzae-flavus | ATGCGATAACT | ASV_43 |
| Fungi | Ascomycota | Eurotiomycetes | Eurotiales | Aspergillaceae | Penicillium | Penicillium sp. | ATGCGATACGT | ASV_46 |
| Fungi | Ascomycota | Saccharomycetes | Saccharomycetale | Debaryomycetace | Debaryomyces | Debaryomyces hansenii | GATAAGTAATA | ASV_47 |
| Fungi | Ascomycota | Saccharomycetes | Saccharomycetale | Saccharomycetac | Nakaseomyces | [Candida] glabrata | GATACGTAATG | ASV_48 |
| Fungi | Basidiomyco | Tremellomycetes | Cystofilobasidiale | Cystofilobasidiace | Cystofilobasidiu | Cystofilobasidium capitatum | TTGCGATAAGT | ASV_49 |
| Fungi | Ascomycota | Eurotiomycetes | Eurotiales | Aspergillaceae | Aspergillus | Aspergillus sp. | ATGCGATAATT | ASV_50 |
| Fungi | Ascomycota | Eurotiomycetes | Eurotiales | Aspergillaceae | Aspergillus | Aspergillus oryzae-flavus | ATGCGATAACT | ASV_57 |
| Fungi | Basidiomyco | Tremellomycetes | Cystofilobasidiale | Cystofilobasidiace | Cystofilobasidiu | Cystofilobasidium capitatum | TTGCGATAAGT | ASV_58 |
| Fungi | Ascomycota | Saccharomycetes | Saccharomycetale | Debaryomycetace | Millerozyma | Millerozyma farinosa | GATAAGTAATA | ASV_59 |
| Fungi | Ascomycota | Dothideomycetes | Pleosporales | Didymellaceae | Ascochyta | Ascochyta pisi | ATGCGATAAGT | ASV_61 |
| Fungi | Ascomycota | Saccharomycetes | Saccharomycetale | Phaffomycetacea | Candida | [Candida] sake | ATGCGATACGT | ASV_62 |
| Fungi | Mucoromyco | Mucoromycetes | Mucorales | Rhizopodaceae | Rhizopus | Rhizopus oryzae-arrhizus | GTGCGATAACT | ASV_75 |
| Fungi | Ascomycota | Saccharomycetes | Saccharomycetale | Saccharomycetac | Saccharomyces | Saccharomyces cerevisiae | ATGCGATACGT | ASV_76 |
| Fungi | Ascomycota | Dothideomycetes | Pleosporales | Didymellaceae | Stagonosporopsi | Stagonosporopsis cucurbitac | ATGCGATAAGT | ASV_81 |
| Fungi | Ascomycota | Eurotiomycetes | Eurotiales | Aspergillaceae | Aspergillus | Aspergillus penicillioides | ATGCGATACGT | ASV_89 |
| Fungi | Ascomycota | Eurotiomycetes | Eurotiales | Aspergillaceae | Aspergillus | Aspergillus oryzae-flavus | ATGCGATAACT | ASV_91 |
| Fungi | Ascomycota | Eurotiomycetes | Eurotiales | Aspergillaceae | Penicillium | Penicillium polonicum | ATGCGATACGT | ASV_93 |
| Fungi | Basidiomyco | Tremellomycetes | Cystofilobasidiale | Cystofilobasidiace | Cystofilobasidiu | Cystofilobasidium capitatum | TTGCGATAAGT | ASV_94 |
| Fungi | Ascomycota | Leotiomycetes | Helotiales | Sclerotiniaceae | Botrytis | Botrytis sp. | ATGCGATAAGT | ASV_97 |
| Fungi | Ascomycota | Eurotiomycetes | Eurotiales | Aspergillaceae | Aspergillus | Aspergillus oryzae-flavus | ATGCGATAACT | ASV_104 |
| Fungi | Basidiomyco | Tremellomycetes | Holtermanniales | Cryptococcaceae | Holtermanniella | Holtermanniella wattica | ATGCGATAAGT | ASV_105 |
| Fungi | Ascomycota | Saccharomycetes | Saccharomycetale | Phaffomycetacea | Wickerhamomyc | Wickerhamomyces anomalu | ATGCGATACGT | ASV_108 |
| Fungi | Ascomycota | Saccharomycetes | Saccharomycetale | Saccharomycetac | Starmerella | Starmerella sp. | GCGCGATAGGT | ASV_134 |
| Fungi | Ascomycota | Eurotiomycetes | Eurotiales | Aspergillaceae | Aspergillus | Aspergillus oryzae-flavus | ATGCGATAACT | ASV_137 |
| Fungi | Ascomycota | Saccharomycetes | Saccharomycetale | Phaffomycetacea | Wickerhamomyc | Wickerhamomyces anomalu | GATACGTATTGT | ASV_140 |
| Fungi | Basidiomyco | Microbotryomyc | Sporidiobolales | Sporidiobolaceae | Curvibasidium | Curvibasidium pallidicorallin | ATGTGATAAGT | ASV_144 |
| Fungi | Ascomycota | Saccharomycetes | Saccharomycetale | Saccharomycetac | Zygosaccharomyc | Zygosaccharomyces parabail | ATGCGATACGT | ASV_146 |
| Fungi | Ascomycota | Saccharomycetes | Saccharomycetale | Pichiaceae | Issatchenkia | Issatchenkia orientalis | ATGCGATACCT | ASV_175 |
| Fungi | Mucoromyco | Mucoromycetes | Mucorales | Rhizopodaceae | Rhizopus | Rhizopus microsporus | GTGCGATAACT | ASV_204 |

| ASV | F.1a | F.1b | F.1c | F1.3a | F1.3b | F1.3c | F2.3a | F2.3b | F2.3c | F3.3a | F3.3b | F3.3c |
|---|---|---|---|---|---|---|---|---|---|---|---|---|
| Fermentation | start | start | start | end | end | end | end | end | end | end | end | end |
| Technical replicate | a | b | c | a | b | c | a | b | c | a | b | c |
| Description | fava bean start | fava bean start | fava bean start | fava bean end_rep1 | fava bean end_rep1 | fava bean end_rep1 | fava bean end_rep2 | fava bean end_rep2 | fava bean end_rep2 | fava bean end_rep3 | fava bean end_rep3 | fava bean end_rep3 |
| ASV_1 | 8119 | 10652 | 9186 | 7244 | 10018 | 6995 | 24820 | 21836 | 21549 | 30587 | 28308 | 21702 |
| ASV_2 | 0 | 0 | 0 | 436 | 356 | 745 | 1027 | 1327 | 1540 | 247 | 262 | 0 |
| ASV_4 | 0 | 0 | 0 | 0 | 0 | 0 | 0 | 0 | 0 | 0 | 0 | 0 |
| ASV_12 | 0 | 0 | 0 | 0 | 0 | 0 | 0 | 0 | 0 | 0 | 0 | 0 |
| ASV_13 | 0 | 0 | 0 | 0 | 0 | 0 | 0 | 0 | 0 | 0 | 0 | 0 |
| ASV_14 | 0 | 0 | 0 | 0 | 0 | 0 | 59 | 59 | 0 | 0 | 0 | 21 |
| ASV_15 | 0 | 0 | 0 | 0 | 0 | 0 | 0 | 0 | 0 | 0 | 0 | 0 |
| ASV_16 | 0 | 0 | 0 | 0 | 0 | 0 | 0 | 0 | 0 | 0 | 0 | 0 |
| ASV_17 | 0 | 0 | 0 | 0 | 0 | 0 | 56 | 74 | 0 | 0 | 27 | 35 |
| ASV_18 | 0 | 0 | 0 | 0 | 0 | 0 | 0 | 0 | 0 | 0 | 0 | 0 |
| ASV_20 | 0 | 0 | 0 | 0 | 0 | 0 | 0 | 0 | 0 | 0 | 0 | 0 |
| ASV_21 | 0 | 0 | 0 | 0 | 0 | 0 | 0 | 0 | 0 | 0 | 0 | 0 |
| ASV_22 | 0 | 0 | 0 | 0 | 0 | 0 | 0 | 0 | 0 | 0 | 0 | 0 |
| ASV_23 | 0 | 0 | 0 | 0 | 0 | 0 | 0 | 0 | 0 | 0 | 0 | 0 |
| ASV_25 | 0 | 0 | 0 | 0 | 0 | 0 | 0 | 0 | 0 | 0 | 0 | 0 |
| ASV_26 | 0 | 0 | 0 | 0 | 0 | 0 | 0 | 0 | 0 | 0 | 0 | 0 |
| ASV_27 | 0 | 0 | 0 | 0 | 0 | 0 | 0 | 0 | 0 | 0 | 0 | 0 |
| ASV_28 | 0 | 0 | 0 | 31 | 15 | 17 | 11 | 17 | 19 | 24 | 29 | 12 |
| ASV_29 | 0 | 0 | 0 | 0 | 0 | 0 | 0 | 0 | 0 | 0 | 0 | 0 |
| ASV_30 | 0 | 0 | 0 | 0 | 0 | 0 | 0 | 0 | 0 | 0 | 0 | 0 |
| ASV_31 | 0 | 0 | 0 | 0 | 0 | 0 | 0 | 0 | 0 | 0 | 0 | 0 |
| ASV_34 | 0 | 0 | 0 | 0 | 0 | 0 | 0 | 0 | 0 | 0 | 0 | 0 |
| ASV_35 | 0 | 0 | 0 | 0 | 0 | 0 | 0 | 0 | 0 | 0 | 0 | 0 |
| ASV_37 | 0 | 0 | 0 | 0 | 0 | 0 | 0 | 0 | 0 | 0 | 0 | 0 |
| ASV_39 | 0 | 0 | 0 | 0 | 0 | 0 | 0 | 0 | 130 | 55 | 0 | 0 |
| ASV_40 | 0 | 0 | 0 | 0 | 0 | 0 | 0 | 0 | 0 | 0 | 0 | 0 |
| ASV_43 | 0 | 0 | 0 | 0 | 0 | 0 | 0 | 0 | 0 | 0 | 0 | 0 |
| ASV_46 | 0 | 0 | 0 | 0 | 0 | 0 | 0 | 0 | 0 | 0 | 0 | 0 |
| ASV_47 | 0 | 0 | 0 | 0 | 0 | 0 | 0 | 0 | 0 | 8 | 0 | 0 |
| ASV_48 | 0 | 0 | 0 | 0 | 0 | 0 | 0 | 0 | 83 | 15 | 0 | 0 |
| ASV_49 | 0 | 0 | 0 | 0 | 0 | 0 | 0 | 0 | 0 | 0 | 0 | 0 |
| ASV_50 | 0 | 0 | 0 | 0 | 0 | 0 | 0 | 0 | 0 | 0 | 0 | 0 |
| ASV_57 | 0 | 0 | 0 | 0 | 0 | 0 | 0 | 0 | 0 | 0 | 0 | 0 |
| ASV_58 | 0 | 0 | 0 | 0 | 0 | 0 | 0 | 0 | 0 | 0 | 0 | 0 |
| ASV_59 | 0 | 0 | 0 | 0 | 0 | 0 | 0 | 0 | 0 | 0 | 0 | 350 |
| ASV_61 | 0 | 0 | 0 | 0 | 0 | 0 | 0 | 0 | 0 | 0 | 0 | 0 |
| ASV_62 | 0 | 0 | 0 | 0 | 0 | 0 | 0 | 0 | 0 | 0 | 0 | 0 |
| ASV_75 | 0 | 0 | 0 | 0 | 0 | 0 | 0 | 0 | 0 | 0 | 0 | 0 |
| ASV_76 | 0 | 0 | 0 | 0 | 0 | 0 | 0 | 0 | 0 | 0 | 0 | 0 |
| ASV_81 | 0 | 0 | 0 | 0 | 0 | 0 | 0 | 0 | 0 | 0 | 0 | 0 |
| ASV_89 | 0 | 0 | 0 | 0 | 0 | 0 | 0 | 0 | 0 | 0 | 0 | 0 |
| ASV_91 | 0 | 0 | 0 | 0 | 0 | 0 | 0 | 0 | 0 | 0 | 0 | 0 |
| ASV_93 | 0 | 0 | 0 | 0 | 0 | 0 | 0 | 0 | 0 | 0 | 0 | 0 |
| ASV_94 | 0 | 0 | 0 | 0 | 0 | 0 | 0 | 0 | 0 | 0 | 0 | 0 |
| ASV_97 | 0 | 0 | 0 | 0 | 0 | 0 | 0 | 0 | 0 | 0 | 0 | 0 |
| ASV_104 | 0 | 0 | 0 | 0 | 0 | 0 | 0 | 0 | 0 | 0 | 0 | 0 |
| ASV_105 | 0 | 0 | 0 | 0 | 0 | 0 | 0 | 0 | 0 | 0 | 0 | 0 |
| ASV_108 | 0 | 0 | 0 | 0 | 0 | 0 | 0 | 0 | 0 | 0 | 0 | 0 |
| ASV_134 | 0 | 0 | 0 | 0 | 0 | 0 | 0 | 0 | 0 | 0 | 0 | 0 |
| ASV_137 | 0 | 0 | 0 | 0 | 0 | 0 | 0 | 0 | 0 | 0 | 0 | 0 |
| ASV_140 | 0 | 0 | 0 | 0 | 0 | 0 | 0 | 0 | 0 | 0 | 0 | 0 |
| ASV_144 | 0 | 0 | 0 | 0 | 0 | 0 | 0 | 0 | 0 | 0 | 0 | 0 |
| ASV_146 | 0 | 0 | 0 | 0 | 0 | 0 | 0 | 0 | 0 | 0 | 0 | 0 |
| ASV_175 | 0 | 0 | 0 | 0 | 0 | 0 | 0 | 0 | 0 | 0 | 0 | 0 |
| ASV_204 | 0 | 0 | 0 | 0 | 0 | 0 | 0 | 0 | 0 | 0 | 0 | 0 |
| | 8119 | 10652 | 9186 | 7711 | 10389 | 7757 | 25973 | 23313 | 23321 | 30936 | 28626 | 22120 |

| ASV | NF.1a | NF.1b | NF.1c | NF1.3a | NF1.3b | NF1.3c | NF2.3a | NF2.3b | NF2.3c | NF3.3a | NF3.3b | NF3.3c |
|---|---|---|---|---|---|---|---|---|---|---|---|---|
| Fermentation | start | start | start | end | end | end | end | end | end | end | end | end |
| Technical replicate | a | b | c | a | b | c | a | b | c | a | b | c |
| Description | nixtamalised fava bean start | nixtamalised fava bean start | nixtamalised fava bean start | nixtamalised fava bean end_rep1 | nixtamalised fava bean end_rep1 | nixtamalised fava bean end_rep1 | nixtamalised fava bean end_rep2 | nixtamalised fava bean end_rep2 | nixtamalised fava bean end_rep2 | nixtamalised fava bean end_rep3 | nixtamalised fava bean end_rep3 | nixtamalised fava bean end_rep3 |
| ASV_1 | 16194 | 18530 | 12214 | 9925 | 12051 | 8562 | 17676 | 16144 | 14797 | 26694 | 26444 | 23527 |
| ASV_2 | 0 | 0 | 0 | 0 | 0 | 0 | 0 | 0 | 0 | 0 | 0 | 0 |
| ASV_4 | 0 | 0 | 0 | 0 | 0 | 0 | 0 | 0 | 0 | 0 | 0 | 0 |
| ASV_12 | 0 | 0 | 0 | 0 | 0 | 0 | 0 | 0 | 0 | 0 | 0 | 0 |
| ASV_13 | 0 | 0 | 0 | 0 | 0 | 0 | 0 | 0 | 0 | 0 | 0 | 0 |
| ASV_14 | 0 | 0 | 0 | 87 | 0 | 52 | 1319 | 792 | 1278 | 0 | 560 | 249 |
| ASV_15 | 0 | 0 | 0 | 0 | 0 | 0 | 0 | 0 | 0 | 0 | 0 | 0 |
| ASV_16 | 0 | 0 | 0 | 0 | 0 | 0 | 0 | 0 | 0 | 0 | 0 | 0 |
| ASV_17 | 0 | 0 | 0 | 162 | 0 | 102 | 2181 | 1432 | 2000 | 0 | 502 | 552 |
| ASV_18 | 0 | 0 | 0 | 0 | 0 | 0 | 0 | 0 | 0 | 0 | 0 | 0 |
| ASV_20 | 0 | 0 | 0 | 0 | 0 | 0 | 0 | 0 | 0 | 0 | 0 | 0 |
| ASV_21 | 0 | 0 | 0 | 0 | 0 | 0 | 0 | 0 | 0 | 0 | 0 | 0 |
| ASV_22 | 0 | 0 | 0 | 0 | 0 | 0 | 0 | 0 | 0 | 0 | 0 | 0 |
| ASV_23 | 0 | 0 | 0 | 0 | 0 | 0 | 0 | 0 | 0 | 0 | 0 | 0 |
| ASV_25 | 0 | 0 | 0 | 0 | 0 | 0 | 0 | 0 | 0 | 0 | 0 | 0 |
| ASV_26 | 0 | 0 | 0 | 0 | 0 | 0 | 0 | 0 | 0 | 0 | 0 | 0 |
| ASV_27 | 0 | 0 | 0 | 0 | 0 | 0 | 0 | 0 | 0 | 0 | 0 | 0 |
| ASV_28 | 0 | 0 | 0 | 0 | 19 | 0 | 135 | 695 | 509 | 56 | 40 | 87 |
| ASV_29 | 0 | 0 | 0 | 0 | 0 | 0 | 0 | 0 | 0 | 0 | 0 | 0 |
| ASV_30 | 0 | 0 | 0 | 0 | 0 | 0 | 0 | 0 | 0 | 0 | 0 | 0 |
| ASV_31 | 0 | 0 | 0 | 0 | 0 | 0 | 0 | 0 | 0 | 0 | 0 | 0 |
| ASV_34 | 0 | 0 | 0 | 0 | 0 | 0 | 0 | 0 | 0 | 0 | 0 | 0 |
| ASV_35 | 0 | 0 | 0 | 0 | 0 | 0 | 0 | 0 | 0 | 0 | 0 | 0 |
| ASV_37 | 0 | 0 | 0 | 0 | 0 | 0 | 0 | 0 | 0 | 0 | 0 | 0 |
| ASV_39 | 0 | 0 | 0 | 0 | 76 | 0 | 0 | 0 | 0 | 1293 | 0 | 0 |
| ASV_40 | 0 | 0 | 0 | 0 | 0 | 0 | 0 | 0 | 0 | 0 | 0 | 0 |
| ASV_43 | 0 | 0 | 0 | 0 | 0 | 0 | 0 | 0 | 0 | 0 | 0 | 0 |
| ASV_46 | 0 | 0 | 0 | 0 | 0 | 0 | 0 | 0 | 0 | 0 | 0 | 0 |
| ASV_47 | 0 | 0 | 0 | 0 | 0 | 0 | 0 | 0 | 0 | 0 | 0 | 0 |
| ASV_48 | 0 | 0 | 0 | 0 | 39 | 0 | 0 | 0 | 0 | 653 | 0 | 0 |
| ASV_49 | 0 | 0 | 0 | 0 | 0 | 0 | 0 | 0 | 0 | 0 | 0 | 0 |
| ASV_50 | 0 | 0 | 0 | 0 | 0 | 0 | 0 | 0 | 0 | 0 | 0 | 0 |
| ASV_57 | 0 | 0 | 0 | 0 | 0 | 0 | 0 | 0 | 0 | 0 | 0 | 0 |
| ASV_58 | 0 | 0 | 0 | 0 | 0 | 0 | 0 | 0 | 0 | 0 | 0 | 0 |
| ASV_59 | 0 | 0 | 0 | 0 | 0 | 0 | 0 | 0 | 0 | 0 | 0 | 0 |
| ASV_61 | 0 | 0 | 0 | 0 | 0 | 0 | 0 | 0 | 0 | 0 | 0 | 0 |
| ASV_62 | 0 | 0 | 0 | 0 | 0 | 0 | 0 | 0 | 0 | 0 | 0 | 0 |
| ASV_75 | 0 | 0 | 0 | 0 | 0 | 0 | 0 | 0 | 0 | 0 | 0 | 0 |
| ASV_76 | 0 | 0 | 0 | 0 | 0 | 0 | 0 | 0 | 0 | 0 | 0 | 0 |
| ASV_81 | 0 | 0 | 0 | 0 | 0 | 0 | 0 | 0 | 0 | 0 | 0 | 0 |
| ASV_89 | 0 | 0 | 0 | 0 | 0 | 0 | 0 | 0 | 0 | 0 | 0 | 0 |
| ASV_91 | 0 | 0 | 0 | 0 | 0 | 0 | 9 | 5 | 0 | 0 | 0 | 0 |
| ASV_93 | 0 | 0 | 0 | 0 | 0 | 0 | 0 | 0 | 0 | 0 | 0 | 0 |
| ASV_94 | 0 | 0 | 0 | 0 | 0 | 0 | 0 | 0 | 0 | 0 | 0 | 0 |
| ASV_97 | 0 | 0 | 0 | 0 | 0 | 0 | 0 | 0 | 0 | 0 | 0 | 0 |
| ASV_104 | 0 | 0 | 0 | 0 | 0 | 0 | 0 | 0 | 0 | 0 | 0 | 0 |
| ASV_105 | 0 | 0 | 0 | 0 | 0 | 0 | 0 | 0 | 0 | 0 | 0 | 0 |
| ASV_108 | 0 | 0 | 0 | 0 | 0 | 0 | 0 | 0 | 0 | 0 | 0 | 0 |
| ASV_134 | 0 | 0 | 0 | 0 | 0 | 0 | 0 | 0 | 0 | 0 | 0 | 0 |
| ASV_137 | 0 | 0 | 0 | 0 | 0 | 0 | 0 | 0 | 0 | 0 | 0 | 0 |
| ASV_140 | 0 | 0 | 0 | 0 | 0 | 0 | 0 | 0 | 0 | 0 | 0 | 0 |
| ASV_144 | 0 | 0 | 0 | 0 | 0 | 0 | 0 | 0 | 0 | 0 | 0 | 0 |
| ASV_146 | 0 | 0 | 0 | 0 | 0 | 0 | 14 | 0 | 0 | 0 | 0 | 0 |
| ASV_175 | 0 | 0 | 0 | 0 | 0 | 0 | 0 | 0 | 0 | 0 | 0 | 0 |
| ASV_204 | 0 | 0 | 0 | 0 | 0 | 0 | 0 | 0 | 0 | 0 | 0 | 0 |
| | 16194 | 18530 | 12214 | 10174 | 12185 | 8716 | 21334 | 19068 | 18584 | 28696 | 27546 | 24415 |

| ASV | L.1a | L.1b | L.1c | L1.3a | L1.3b | L1.3c | L2.3a | L2.3b | L2.3c | L3.3a | L3.3b | L3.3c |
|---|---|---|---|---|---|---|---|---|---|---|---|---|
| Fermentation | start | start | start | end | end | end | end | end | end | end | end | end |
| Technical replicate | a | b | c | a | b | c | a | b | c | a | b | c |
| Description | Gotland lentil start | Gotland lentil start | Gotland lentil start | Gotland lentil end rep_1 | Gotland lentil end rep_1 | Gotland lentil end rep_1 | Gotland lentil end rep_2 | Gotland lentil end rep_2 | Gotland lentil end rep_2 | Gotland lentil end rep_3 | Gotland lentil end rep_3 | Gotland lentil end rep_3 |
| ASV_1 | 6071 | 10875 | 7126 | 9694 | 7979 | 27 | 35248 | 28775 | 28005 | 40202 | 31471 | 33396 |
| ASV_2 | 0 | 0 | 0 | 72 | 67 | 0 | 633 | 204 | 264 | 2571 | 1621 | 1605 |
| ASV_4 | 19 | 15 | 29 | 8 | 0 | 0 | 67 | 38 | 72 | 12 | 0 | 23 |
| ASV_12 | 0 | 0 | 0 | 0 | 0 | 0 | 0 | 0 | 0 | 0 | 0 | 0 |
| ASV_13 | 0 | 0 | 0 | 0 | 0 | 0 | 0 | 0 | 0 | 0 | 0 | 0 |
| ASV_14 | 0 | 0 | 0 | 0 | 0 | 0 | 0 | 0 | 0 | 0 | 0 | 0 |
| ASV_15 | 0 | 0 | 0 | 0 | 0 | 0 | 0 | 0 | 0 | 0 | 0 | 0 |
| ASV_16 | 0 | 0 | 0 | 0 | 0 | 0 | 0 | 0 | 0 | 0 | 0 | 0 |
| ASV_17 | 0 | 0 | 0 | 0 | 0 | 0 | 0 | 0 | 0 | 0 | 0 | 0 |
| ASV_18 | 0 | 0 | 0 | 0 | 0 | 0 | 0 | 0 | 0 | 0 | 0 | 0 |
| ASV_20 | 0 | 0 | 0 | 0 | 0 | 0 | 0 | 0 | 0 | 0 | 0 | 0 |
| ASV_21 | 0 | 0 | 0 | 0 | 0 | 0 | 0 | 0 | 0 | 0 | 0 | 0 |
| ASV_22 | 0 | 0 | 0 | 0 | 0 | 0 | 0 | 0 | 0 | 0 | 0 | 0 |
| ASV_23 | 0 | 0 | 0 | 0 | 0 | 0 | 0 | 0 | 0 | 0 | 0 | 0 |
| ASV_25 | 0 | 0 | 0 | 0 | 0 | 0 | 0 | 0 | 0 | 0 | 0 | 0 |
| ASV_26 | 0 | 0 | 0 | 0 | 0 | 0 | 0 | 0 | 0 | 0 | 0 | 0 |
| ASV_27 | 0 | 0 | 0 | 0 | 0 | 0 | 0 | 0 | 0 | 0 | 0 | 0 |
| ASV_28 | 0 | 0 | 0 | 78 | 134 | 0 | 294 | 111 | 41 | 311 | 421 | 151 |
| ASV_29 | 0 | 0 | 0 | 0 | 0 | 0 | 0 | 0 | 0 | 0 | 0 | 0 |
| ASV_30 | 0 | 0 | 0 | 0 | 0 | 0 | 0 | 0 | 0 | 0 | 0 | 0 |
| ASV_31 | 0 | 0 | 0 | 0 | 0 | 0 | 0 | 0 | 0 | 0 | 0 | 0 |
| ASV_34 | 0 | 0 | 0 | 0 | 0 | 0 | 0 | 0 | 0 | 0 | 0 | 0 |
| ASV_35 | 0 | 0 | 0 | 0 | 0 | 0 | 0 | 0 | 0 | 0 | 0 | 0 |
| ASV_37 | 0 | 0 | 0 | 0 | 0 | 0 | 0 | 0 | 0 | 0 | 0 | 0 |
| ASV_39 | 0 | 0 | 0 | 0 | 0 | 0 | 0 | 0 | 0 | 0 | 0 | 0 |
| ASV_40 | 0 | 0 | 0 | 0 | 0 | 0 | 0 | 0 | 0 | 0 | 0 | 0 |
| ASV_43 | 0 | 0 | 0 | 0 | 0 | 0 | 0 | 0 | 0 | 0 | 0 | 0 |
| ASV_46 | 0 | 0 | 0 | 0 | 0 | 0 | 0 | 0 | 0 | 0 | 0 | 0 |
| ASV_47 | 0 | 0 | 0 | 0 | 0 | 0 | 0 | 0 | 0 | 0 | 0 | 0 |
| ASV_48 | 0 | 0 | 0 | 0 | 0 | 0 | 0 | 0 | 0 | 0 | 0 | 0 |
| ASV_49 | 0 | 0 | 0 | 0 | 0 | 0 | 0 | 0 | 0 | 0 | 0 | 0 |
| ASV_50 | 0 | 0 | 0 | 0 | 0 | 0 | 0 | 0 | 0 | 28 | 0 | 202 |
| ASV_57 | 0 | 0 | 0 | 0 | 0 | 0 | 0 | 0 | 0 | 0 | 0 | 0 |
| ASV_58 | 0 | 0 | 0 | 0 | 0 | 0 | 0 | 0 | 0 | 0 | 0 | 0 |
| ASV_59 | 0 | 0 | 0 | 0 | 0 | 0 | 0 | 0 | 0 | 0 | 0 | 0 |
| ASV_61 | 0 | 0 | 0 | 0 | 0 | 0 | 0 | 0 | 0 | 0 | 0 | 0 |
| ASV_62 | 0 | 0 | 0 | 0 | 0 | 0 | 0 | 0 | 0 | 0 | 0 | 0 |
| ASV_75 | 0 | 0 | 0 | 0 | 0 | 0 | 0 | 0 | 0 | 0 | 0 | 0 |
| ASV_76 | 0 | 0 | 0 | 0 | 0 | 0 | 0 | 0 | 0 | 0 | 0 | 0 |
| ASV_81 | 0 | 0 | 0 | 0 | 0 | 0 | 0 | 0 | 0 | 0 | 0 | 0 |
| ASV_89 | 0 | 0 | 0 | 0 | 0 | 0 | 0 | 0 | 0 | 0 | 0 | 0 |
| ASV_91 | 0 | 0 | 0 | 0 | 0 | 0 | 14 | 14 | 2 | 8 | 0 | 0 |
| ASV_93 | 0 | 0 | 0 | 0 | 0 | 0 | 0 | 0 | 0 | 0 | 0 | 0 |
| ASV_94 | 0 | 0 | 0 | 0 | 0 | 0 | 0 | 0 | 0 | 0 | 0 | 0 |
| ASV_97 | 0 | 0 | 0 | 0 | 0 | 0 | 0 | 0 | 0 | 0 | 0 | 0 |
| ASV_104 | 0 | 0 | 0 | 0 | 0 | 0 | 0 | 0 | 0 | 0 | 0 | 0 |
| ASV_105 | 0 | 0 | 0 | 0 | 0 | 0 | 0 | 0 | 0 | 0 | 0 | 0 |
| ASV_108 | 0 | 0 | 0 | 0 | 0 | 0 | 0 | 0 | 0 | 0 | 0 | 0 |
| ASV_134 | 0 | 0 | 0 | 0 | 0 | 0 | 0 | 0 | 0 | 0 | 0 | 0 |
| ASV_137 | 0 | 0 | 0 | 0 | 0 | 0 | 0 | 0 | 0 | 0 | 0 | 0 |
| ASV_140 | 0 | 0 | 0 | 0 | 0 | 0 | 0 | 0 | 0 | 0 | 0 | 0 |
| ASV_144 | 0 | 0 | 0 | 0 | 0 | 0 | 0 | 0 | 0 | 0 | 0 | 0 |
| ASV_146 | 0 | 0 | 0 | 0 | 0 | 0 | 0 | 0 | 0 | 0 | 0 | 0 |
| ASV_175 | 0 | 0 | 0 | 0 | 0 | 0 | 0 | 0 | 0 | 0 | 0 | 0 |
| ASV_204 | 0 | 0 | 0 | 0 | 0 | 0 | 0 | 0 | 0 | 0 | 0 | 0 |
| | 6090 | 10890 | 7155 | 9852 | 8180 | 27 | 36256 | 29142 | 28384 | 43132 | 33513 | 35377 |

| ASV | NL.1a | NL.1b | NL.1c | NL1.3a | NL1.3b | NL1.3c | NL2.3a | NL2.3b | NL2.3c | NL3.3a | NL3.3b | NL3.3c |
|---|---|---|---|---|---|---|---|---|---|---|---|---|
| Fermentation | start | start | start | end | end | end | end | end | end | end | end | end |
| Technical replicate | a | b | c | a | b | c | a | b | c | a | b | c |
| Description | nixtamalised Gotland lentil start | nixtamalised Gotland lentil start | nixtamalised Gotland lentil start | nixtamalised Gotland lentil end_rep1 | nixtamalised Gotland lentil end_rep1 | nixtamalised Gotland lentil end_rep1 | nixtamalised Gotland lentil end_rep2 | nixtamalised Gotland lentil end_rep2 | nixtamalised Gotland lentil end_rep2 | nixtamalised Gotland lentil end_rep3 | nixtamalised Gotland lentil end_rep3 | nixtamalised Gotland lentil end_rep3 |
| ASV_1 | 42889 | 67706 | 56538 | 11589 | 18859 | 15164 | 19600 | 19324 | 19747 | 1E+05 | 98370 | 89240 |
| ASV_2 | 0 | 0 | 0 | 0 | 0 | 0 | 0 | 0 | 0 | 0 | 0 | 0 |
| ASV_4 | 0 | 0 | 0 | 0 | 0 | 0 | 0 | 0 | 0 | 0 | 0 | 0 |
| ASV_12 | 0 | 0 | 0 | 653 | 793 | 563 | 11106 | 10294 | 10077 | 0 | 0 | 0 |
| ASV_13 | 0 | 0 | 0 | 0 | 0 | 0 | 0 | 0 | 0 | 0 | 0 | 0 |
| ASV_14 | 0 | 0 | 0 | 1159 | 1595 | 1271 | 241 | 137 | 213 | 768 | 394 | 547 |
| ASV_15 | 0 | 0 | 0 | 0 | 0 | 0 | 0 | 0 | 0 | 0 | 0 | 0 |
| ASV_16 | 0 | 0 | 0 | 0 | 0 | 0 | 0 | 0 | 0 | 0 | 0 | 0 |
| ASV_17 | 0 | 0 | 0 | 1638 | 2594 | 1895 | 396 | 239 | 400 | 1283 | 487 | 686 |
| ASV_18 | 0 | 0 | 0 | 0 | 0 | 0 | 0 | 0 | 0 | 0 | 0 | 0 |
| ASV_20 | 0 | 0 | 0 | 0 | 0 | 0 | 0 | 0 | 0 | 0 | 0 | 0 |
| ASV_21 | 0 | 0 | 0 | 0 | 0 | 0 | 6200 | 4618 | 4209 | 0 | 0 | 0 |
| ASV_22 | 0 | 0 | 0 | 0 | 0 | 0 | 3946 | 3241 | 2347 | 0 | 0 | 0 |
| ASV_23 | 0 | 0 | 0 | 1379 | 1891 | 1866 | 0 | 0 | 0 | 0 | 0 | 0 |
| ASV_25 | 0 | 0 | 0 | 398 | 334 | 353 | 0 | 0 | 0 | 0 | 0 | 0 |
| ASV_26 | 0 | 0 | 0 | 0 | 0 | 0 | 0 | 0 | 0 | 0 | 0 | 0 |
| ASV_27 | 0 | 0 | 0 | 0 | 0 | 0 | 0 | 0 | 0 | 0 | 0 | 0 |
| ASV_28 | 0 | 0 | 0 | 36 | 19 | 22 | 0 | 0 | 33 | 0 | 57 | 0 |
| ASV_29 | 0 | 0 | 0 | 0 | 0 | 0 | 0 | 0 | 0 | 0 | 0 | 0 |
| ASV_30 | 0 | 0 | 0 | 0 | 0 | 0 | 0 | 0 | 0 | 0 | 0 | 0 |
| ASV_31 | 0 | 0 | 0 | 0 | 0 | 0 | 0 | 0 | 0 | 0 | 0 | 0 |
| ASV_34 | 0 | 0 | 0 | 0 | 0 | 0 | 0 | 0 | 0 | 0 | 0 | 0 |
| ASV_35 | 0 | 0 | 0 | 0 | 0 | 0 | 0 | 0 | 0 | 0 | 0 | 0 |
| ASV_37 | 0 | 0 | 0 | 0 | 0 | 0 | 0 | 0 | 0 | 0 | 0 | 0 |
| ASV_39 | 0 | 0 | 0 | 0 | 0 | 0 | 0 | 0 | 0 | 0 | 0 | 0 |
| ASV_40 | 0 | 0 | 0 | 0 | 0 | 0 | 0 | 0 | 0 | 0 | 0 | 0 |
| ASV_43 | 0 | 0 | 0 | 0 | 0 | 0 | 88 | 96 | 99 | 0 | 0 | 0 |
| ASV_46 | 0 | 0 | 0 | 0 | 0 | 0 | 0 | 0 | 0 | 0 | 0 | 0 |
| ASV_47 | 0 | 0 | 0 | 0 | 0 | 0 | 0 | 0 | 0 | 0 | 0 | 0 |
| ASV_48 | 0 | 0 | 0 | 0 | 0 | 0 | 0 | 0 | 0 | 0 | 0 | 0 |
| ASV_49 | 0 | 0 | 0 | 0 | 0 | 0 | 0 | 0 | 0 | 0 | 0 | 0 |
| ASV_50 | 0 | 0 | 0 | 0 | 0 | 0 | 0 | 0 | 0 | 0 | 0 | 0 |
| ASV_57 | 0 | 0 | 0 | 0 | 0 | 0 | 146 | 147 | 152 | 0 | 0 | 0 |
| ASV_58 | 0 | 0 | 0 | 0 | 0 | 0 | 0 | 0 | 0 | 0 | 0 | 0 |
| ASV_59 | 0 | 0 | 0 | 0 | 0 | 0 | 0 | 0 | 0 | 0 | 0 | 0 |
| ASV_61 | 0 | 0 | 0 | 0 | 0 | 0 | 0 | 0 | 0 | 0 | 0 | 0 |
| ASV_62 | 0 | 0 | 0 | 0 | 0 | 0 | 0 | 0 | 0 | 0 | 0 | 0 |
| ASV_75 | 0 | 0 | 0 | 0 | 0 | 21 | 0 | 0 | 0 | 0 | 0 | 0 |
| ASV_76 | 0 | 0 | 0 | 0 | 0 | 0 | 85 | 54 | 70 | 0 | 0 | 0 |
| ASV_81 | 0 | 0 | 0 | 0 | 0 | 0 | 0 | 0 | 0 | 0 | 0 | 0 |
| ASV_89 | 0 | 0 | 0 | 0 | 0 | 0 | 0 | 0 | 0 | 0 | 0 | 0 |
| ASV_91 | 0 | 0 | 0 | 0 | 0 | 0 | 0 | 0 | 0 | 0 | 0 | 0 |
| ASV_93 | 0 | 0 | 0 | 0 | 0 | 0 | 0 | 0 | 0 | 0 | 0 | 0 |
| ASV_94 | 0 | 0 | 0 | 0 | 0 | 0 | 0 | 0 | 0 | 0 | 0 | 0 |
| ASV_97 | 0 | 0 | 0 | 0 | 0 | 0 | 28 | 12 | 0 | 0 | 63 | 0 |
| ASV_104 | 0 | 0 | 0 | 0 | 0 | 0 | 33 | 33 | 30 | 0 | 0 | 0 |
| ASV_105 | 0 | 0 | 0 | 0 | 0 | 0 | 0 | 0 | 0 | 0 | 0 | 0 |
| ASV_108 | 0 | 0 | 0 | 0 | 0 | 0 | 0 | 0 | 0 | 0 | 0 | 0 |
| ASV_134 | 0 | 0 | 0 | 0 | 0 | 0 | 0 | 0 | 0 | 0 | 0 | 0 |
| ASV_137 | 0 | 0 | 0 | 0 | 0 | 0 | 0 | 0 | 0 | 0 | 0 | 0 |
| ASV_140 | 0 | 0 | 0 | 0 | 0 | 0 | 0 | 0 | 0 | 0 | 0 | 0 |
| ASV_144 | 0 | 0 | 0 | 0 | 0 | 0 | 0 | 0 | 0 | 0 | 0 | 0 |
| ASV_146 | 0 | 0 | 0 | 0 | 0 | 0 | 0 | 0 | 0 | 0 | 16 | 0 |
| ASV_175 | 0 | 0 | 0 | 0 | 0 | 0 | 0 | 0 | 0 | 0 | 0 | 0 |
| ASV_204 | 0 | 0 | 0 | 0 | 0 | 0 | 0 | 0 | 0 | 0 | 0 | 0 |
| | 42889 | 67706 | 56538 | 16852 | 26085 | 21155 | 41869 | 38195 | 37377 | 1E+05 | 99387 | 90473 |

| ASV | P.1a | P.1b | P.1c | P1.3a | P1.3b | P1.3c | P2.3a | P2.3b | P2.3c | P3.3a | P3.3b | P3.3c |
|---|---|---|---|---|---|---|---|---|---|---|---|---|
| Fermentation | start | start | start | end | end | end | end | end | end | end | end | end |
| Technical replicate | a | b | c | a | b | c | a | b | c | a | b | c |
| Description | yellow pea start | yellow pea start | yellow pea start | yellow pea end_rep1 | yellow pea end_rep1 | yellow pea end_rep1 | yellow pea end_rep2 | yellow pea end_rep2 | yellow pea end_rep2 | yellow pea end_rep3 | yellow pea end_rep3 | yellow pea end_rep3 |
| ASV_1 | 12631 | 24496 | 16345 | 1780 | 2398 | 1272 | 15046 | 15677 | 11774 | 42503 | 42138 | 47876 |
| ASV_2 | 0 | 0 | 0 | 11690 | 14060 | 6636 | 30015 | 29067 | 20206 | 2990 | 2492 | 3062 |
| ASV_4 | 0 | 0 | 0 | 0 | 0 | 0 | 0 | 0 | 0 | 0 | 0 | 0 |
| ASV_12 | 0 | 0 | 0 | 0 | 0 | 0 | 0 | 0 | 0 | 0 | 0 | 0 |
| ASV_13 | 0 | 0 | 0 | 0 | 0 | 0 | 0 | 0 | 0 | 0 | 0 | 0 |
| ASV_14 | 0 | 0 | 0 | 0 | 0 | 0 | 29 | 35 | 69 | 0 | 0 | 0 |
| ASV_15 | 193 | 353 | 237 | 0 | 0 | 0 | 212 | 225 | 0 | 561 | 539 | 709 |
| ASV_16 | 0 | 0 | 0 | 0 | 0 | 0 | 0 | 0 | 0 | 0 | 0 | 0 |
| ASV_17 | 0 | 0 | 0 | 0 | 0 | 0 | 0 | 27 | 49 | 0 | 0 | 0 |
| ASV_18 | 0 | 0 | 0 | 0 | 0 | 0 | 0 | 0 | 0 | 0 | 0 | 0 |
| ASV_20 | 0 | 0 | 0 | 0 | 0 | 0 | 0 | 0 | 0 | 0 | 0 | 0 |
| ASV_21 | 0 | 0 | 0 | 0 | 0 | 0 | 0 | 0 | 0 | 0 | 0 | 0 |
| ASV_22 | 0 | 0 | 0 | 0 | 0 | 0 | 0 | 0 | 0 | 0 | 0 | 0 |
| ASV_23 | 0 | 0 | 0 | 0 | 0 | 0 | 0 | 0 | 0 | 0 | 0 | 0 |
| ASV_25 | 0 | 0 | 0 | 0 | 0 | 0 | 0 | 0 | 0 | 0 | 0 | 0 |
| ASV_26 | 0 | 0 | 0 | 0 | 0 | 0 | 0 | 0 | 0 | 0 | 0 | 0 |
| ASV_27 | 0 | 0 | 0 | 0 | 0 | 0 | 0 | 0 | 0 | 0 | 0 | 0 |
| ASV_28 | 0 | 0 | 0 | 0 | 0 | 0 | 52 | 16 | 17 | 269 | 20 | 241 |
| ASV_29 | 0 | 0 | 0 | 163 | 214 | 0 | 425 | 399 | 334 | 0 | 0 | 0 |
| ASV_30 | 0 | 0 | 0 | 0 | 0 | 0 | 0 | 0 | 0 | 0 | 0 | 0 |
| ASV_31 | 0 | 0 | 0 | 0 | 0 | 0 | 0 | 0 | 0 | 0 | 0 | 0 |
| ASV_34 | 0 | 0 | 0 | 0 | 0 | 0 | 0 | 0 | 0 | 0 | 0 | 0 |
| ASV_35 | 0 | 0 | 0 | 0 | 0 | 0 | 0 | 0 | 0 | 0 | 0 | 0 |
| ASV_37 | 0 | 0 | 0 | 0 | 0 | 0 | 0 | 0 | 0 | 0 | 0 | 0 |
| ASV_39 | 0 | 0 | 0 | 0 | 0 | 0 | 0 | 0 | 0 | 0 | 0 | 0 |
| ASV_40 | 0 | 0 | 0 | 0 | 0 | 0 | 0 | 0 | 0 | 0 | 0 | 0 |
| ASV_43 | 0 | 0 | 0 | 0 | 0 | 0 | 0 | 0 | 0 | 0 | 0 | 0 |
| ASV_46 | 0 | 0 | 0 | 0 | 0 | 0 | 0 | 0 | 0 | 0 | 0 | 0 |
| ASV_47 | 0 | 0 | 0 | 0 | 0 | 0 | 0 | 0 | 0 | 28 | 0 | 0 |
| ASV_48 | 0 | 0 | 0 | 0 | 0 | 0 | 0 | 0 | 0 | 0 | 0 | 0 |
| ASV_49 | 0 | 0 | 0 | 0 | 0 | 0 | 0 | 0 | 0 | 0 | 0 | 0 |
| ASV_50 | 0 | 0 | 0 | 0 | 0 | 0 | 0 | 0 | 0 | 0 | 0 | 0 |
| ASV_57 | 0 | 0 | 0 | 0 | 0 | 0 | 0 | 0 | 0 | 0 | 0 | 0 |
| ASV_58 | 0 | 0 | 0 | 0 | 0 | 0 | 0 | 0 | 0 | 0 | 0 | 0 |
| ASV_59 | 0 | 0 | 0 | 0 | 0 | 0 | 0 | 0 | 0 | 0 | 0 | 0 |
| ASV_61 | 0 | 0 | 0 | 0 | 0 | 0 | 0 | 0 | 0 | 0 | 0 | 0 |
| ASV_62 | 0 | 0 | 0 | 0 | 0 | 0 | 0 | 0 | 0 | 0 | 0 | 0 |
| ASV_75 | 0 | 0 | 0 | 0 | 0 | 0 | 0 | 0 | 0 | 0 | 0 | 0 |
| ASV_76 | 0 | 0 | 0 | 0 | 0 | 0 | 0 | 0 | 0 | 0 | 0 | 0 |
| ASV_81 | 0 | 0 | 0 | 0 | 0 | 0 | 0 | 0 | 0 | 0 | 0 | 0 |
| ASV_89 | 0 | 0 | 0 | 0 | 0 | 0 | 0 | 0 | 0 | 0 | 0 | 0 |
| ASV_91 | 0 | 0 | 0 | 0 | 0 | 0 | 0 | 0 | 0 | 0 | 0 | 0 |
| ASV_93 | 0 | 0 | 0 | 0 | 0 | 0 | 0 | 0 | 0 | 0 | 0 | 0 |
| ASV_94 | 0 | 0 | 0 | 0 | 0 | 0 | 0 | 0 | 0 | 0 | 0 | 0 |
| ASV_97 | 0 | 0 | 0 | 0 | 0 | 0 | 0 | 0 | 0 | 0 | 0 | 0 |
| ASV_104 | 0 | 0 | 0 | 0 | 0 | 0 | 0 | 0 | 0 | 0 | 0 | 0 |
| ASV_105 | 0 | 0 | 0 | 0 | 0 | 0 | 0 | 0 | 0 | 0 | 0 | 0 |
| ASV_108 | 0 | 0 | 0 | 0 | 0 | 0 | 0 | 0 | 0 | 0 | 0 | 0 |
| ASV_134 | 0 | 0 | 0 | 0 | 0 | 0 | 0 | 0 | 0 | 0 | 0 | 0 |
| ASV_137 | 0 | 0 | 0 | 0 | 0 | 0 | 0 | 0 | 0 | 0 | 0 | 0 |
| ASV_140 | 0 | 0 | 0 | 0 | 0 | 0 | 0 | 0 | 0 | 0 | 0 | 0 |
| ASV_144 | 0 | 0 | 0 | 0 | 0 | 0 | 0 | 0 | 0 | 0 | 0 | 0 |
| ASV_146 | 0 | 0 | 0 | 0 | 0 | 0 | 0 | 0 | 0 | 0 | 0 | 0 |
| ASV_175 | 0 | 0 | 0 | 0 | 0 | 0 | 0 | 0 | 0 | 0 | 0 | 0 |
| ASV_204 | 0 | 0 | 0 | 0 | 0 | 0 | 0 | 0 | 0 | 0 | 0 | 0 |
| | 12824 | 24849 | 16582 | 13633 | 16672 | 7908 | 45779 | 45446 | 32449 | 46351 | 45189 | 51888 |

| ASV | NP.1a | NP.1b | NP.1c | NP1.3a | NP1.3b | NP1.3c | NP2.3a | NP2.3b | NP2.3c | NP3.3a | NP3.3b | NP3.3c |
|---|---|---|---|---|---|---|---|---|---|---|---|---|
| Fermentation | start | start | start | end | end | end | end | end | end | end | end | end |
| Technical replicate | a | b | c | a | b | c | a | b | c | a | b | c |
| Description | nixtamalised yellow pea start | nixtamalised yellow pea start | nixtamalised yellow pea start | nixtamalised yellow pea end_rep1 | nixtamalised yellow pea end_rep1 | nixtamalised yellow pea end_rep1 | nixtamalised yellow pea end_rep2 | nixtamalised yellow pea end_rep2 | nixtamalised yellow pea end_rep2 | nixtamalised yellow pea end_rep3 | nixtamalised yellow pea end_rep3 | nixtamalised yellow pea end_rep3 |
| ASV_1 | 38702 | 65417 | 44905 | 26560 | 36395 | 26665 | 63611 | 57585 | 40064 | 84260 | 83716 | 69934 |
| ASV_2 | 0 | 0 | 0 | 0 | 0 | 0 | 0 | 0 | 0 | 0 | 0 | 0 |
| ASV_4 | 0 | 0 | 0 | 0 | 0 | 0 | 0 | 0 | 0 | 0 | 0 | 0 |
| ASV_12 | 0 | 0 | 0 | 0 | 0 | 0 | 0 | 0 | 0 | 0 | 0 | 0 |
| ASV_13 | 0 | 0 | 0 | 0 | 0 | 0 | 0 | 0 | 0 | 0 | 0 | 0 |
| ASV_14 | 0 | 0 | 0 | 108 | 165 | 157 | 339 | 48 | 259 | 259 | 224 | 357 |
| ASV_15 | 0 | 0 | 0 | 0 | 0 | 0 | 0 | 0 | 0 | 0 | 0 | 0 |
| ASV_16 | 0 | 0 | 0 | 0 | 0 | 0 | 0 | 0 | 0 | 0 | 0 | 0 |
| ASV_17 | 0 | 0 | 0 | 221 | 325 | 240 | 476 | 414 | 1113 | 446 | 327 | 363 |
| ASV_18 | 0 | 0 | 0 | 0 | 0 | 0 | 0 | 0 | 0 | 0 | 0 | 0 |
| ASV_20 | 0 | 0 | 0 | 0 | 0 | 0 | 0 | 0 | 0 | 0 | 0 | 0 |
| ASV_21 | 0 | 0 | 0 | 0 | 0 | 0 | 0 | 0 | 0 | 0 | 0 | 0 |
| ASV_22 | 0 | 0 | 0 | 0 | 0 | 0 | 0 | 0 | 0 | 0 | 0 | 0 |
| ASV_23 | 0 | 0 | 0 | 0 | 0 | 0 | 0 | 0 | 0 | 872 | 926 | 819 |
| ASV_25 | 0 | 0 | 0 | 0 | 0 | 0 | 0 | 0 | 0 | 2061 | 1721 | 1640 |
| ASV_26 | 0 | 0 | 0 | 0 | 0 | 0 | 0 | 0 | 0 | 0 | 0 | 0 |
| ASV_27 | 0 | 0 | 0 | 0 | 0 | 0 | 0 | 0 | 0 | 0 | 0 | 0 |
| ASV_28 | 0 | 0 | 0 | 17 | 73 | 93 | 41 | 0 | 0 | 41 | 48 | 28 |
| ASV_29 | 0 | 0 | 0 | 0 | 0 | 0 | 0 | 0 | 0 | 0 | 0 | 0 |
| ASV_30 | 0 | 0 | 0 | 0 | 0 | 0 | 0 | 0 | 0 | 0 | 0 | 0 |
| ASV_31 | 0 | 0 | 0 | 0 | 0 | 0 | 0 | 0 | 0 | 0 | 0 | 0 |
| ASV_34 | 0 | 0 | 0 | 0 | 0 | 0 | 0 | 0 | 0 | 0 | 0 | 0 |
| ASV_35 | 0 | 0 | 0 | 0 | 0 | 0 | 0 | 0 | 0 | 0 | 0 | 0 |
| ASV_37 | 0 | 0 | 0 | 0 | 0 | 0 | 0 | 0 | 0 | 0 | 0 | 0 |
| ASV_39 | 0 | 0 | 0 | 0 | 0 | 0 | 0 | 0 | 0 | 0 | 0 | 0 |
| ASV_40 | 0 | 0 | 0 | 0 | 0 | 0 | 0 | 0 | 0 | 0 | 0 | 0 |
| ASV_43 | 0 | 0 | 0 | 0 | 0 | 0 | 0 | 0 | 0 | 119 | 0 | 98 |
| ASV_46 | 0 | 0 | 0 | 0 | 0 | 0 | 0 | 0 | 0 | 0 | 0 | 0 |
| ASV_47 | 0 | 0 | 0 | 0 | 0 | 0 | 0 | 0 | 0 | 0 | 0 | 0 |
| ASV_48 | 0 | 0 | 0 | 0 | 0 | 0 | 0 | 0 | 0 | 0 | 0 | 0 |
| ASV_49 | 0 | 0 | 0 | 0 | 0 | 0 | 0 | 0 | 0 | 0 | 0 | 0 |
| ASV_50 | 0 | 0 | 0 | 0 | 0 | 0 | 0 | 0 | 0 | 0 | 17 | 0 |
| ASV_57 | 0 | 0 | 0 | 0 | 0 | 0 | 0 | 0 | 0 | 0 | 0 | 0 |
| ASV_58 | 0 | 0 | 0 | 0 | 0 | 0 | 0 | 0 | 0 | 0 | 0 | 0 |
| ASV_59 | 0 | 0 | 0 | 0 | 0 | 0 | 0 | 0 | 0 | 0 | 0 | 0 |
| ASV_61 | 0 | 0 | 0 | 0 | 0 | 0 | 269 | 0 | 0 | 0 | 0 | 69 |
| ASV_62 | 0 | 0 | 0 | 0 | 0 | 0 | 0 | 0 | 0 | 0 | 0 | 0 |
| ASV_75 | 0 | 0 | 0 | 0 | 0 | 33 | 0 | 0 | 0 | 0 | 0 | 0 |
| ASV_76 | 0 | 0 | 0 | 0 | 0 | 0 | 0 | 0 | 0 | 0 | 0 | 0 |
| ASV_81 | 0 | 0 | 0 | 0 | 0 | 0 | 0 | 0 | 0 | 0 | 0 | 0 |
| ASV_89 | 0 | 0 | 0 | 0 | 0 | 0 | 0 | 0 | 0 | 0 | 0 | 0 |
| ASV_91 | 0 | 0 | 0 | 0 | 0 | 0 | 21 | 11 | 6 | 0 | 0 | 0 |
| ASV_93 | 0 | 0 | 0 | 0 | 0 | 0 | 0 | 0 | 0 | 0 | 0 | 0 |
| ASV_94 | 0 | 0 | 0 | 0 | 0 | 0 | 0 | 0 | 0 | 0 | 0 | 0 |
| ASV_97 | 0 | 0 | 0 | 0 | 0 | 0 | 0 | 0 | 0 | 0 | 8 | 0 |
| ASV_104 | 0 | 0 | 0 | 0 | 0 | 0 | 0 | 0 | 0 | 0 | 0 | 0 |
| ASV_105 | 0 | 0 | 0 | 0 | 0 | 0 | 0 | 0 | 0 | 0 | 0 | 0 |
| ASV_108 | 0 | 0 | 0 | 0 | 0 | 0 | 0 | 0 | 0 | 0 | 0 | 0 |
| ASV_134 | 0 | 0 | 0 | 0 | 0 | 0 | 0 | 0 | 0 | 0 | 0 | 0 |
| ASV_137 | 0 | 0 | 0 | 0 | 0 | 0 | 0 | 0 | 0 | 0 | 0 | 0 |
| ASV_140 | 0 | 0 | 0 | 0 | 0 | 0 | 0 | 0 | 0 | 0 | 0 | 0 |
| ASV_144 | 0 | 0 | 0 | 0 | 0 | 0 | 0 | 0 | 0 | 0 | 0 | 0 |
| ASV_146 | 0 | 0 | 0 | 0 | 0 | 0 | 0 | 0 | 0 | 0 | 0 | 0 |
| ASV_175 | 0 | 0 | 0 | 0 | 0 | 0 | 0 | 0 | 0 | 0 | 0 | 0 |
| ASV_204 | 0 | 0 | 0 | 0 | 0 | 0 | 0 | 0 | 0 | 0 | 0 | 0 |
| | 38702 | 65417 | 44905 | 26906 | 36958 | 27188 | 64757 | 58058 | 41442 | 88058 | 86987 | 73308 |

| ASV | R.1a | R.1b | R.1c | R1.3a | R1.3b | R1.3c | R2.3a | R2.3b | R2.3c | R3.3a | R3.3b | R3.3c |
|---|---|---|---|---|---|---|---|---|---|---|---|---|
| Fermentation | start | start | start | end | end | end | end | end | end | end | end | end |
| Technical replicate | a | b | c | a | b | c | a | b | c | a | b | c |
| Description | rye bread start | rye bread start | rye bread start | rye bread end_rep1 | rye bread end_rep1 | rye bread end_rep1 | rye bread end_rep2 | rye bread end_rep2 | rye bread end_rep2 | rye bread end_rep3 | rye bread end_rep3 | rye bread end_rep3 |
| ASV_1 | 1E+05 | 2E+05 | 1E+05 | 38989 | 50517 | 32086 | 1E+05 | 1E+05 | 1E+05 | 1E+05 | 1E+05 | 1E+05 |
| ASV_2 | 0 | 0 | 0 | 0 | 0 | 0 | 0 | 0 | 0 | 0 | 0 | 0 |
| ASV_4 | 0 | 0 | 0 | 0 | 0 | 0 | 0 | 0 | 0 | 0 | 0 | 0 |
| ASV_12 | 0 | 0 | 0 | 0 | 0 | 0 | 3700 | 3261 | 3262 | 50 | 0 | 41 |
| ASV_13 | 0 | 0 | 0 | 0 | 0 | 0 | 0 | 0 | 0 | 0 | 0 | 0 |
| ASV_14 | 0 | 0 | 0 | 0 | 0 | 0 | 0 | 0 | 0 | 0 | 0 | 0 |
| ASV_15 | 1677 | 2390 | 1880 | 597 | 650 | 421 | 1647 | 1655 | 1553 | 1714 | 2078 | 1943 |
| ASV_16 | 0 | 0 | 0 | 0 | 0 | 0 | 0 | 0 | 0 | 0 | 0 | 0 |
| ASV_17 | 0 | 0 | 0 | 0 | 0 | 0 | 0 | 0 | 0 | 0 | 0 | 0 |
| ASV_18 | 0 | 0 | 0 | 0 | 0 | 0 | 0 | 0 | 0 | 0 | 0 | 0 |
| ASV_20 | 0 | 0 | 0 | 0 | 0 | 0 | 0 | 0 | 0 | 0 | 0 | 0 |
| ASV_21 | 0 | 0 | 0 | 0 | 0 | 0 | 0 | 0 | 0 | 0 | 0 | 0 |
| ASV_22 | 0 | 0 | 0 | 0 | 0 | 0 | 0 | 0 | 0 | 0 | 0 | 0 |
| ASV_23 | 0 | 0 | 0 | 18 | 43 | 0 | 87 | 34 | 115 | 109 | 0 | 0 |
| ASV_25 | 0 | 0 | 0 | 0 | 0 | 21 | 0 | 0 | 0 | 0 | 109 | 0 |
| ASV_26 | 0 | 0 | 0 | 0 | 0 | 0 | 0 | 0 | 0 | 0 | 0 | 0 |
| ASV_27 | 0 | 0 | 0 | 0 | 0 | 0 | 0 | 0 | 0 | 0 | 0 | 0 |
| ASV_28 | 0 | 0 | 0 | 15 | 25 | 43 | 16 | 20 | 18 | 37 | 22 | 24 |
| ASV_29 | 0 | 0 | 0 | 0 | 0 | 0 | 0 | 0 | 0 | 0 | 0 | 0 |
| ASV_30 | 0 | 0 | 0 | 0 | 0 | 0 | 0 | 0 | 0 | 0 | 0 | 0 |
| ASV_31 | 0 | 0 | 0 | 0 | 0 | 0 | 0 | 0 | 0 | 0 | 0 | 0 |
| ASV_34 | 0 | 0 | 0 | 0 | 0 | 0 | 0 | 0 | 0 | 0 | 0 | 0 |
| ASV_35 | 0 | 0 | 0 | 0 | 0 | 0 | 0 | 0 | 0 | 0 | 0 | 0 |
| ASV_37 | 0 | 0 | 0 | 0 | 0 | 0 | 0 | 0 | 0 | 0 | 0 | 0 |
| ASV_39 | 0 | 0 | 0 | 0 | 0 | 0 | 0 | 0 | 0 | 0 | 0 | 0 |
| ASV_40 | 0 | 0 | 0 | 0 | 0 | 0 | 0 | 0 | 0 | 0 | 0 | 0 |
| ASV_43 | 0 | 0 | 0 | 0 | 0 | 0 | 69 | 109 | 114 | 0 | 0 | 0 |
| ASV_46 | 0 | 0 | 0 | 0 | 0 | 0 | 0 | 0 | 0 | 0 | 0 | 0 |
| ASV_47 | 0 | 0 | 0 | 0 | 0 | 0 | 0 | 0 | 0 | 0 | 0 | 0 |
| ASV_48 | 0 | 0 | 0 | 0 | 0 | 0 | 0 | 0 | 0 | 0 | 0 | 0 |
| ASV_49 | 0 | 0 | 0 | 0 | 0 | 0 | 0 | 0 | 0 | 0 | 0 | 0 |
| ASV_50 | 0 | 0 | 0 | 203 | 195 | 117 | 0 | 0 | 0 | 0 | 0 | 0 |
| ASV_57 | 0 | 0 | 0 | 0 | 0 | 0 | 0 | 0 | 0 | 0 | 0 | 0 |
| ASV_58 | 0 | 0 | 0 | 0 | 0 | 0 | 0 | 0 | 0 | 0 | 0 | 0 |
| ASV_59 | 0 | 0 | 0 | 0 | 0 | 0 | 0 | 0 | 0 | 0 | 0 | 0 |
| ASV_61 | 0 | 0 | 0 | 0 | 0 | 0 | 0 | 0 | 0 | 0 | 0 | 0 |
| ASV_62 | 0 | 0 | 0 | 0 | 0 | 0 | 0 | 0 | 0 | 0 | 0 | 0 |
| ASV_75 | 0 | 0 | 0 | 0 | 0 | 0 | 0 | 0 | 0 | 0 | 0 | 0 |
| ASV_76 | 0 | 0 | 0 | 0 | 0 | 0 | 0 | 0 | 0 | 0 | 0 | 0 |
| ASV_81 | 0 | 0 | 0 | 0 | 0 | 0 | 0 | 0 | 0 | 0 | 0 | 0 |
| ASV_89 | 0 | 0 | 0 | 0 | 0 | 0 | 0 | 0 | 0 | 30 | 50 | 61 |
| ASV_91 | 0 | 0 | 0 | 0 | 0 | 0 | 21 | 28 | 0 | 0 | 0 | 0 |
| ASV_93 | 0 | 0 | 0 | 0 | 0 | 0 | 0 | 0 | 0 | 0 | 0 | 0 |
| ASV_94 | 0 | 0 | 0 | 0 | 0 | 0 | 0 | 0 | 0 | 0 | 0 | 0 |
| ASV_97 | 0 | 0 | 0 | 0 | 0 | 0 | 0 | 0 | 0 | 0 | 0 | 0 |
| ASV_104 | 0 | 0 | 0 | 0 | 0 | 0 | 0 | 0 | 0 | 0 | 0 | 0 |
| ASV_105 | 0 | 0 | 0 | 0 | 0 | 0 | 0 | 0 | 0 | 0 | 0 | 0 |
| ASV_108 | 0 | 0 | 0 | 0 | 0 | 0 | 0 | 0 | 0 | 0 | 0 | 0 |
| ASV_134 | 0 | 0 | 0 | 0 | 0 | 0 | 0 | 0 | 0 | 0 | 0 | 0 |
| ASV_137 | 0 | 0 | 0 | 0 | 0 | 0 | 20 | 20 | 0 | 0 | 0 | 0 |
| ASV_140 | 0 | 0 | 0 | 0 | 0 | 0 | 0 | 0 | 0 | 0 | 0 | 0 |
| ASV_144 | 0 | 0 | 0 | 0 | 0 | 0 | 0 | 0 | 0 | 0 | 0 | 0 |
| ASV_146 | 0 | 0 | 0 | 0 | 0 | 0 | 0 | 0 | 0 | 0 | 16 | 0 |
| ASV_175 | 0 | 0 | 0 | 0 | 0 | 0 | 0 | 0 | 0 | 0 | 0 | 0 |
| ASV_204 | 0 | 0 | 0 | 0 | 0 | 8 | 0 | 5 | 0 | 0 | 0 | 0 |
| | 1E+05 | 2E+05 | 1E+05 | 39822 | 51430 | 32696 | 1E+05 | 1E+05 | 1E+05 | 1E+05 | 1E+05 | 1E+05 |

| ASV | S.1a | S.1b | S.1c | S1.3a | S1.3b | S1.3c | S2.3a | S2.3b | S2.3c | S3.3a | S3.3b | S3.3c |
|---|---|---|---|---|---|---|---|---|---|---|---|---|
| Fermentation | start | start | start | end | end | end | end | end | end | end | end | end |
| Technical replicate | a | b | c | a | b | c | a | b | c | a | b | c |
| Description | soybean start | soybean start | soybean start | soybean end_rep1 | soybean end_rep1 | soybean end_rep1 | soybean end_rep2 | soybean end_rep2 | soybean end_rep2 | soybean end_rep3 | soybean end_rep3 | soybean end_rep3 |
| ASV_1 | 61958 | 79256 | 67196 | 41641 | 49905 | 33335 | 92582 | 89264 | 65890 | 38702 | 39541 | 33183 |
| ASV_2 | 0 | 15 | 0 | 18 | 0 | 28 | 4356 | 4234 | 5161 | 72365 | 72696 | 60572 |
| ASV_4 | 0 | 0 | 0 | 0 | 0 | 0 | 0 | 0 | 0 | 0 | 0 | 0 |
| ASV_12 | 0 | 0 | 0 | 0 | 0 | 0 | 0 | 0 | 0 | 0 | 0 | 0 |
| ASV_13 | 0 | 0 | 0 | 0 | 0 | 0 | 0 | 0 | 0 | 0 | 0 | 0 |
| ASV_14 | 0 | 0 | 0 | 0 | 0 | 0 | 23 | 0 | 19 | 0 | 0 | 0 |
| ASV_15 | 849 | 1024 | 875 | 494 | 634 | 411 | 1293 | 1164 | 823 | 530 | 567 | 450 |
| ASV_16 | 0 | 0 | 0 | 0 | 0 | 0 | 0 | 0 | 0 | 0 | 0 | 0 |
| ASV_17 | 0 | 0 | 0 | 0 | 0 | 0 | 0 | 0 | 0 | 0 | 0 | 0 |
| ASV_18 | 0 | 0 | 0 | 0 | 0 | 0 | 0 | 0 | 0 | 0 | 0 | 0 |
| ASV_20 | 0 | 0 | 0 | 0 | 0 | 0 | 0 | 0 | 0 | 0 | 0 | 0 |
| ASV_21 | 0 | 0 | 0 | 0 | 0 | 0 | 0 | 0 | 0 | 0 | 0 | 0 |
| ASV_22 | 0 | 0 | 0 | 0 | 0 | 0 | 0 | 0 | 0 | 0 | 0 | 0 |
| ASV_23 | 0 | 0 | 0 | 0 | 0 | 0 | 0 | 0 | 0 | 0 | 0 | 0 |
| ASV_25 | 0 | 0 | 0 | 0 | 0 | 0 | 0 | 0 | 0 | 0 | 0 | 0 |
| ASV_26 | 0 | 0 | 0 | 0 | 0 | 0 | 0 | 0 | 0 | 0 | 0 | 0 |
| ASV_27 | 0 | 0 | 0 | 0 | 0 | 0 | 0 | 0 | 0 | 0 | 0 | 0 |
| ASV_28 | 0 | 0 | 0 | 0 | 0 | 0 | 21 | 0 | 0 | 0 | 3 | 0 |
| ASV_29 | 0 | 0 | 0 | 0 | 0 | 0 | 0 | 0 | 0 | 1009 | 944 | 890 |
| ASV_30 | 0 | 0 | 0 | 0 | 0 | 0 | 0 | 0 | 0 | 0 | 0 | 0 |
| ASV_31 | 0 | 0 | 0 | 0 | 0 | 0 | 0 | 0 | 0 | 0 | 0 | 0 |
| ASV_34 | 0 | 0 | 0 | 10 | 0 | 0 | 0 | 15 | 0 | 21 | 0 | 18 |
| ASV_35 | 0 | 0 | 0 | 0 | 0 | 0 | 0 | 0 | 0 | 0 | 0 | 0 |
| ASV_37 | 0 | 0 | 0 | 0 | 0 | 0 | 0 | 0 | 0 | 0 | 0 | 0 |
| ASV_39 | 0 | 0 | 0 | 0 | 0 | 0 | 0 | 0 | 0 | 0 | 0 | 0 |
| ASV_40 | 0 | 0 | 0 | 0 | 0 | 0 | 0 | 0 | 0 | 0 | 0 | 0 |
| ASV_43 | 0 | 0 | 0 | 0 | 0 | 0 | 0 | 0 | 0 | 0 | 0 | 0 |
| ASV_46 | 0 | 0 | 0 | 0 | 0 | 0 | 0 | 0 | 0 | 0 | 0 | 0 |
| ASV_47 | 0 | 0 | 0 | 0 | 0 | 0 | 0 | 0 | 0 | 0 | 0 | 0 |
| ASV_48 | 0 | 0 | 0 | 0 | 0 | 0 | 0 | 0 | 0 | 0 | 0 | 0 |
| ASV_49 | 0 | 0 | 0 | 0 | 0 | 0 | 0 | 0 | 0 | 0 | 0 | 0 |
| ASV_50 | 0 | 0 | 0 | 0 | 0 | 0 | 0 | 0 | 0 | 0 | 0 | 0 |
| ASV_57 | 0 | 0 | 0 | 0 | 0 | 0 | 0 | 0 | 0 | 0 | 0 | 0 |
| ASV_58 | 0 | 0 | 0 | 0 | 0 | 0 | 0 | 0 | 0 | 0 | 0 | 0 |
| ASV_59 | 0 | 0 | 0 | 0 | 15 | 0 | 0 | 0 | 0 | 0 | 0 | 0 |
| ASV_61 | 0 | 0 | 0 | 0 | 0 | 0 | 0 | 0 | 0 | 0 | 0 | 0 |
| ASV_62 | 0 | 0 | 0 | 0 | 0 | 0 | 0 | 0 | 0 | 0 | 0 | 0 |
| ASV_75 | 0 | 0 | 0 | 0 | 0 | 0 | 0 | 0 | 0 | 0 | 0 | 0 |
| ASV_76 | 0 | 0 | 0 | 0 | 0 | 0 | 0 | 0 | 0 | 0 | 0 | 0 |
| ASV_81 | 0 | 0 | 0 | 0 | 0 | 0 | 0 | 0 | 0 | 0 | 0 | 0 |
| ASV_89 | 0 | 0 | 0 | 0 | 0 | 0 | 0 | 0 | 0 | 0 | 0 | 0 |
| ASV_91 | 0 | 0 | 0 | 0 | 0 | 0 | 0 | 0 | 0 | 0 | 0 | 0 |
| ASV_93 | 0 | 0 | 0 | 0 | 0 | 0 | 0 | 0 | 0 | 0 | 0 | 0 |
| ASV_94 | 0 | 0 | 0 | 0 | 0 | 0 | 0 | 0 | 0 | 0 | 0 | 0 |
| ASV_97 | 0 | 0 | 0 | 0 | 0 | 0 | 0 | 0 | 0 | 0 | 0 | 0 |
| ASV_104 | 0 | 0 | 0 | 0 | 0 | 0 | 0 | 0 | 0 | 0 | 0 | 0 |
| ASV_105 | 0 | 0 | 0 | 0 | 0 | 0 | 0 | 0 | 0 | 0 | 0 | 0 |
| ASV_108 | 0 | 0 | 0 | 0 | 0 | 0 | 0 | 0 | 0 | 0 | 0 | 0 |
| ASV_134 | 0 | 0 | 0 | 0 | 0 | 0 | 0 | 0 | 0 | 0 | 0 | 0 |
| ASV_137 | 0 | 0 | 0 | 0 | 0 | 0 | 0 | 0 | 0 | 0 | 0 | 0 |
| ASV_140 | 0 | 0 | 0 | 0 | 0 | 0 | 0 | 0 | 0 | 0 | 0 | 0 |
| ASV_144 | 0 | 0 | 0 | 0 | 0 | 0 | 0 | 0 | 0 | 0 | 0 | 0 |
| ASV_146 | 0 | 0 | 0 | 0 | 0 | 0 | 0 | 0 | 0 | 0 | 0 | 0 |
| ASV_175 | 0 | 11 | 3 | 0 | 11 | 0 | 0 | 0 | 0 | 0 | 0 | 0 |
| ASV_204 | 0 | 0 | 0 | 0 | 0 | 0 | 0 | 0 | 0 | 0 | 0 | 0 |
| | 62807 | 80306 | 68074 | 42163 | 50565 | 33774 | 98275 | 94677 | 71893 | 1E+05 | 1E+05 | 95113 |

| ASV | HB.a | HB.b | HB.c | MS.a | MS.b | MS.c | PB.a | PB.b | PB.c |
|---|---|---|---|---|---|---|---|---|---|
| Technical replicate | a | b | c | a | b | c | a | b | c |
| Description | INUA Habanero Barley Koji Miso | INUA Habanero Barley Koji Miso | INUA Habanero Barley Koji Miso | INUA Maitake Soy Miso | INUA Maitake Soy Miso | INUA Maitake Soy Miso | INUA Yellow Peaso w/ B koji | INUA Yellow Peaso w/ B koji | INUA Yellow Peaso w/ B koji |
| ASV_1 | 6482 | 3573 | 4675 | 2277 | 2367 | 4023 | 18816 | 12725 | 17139 |
| ASV_2 | 0 | 0 | 0 | 0 | 0 | 0 | 0 | 0 | 0 |
| ASV_4 | 63694 | 45877 | 62449 | 253 | 1420 | 0 | 0 | 0 | 0 |
| ASV_12 | 0 | 0 | 0 | 0 | 0 | 0 | 1091 | 729 | 822 |
| ASV_13 | 132 | 124 | 84 | 2860 | 2400 | 2289 | 0 | 0 | 0 |
| ASV_14 | 0 | 0 | 0 | 1118 | 825 | 1105 | 0 | 0 | 0 |
| ASV_15 | 0 | 0 | 0 | 0 | 0 | 0 | 259 | 163 | 224 |
| ASV_16 | 0 | 81 | 0 | 6244 | 6019 | 5498 | 562 | 678 | 911 |
| ASV_17 | 0 | 0 | 0 | 0 | 0 | 0 | 0 | 0 | 0 |
| ASV_18 | 0 | 0 | 0 | 18 | 0 | 13 | 0 | 0 | 0 |
| ASV_20 | 44 | 35 | 46 | 3467 | 3254 | 4230 | 486 | 778 | 0 |
| ASV_21 | 0 | 0 | 0 | 0 | 0 | 0 | 0 | 0 | 0 |
| ASV_22 | 0 | 0 | 0 | 0 | 0 | 0 | 0 | 0 | 0 |
| ASV_23 | 0 | 0 | 0 | 0 | 0 | 0 | 0 | 0 | 0 |
| ASV_25 | 0 | 0 | 0 | 0 | 0 | 0 | 0 | 0 | 0 |
| ASV_26 | 127 | 81 | 108 | 0 | 0 | 0 | 2148 | 1418 | 1664 |
| ASV_27 | 0 | 0 | 0 | 509 | 242 | 287 | 114 | 177 | 166 |
| ASV_28 | 0 | 0 | 8 | 0 | 0 | 0 | 0 | 0 | 5 |
| ASV_29 | 0 | 0 | 0 | 0 | 0 | 0 | 0 | 0 | 0 |
| ASV_30 | 0 | 0 | 0 | 736 | 454 | 1368 | 26 | 0 | 94 |
| ASV_31 | 0 | 0 | 0 | 908 | 896 | 858 | 0 | 0 | 0 |
| ASV_34 | 0 | 0 | 0 | 158 | 70 | 100 | 0 | 0 | 0 |
| ASV_35 | 0 | 0 | 49 | 135 | 82 | 116 | 0 | 0 | 0 |
| ASV_37 | 0 | 0 | 0 | 624 | 377 | 834 | 28 | 15 | 43 |
| ASV_39 | 0 | 0 | 0 | 0 | 0 | 0 | 0 | 0 | 0 |
| ASV_40 | 7 | 0 | 6 | 359 | 509 | 324 | 55 | 52 | 71 |
| ASV_43 | 0 | 0 | 0 | 0 | 32 | 0 | 0 | 0 | 0 |
| ASV_46 | 0 | 0 | 0 | 0 | 0 | 46 | 356 | 212 | 315 |
| ASV_47 | 354 | 0 | 295 | 0 | 0 | 0 | 0 | 0 | 0 |
| ASV_48 | 0 | 0 | 0 | 0 | 0 | 0 | 0 | 0 | 0 |
| ASV_49 | 0 | 0 | 0 | 59 | 0 | 585 | 51 | 0 | 27 |
| ASV_50 | 0 | 0 | 0 | 0 | 0 | 0 | 0 | 0 | 0 |
| ASV_57 | 0 | 0 | 0 | 0 | 0 | 0 | 0 | 0 | 0 |
| ASV_58 | 0 | 0 | 0 | 178 | 0 | 190 | 40 | 0 | 39 |
| ASV_59 | 0 | 0 | 0 | 0 | 0 | 0 | 10 | 0 | 0 |
| ASV_61 | 0 | 0 | 0 | 0 | 0 | 0 | 0 | 0 | 0 |
| ASV_62 | 0 | 0 | 0 | 105 | 88 | 141 | 0 | 0 | 0 |
| ASV_75 | 0 | 0 | 0 | 0 | 0 | 147 | 0 | 11 | 0 |
| ASV_76 | 0 | 0 | 0 | 0 | 0 | 0 | 0 | 0 | 0 |
| ASV_81 | 0 | 0 | 0 | 93 | 0 | 74 | 0 | 0 | 0 |
| ASV_89 | 0 | 0 | 0 | 0 | 0 | 0 | 0 | 0 | 0 |
| ASV_91 | 0 | 0 | 0 | 0 | 0 | 0 | 0 | 0 | 0 |
| ASV_93 | 0 | 0 | 0 | 45 | 86 | 0 | 0 | 0 | 0 |
| ASV_94 | 0 | 0 | 0 | 0 | 0 | 78 | 0 | 50 | 0 |
| ASV_97 | 0 | 0 | 0 | 0 | 0 | 0 | 0 | 0 | 0 |
| ASV_104 | 0 | 0 | 0 | 0 | 0 | 0 | 0 | 0 | 0 |
| ASV_105 | 0 | 0 | 0 | 0 | 78 | 0 | 0 | 12 | 0 |
| ASV_108 | 0 | 0 | 0 | 0 | 0 | 0 | 0 | 0 | 0 |
| ASV_134 | 0 | 0 | 0 | 0 | 0 | 0 | 0 | 0 | 0 |
| ASV_137 | 0 | 0 | 0 | 0 | 14 | 0 | 0 | 0 | 0 |
| ASV_140 | 0 | 0 | 0 | 0 | 0 | 0 | 0 | 0 | 0 |
| ASV_144 | 0 | 0 | 0 | 37 | 0 | 0 | 0 | 0 | 11 |
| ASV_146 | 0 | 0 | 0 | 0 | 0 | 0 | 0 | 0 | 0 |
| ASV_175 | 0 | 0 | 0 | 0 | 0 | 0 | 0 | 0 | 0 |
| ASV_204 | 0 | 0 | 0 | 0 | 0 | 0 | 0 | 0 | 0 |
| | 70840 | 49771 | 67720 | 20183 | 19213 | 22306 | 24042 | 17020 | 21531 |

| ASV | PR.a | PR.b | PR.c | SY.a | SY.b | SY.c | TS.a | TS.b | TS.c |
|---|---|---|---|---|---|---|---|---|---|
| Technical replicate | a | b | c | a | b | c | a | b | c |
| Description | INUA Pea-so w/ Rice Koji 21.7 | INUA Pea-so w/ Rice Koji 21.7 | INUA Pea-so w/ Rice Koji 21.7 | INUA Soy Miso w/ Yuzu Pulp | INUA Soy Miso w/ Yuzu Pulp | INUA Soy Miso w/ Yuzu Pulp | INUA Toasted Sesame Miso | INUA Toasted Sesame Miso | INUA Toasted Sesame Miso |
| ASV_1 | 25724 | 19106 | 31426 | 1637 | 2131 | 2564 | 4805 | 7519 | 6412 |
| ASV_2 | 0 | 0 | 0 | 0 | 0 | 0 | 0 | 0 | 0 |
| ASV_4 | 0 | 0 | 0 | 0 | 0 | 0 | 0 | 0 | 0 |
| ASV_12 | 0 | 0 | 0 | 0 | 0 | 0 | 0 | 0 | 0 |
| ASV_13 | 0 | 0 | 0 | 0 | 0 | 0 | 14948 | 10360 | 11921 |
| ASV_14 | 0 | 0 | 0 | 0 | 0 | 0 | 5655 | 6303 | 4964 |
| ASV_15 | 0 | 0 | 0 | 0 | 0 | 0 | 0 | 0 | 0 |
| ASV_16 | 0 | 0 | 0 | 128 | 120 | 173 | 1570 | 919 | 1296 |
| ASV_17 | 0 | 0 | 0 | 0 | 0 | 0 | 0 | 0 | 0 |
| ASV_18 | 0 | 0 | 0 | 0 | 0 | 0 | 5284 | 8421 | 6452 |
| ASV_20 | 4 | 0 | 10 | 271 | 271 | 440 | 0 | 1146 | 673 |
| ASV_21 | 0 | 0 | 0 | 0 | 0 | 0 | 0 | 0 | 0 |
| ASV_22 | 0 | 0 | 0 | 0 | 0 | 0 | 0 | 0 | 0 |
| ASV_23 | 0 | 0 | 0 | 0 | 0 | 0 | 0 | 0 | 0 |
| ASV_25 | 0 | 0 | 0 | 0 | 0 | 0 | 0 | 0 | 0 |
| ASV_26 | 0 | 0 | 0 | 0 | 0 | 0 | 0 | 0 | 0 |
| ASV_27 | 1237 | 841 | 1335 | 0 | 0 | 0 | 0 | 0 | 374 |
| ASV_28 | 0 | 0 | 0 | 0 | 0 | 0 | 0 | 0 | 0 |
| ASV_29 | 0 | 0 | 0 | 0 | 0 | 0 | 0 | 0 | 0 |
| ASV_30 | 0 | 0 | 0 | 15 | 11 | 0 | 0 | 120 | 0 |
| ASV_31 | 0 | 0 | 0 | 0 | 0 | 0 | 0 | 0 | 0 |
| ASV_34 | 0 | 0 | 0 | 0 | 0 | 0 | 770 | 583 | 563 |
| ASV_35 | 0 | 0 | 0 | 18 | 0 | 0 | 678 | 586 | 565 |
| ASV_37 | 0 | 0 | 0 | 0 | 0 | 0 | 0 | 0 | 0 |
| ASV_39 | 0 | 0 | 0 | 0 | 0 | 0 | 0 | 0 | 0 |
| ASV_40 | 0 | 0 | 0 | 0 | 19 | 0 | 0 | 50 | 31 |
| ASV_43 | 75 | 0 | 91 | 0 | 0 | 0 | 0 | 0 | 0 |
| ASV_46 | 0 | 0 | 0 | 0 | 0 | 16 | 0 | 0 | 0 |
| ASV_47 | 0 | 0 | 0 | 0 | 0 | 0 | 0 | 0 | 0 |
| ASV_48 | 0 | 0 | 0 | 0 | 0 | 0 | 0 | 0 | 0 |
| ASV_49 | 0 | 0 | 0 | 0 | 0 | 22 | 58 | 0 | 0 |
| ASV_50 | 0 | 0 | 0 | 0 | 0 | 0 | 0 | 0 | 0 |
| ASV_57 | 0 | 0 | 0 | 56 | 54 | 62 | 0 | 0 | 0 |
| ASV_58 | 0 | 0 | 0 | 19 | 0 | 0 | 0 | 0 | 0 |
| ASV_59 | 0 | 0 | 0 | 0 | 0 | 0 | 22 | 0 | 15 |
| ASV_61 | 0 | 0 | 0 | 0 | 0 | 0 | 0 | 0 | 0 |
| ASV_62 | 0 | 0 | 0 | 0 | 0 | 0 | 0 | 0 | 0 |
| ASV_75 | 0 | 0 | 0 | 0 | 0 | 0 | 0 | 0 | 0 |
| ASV_76 | 0 | 0 | 0 | 0 | 0 | 0 | 0 | 0 | 0 |
| ASV_81 | 0 | 0 | 0 | 0 | 0 | 0 | 0 | 0 | 0 |
| ASV_89 | 0 | 0 | 0 | 0 | 0 | 0 | 0 | 0 | 0 |
| ASV_91 | 0 | 0 | 0 | 0 | 0 | 0 | 0 | 0 | 0 |
| ASV_93 | 0 | 0 | 0 | 0 | 0 | 0 | 0 | 0 | 0 |
| ASV_94 | 0 | 0 | 0 | 0 | 0 | 0 | 0 | 0 | 0 |
| ASV_97 | 0 | 0 | 0 | 0 | 0 | 0 | 0 | 0 | 0 |
| ASV_104 | 0 | 0 | 0 | 0 | 0 | 0 | 0 | 0 | 0 |
| ASV_105 | 0 | 0 | 0 | 0 | 0 | 6 | 0 | 0 | 0 |
| ASV_108 | 0 | 0 | 0 | 28 | 0 | 60 | 0 | 0 | 0 |
| ASV_134 | 0 | 0 | 0 | 28 | 29 | 0 | 0 | 0 | 0 |
| ASV_137 | 0 | 0 | 0 | 0 | 0 | 0 | 0 | 0 | 0 |
| ASV_140 | 0 | 0 | 0 | 0 | 46 | 0 | 0 | 0 | 0 |
| ASV_144 | 0 | 0 | 0 | 0 | 0 | 0 | 0 | 0 | 0 |
| ASV_146 | 0 | 0 | 0 | 0 | 0 | 0 | 0 | 0 | 0 |
| ASV_175 | 0 | 0 | 0 | 0 | 0 | 0 | 0 | 0 | 0 |
| ASV_204 | 0 | 0 | 0 | 0 | 0 | 0 | 0 | 0 | 0 |
| | 27040 | 19947 | 32862 | 2200 | 2681 | 3343 | 33790 | 36007 | 33266 |

**Table S3.** Statistical analyses of the metabarcoding data evaluating the effects of substrate, nixtamalisation, time, and location. The Kruskal-Wallis and Tukey methods were applied for measures of alpha diversity. PERMANOVA and Deseq were used for beta diversity analyses.

**Kruskal-Wallis, alpha-diversity**

| | **Substrates: 8 types** | | | | **Treatment: standard vs nixt** | | | | **Time: start vs end** | | | | **Location: NOMA vs INUA** | | | |
|---|---|---|---|---|---|---|---|---|---|---|---|---|---|---|---|---|
| | ***16S*** | | ***ITS*** | | ***16S*** | | ***ITS*** | | ***16S*** | | ***ITS*** | | ***16S*** | | ***ITS*** | |
| | chi-squarec | p-value | chi-squarec | p-value | chi-squared | p-value | chi-squared | p-value | chi-squared | p-value | chi-squared | p-value | chi-squarec | p-value | chi-squarec | p-value |
| Observed | 17.91000 | 0.01238 | 7.02260 | 0.42650 | 0.20095 | 0.65400 | 0.72929 | 0.39310 | 1.30300 | 0.25370 | 17.81400 | 0.00002 | 0.88375 | 0.34720 | 7.49220 | 0.00620 |
| Chao1 | 17.33100 | 0.01538 | 7.38620 | 0.38980 | 0.07188 | 0.78860 | 0.89763 | 0.34340 | 1.10430 | 0.29330 | 17.79400 | 0.00002 | 0.97433 | 0.32360 | 8.76460 | 0.00307 |
| Shannon | 19.18700 | 0.00762 | 7.09330 | 0.41920 | 1.42110 | 0.23320 | 0.04873 | 0.82530 | 12.73500 | 0.00036 | 16.41100 | 0.00005 | 0.09677 | 0.75570 | 6.45430 | 0.01107 |
| InvSimp | 18.73300 | 0.00907 | 6.72000 | 0.45860 | 2.12280 | 0.14510 | 0.23587 | 0.62720 | 10.93900 | 0.00094 | 15.36900 | 0.00009 | 0.04301 | 0.83570 | 5.68820 | 0.01708 |

**Tukey**

**Bacteria**

| | **Observed** | **groups** | **Chao1** | **groups** |
|---|---|---|---|---|
| Lentil | 19 | a | 20 | a |
| Rye-bread | 19 | a | 19.44444 | a |
| Yellow-pea | 17.33333 | ab | 17.33333 | ab |
| Lentil-Nixt | 16.33333 | abc | 16.5 | ab |
| Yellow-pea | 14.66667 | abc | 15 | ab |
| Fava-Nixt | 13.66667 | abc | 13.66667 | ab |
| Fava | 11 | bc | 11 | bc |
| Soy | 10.66667 | c | 10.83333 | bc |

| | **Shannon** | **groups** | **InvSimpsor** | **groups** |
|---|---|---|---|---|
| Rye-bread | 1.36229 | a | 2.886239 | a |
| Yellow-pea | 1.265495 | a | 2.728478 | ab |
| Lentil | 1.262125 | a | 2.41788 | abc |
| Soy | 0.678453 | b | 1.553587 | bc |
| Fava-Nixt | 0.748955 | b | 1.405315 | c |
| Yellow-pea | 0.714926 | b | 1.374115 | c |
| Lentil-Nixt | 0.64184 | b | 1.314413 | c |
| Fava | 0.524263 | b | 1.275738 | c |

**PERMANOVA**

| | **Factors** | **Df** | **SumsOfSqs** | **R2** | **F.Model** | **Pr(>F)** |
|---|---|---|---|---|---|---|
| **Bacteria** | Substrate | 7 | 4.7589 | 0.90311 | 21.305 | 0.001 |
| | Residuals | 16 | 0.5106 | 0.09689 | | |
| | Total | 23 | 5.2694 | 1 | | |
| | Time | 1 | 2.2865 | 0.27404 | 11.325 | 0.001 |
| | Residuals | 30 | 6.0571 | 0.72596 | | |
| | Total | 31 | 8.3437 | 1 | | |
| | Treatment | 1 | 0.73148 | 0.35454 | 8.7885 | 0.008 |
| | Residuals | 16 | 1.3317 | 0.64546 | | |
| | Total | 17 | 2.06318 | 1 | | |
| | Location | 1 | 0.9865 | 0.12311 | 3.9309 | 0.019 |
| | Residuals | 28 | 7.027 | 0.87689 | | |
| | Total | 29 | 8.0135 | 1 | | |
| **Fungi** | Substrate | 7 | 0.83418 | 0.51388 | 2.4163 | 0.16 |
| | Residuals | 16 | 0.78911 | 0.48612 | | |
| | Total | 23 | 1.62329 | 1 | | |
| | Time | 1 | 0.08484 | 0.04983 | 1.5732 | 0.131 |
| | Residuals | 30 | 1.61788 | 0.95017 | | |
| | Total | 31 | 1.70272 | 1 | | |
| | Treatment | 1 | 0.17278 | 0.13579 | 2.514 | 0.078 |
| | Residuals | 16 | 1.09967 | 0.86421 | | |
| | Total | 17 | 1.27245 | 1 | | |
| | Location | 1 | 0.6317 | 0.17443 | 5.9161 | 0.008 |
| | Residuals | 28 | 2.9898 | 0.82557 | | |
| | Total | 29 | 3.6215 | 1 | | |

| Product | Treatment | Kingdom | log2FoldChange | baseMean | ASV | Species | $p_{adj}$ |
|---|---|---|---|---|---|---|---|
| Substrate | Soy vs yellow pea | Bacteria | 8.50994 | 43,201.5 | Cluster_1 | *Pediococcus pentosaceus* | 1.11068e-33 |
| | | | 8.05739 | 3357.79 | Cluster_3 | *Enterobacteriaceae family* | 3.50853e-173 |
| | | | 3.48172 | 2096.64 | Cluster_4 | *Bacillus sp.* | 2.42972e-30 |
| | Soy vs Gotland lentil | Bacteria | 6.82378 | 3357.79 | Cluster_3 | *Enterobacteriaceae family* | 3.2125E-124 |
| | | | 9.61815 | 43201.5 | Cluster_1 | *Pediococcus pentosaceus* | 1.22344E-42 |
| | | | 3.6329 | 2096.64 | Cluster_4 | *Bacillus sp.* | 6.87943E-33 |
| | | | -12.1196 | 1016.24 | Cluster_13 | *Tetragenococcus halophilus* | 8.04746E-09 |
| | | Fungi | -27.7616 | 57.3131 | Cluster_19 | *Millerozyma farinosa* | 5.47247e-7 |
| | Soy vs fava | Bacteria | 6.472 | 3357.79 | Cluster_3 | *Enterobacteriaceae family* | 3.9815E-111 |
| | | | 13.0475 | 43201.5 | Cluster_1 | *Pediococcus pentosaceus* | 1.7067E-77 |
| | | | -25.4138 | 1016.24 | Cluster_13 | *Tetragenococcus halophilus* | 1.38238E-33 |
| | | | 3.38297 | 2096.64 | Cluster_4 | *Bacillus sp.* | 2.25028E-28 |
| | Soy vs yellow pea nixtamilised | Bacteria | -54.5387 | 1016.24 | Cluster_13 | *Tetragenococcus halophilus* | 5.7416E-151 |
| | | | -7.28534 | 8520.75 | Cluster_2 | *Staphylococcus sp.* | 7.39838E-49 |
| | | | 11.0822 | 43201.5 | Cluster_1 | *Pediococcus pentosaceus* | 3.5155E-56 |
| | | | 6.41566 | 3357.79 | Cluster_3 | *Enterobacteriaceae family* | 5.0949E-110 |
| | | | 3.50048 | 2096.64 | Cluster_4 | *Bacillus sp.* | 9.91041E-31 |
| | | Fungi | -15.2649 | 5340.93 | Cluster_2 | *Millerozyma farinosa* | 1.38450e-18 |
| | Soy vs Gotland lentil nixt | Bacteria | -10.6971 | 1016.24 | Cluster_13 | *Tetragenococcus halophilus* | 2.297E-07 |
| | | | -6.75406 | 8520.75 | Cluster_2 | *Staphylococcus sp.* | 2.35508E-42 |
| | | | 11.7622 | 43201.5 | Cluster_1 | *Pediococcus pentosaceus* | 2.61765E-63 |
| | | | 6.57973 | 3357.79 | Cluster_3 | *Enterobacteriaceae family* | 1.4624E-115 |
| | | | 4.08292 | 2096.64 | Cluster_4 | *Bacillus sp.* | 1.79938E-41 |
| | | Fungi | -17.6899 | 5340.93 | Cluster_2 | *Millerozyma farinosa* | 8.19830e-25 |
| | | | -3.12239 | 58,071.2 | Cluster_1 | *Aspergillus oryzae* | 0.000288214 |
| | | | 32.0069 | 194.904 | Cluster_4 | *Zygosaccharomyces parabailii-bailii* | 5.89754e-14 |
| | | | 30.8736 | 56.0793 | Cluster_12 | *Schizosaccharomyces sp.* | 4.01916e-7 |
| | | | 9.66646 | 193.356 | Cluster_9 | *[Candida] glabrata* | 3.31776E-06 |
| | Soy vs fava nixt | Bacteria | -8.08345 | 1016.24 | Cluster_13 | *Tetragenococcus halophilus* | 0.000164988 |
| | | | -6.45183 | 8520.75 | Cluster_2 | *Staphylococcus sp.* | 2.87595E-38 |
| | | | 11.415 | 43201.5 | Cluster_1 | *Pediococcus pentosaceus* | 1.44739E-59 |
| | | | 6.54964 | 3357.79 | Cluster_3 | *Enterobacteriaceae family* | 4.7067E-114 |
| | | | 4.18754 | 2096.64 | Cluster_4 | *Bacillus sp.* | 2.66906E-43 |
| | | Fungi | -15.3442 | 5340.93 | Cluster_2 | *Millerozyma farinosa* | 6.09274E-19 |
| | | | 11.5316 | 193.356 | Cluster_9 | *[Candida] glabrata* | 3.89524E-08 |
| | Soy vs rye bread | Bacteria | -12.6968 | 1016.24 | Cluster_13 | *Tetragenococcus halophilus* | 8.15411E-10 |
| | | | -10.6199 | 8520.75 | Cluster_2 | *Staphylococcus sp.* | 4.821E-101 |
| | | | 7.95702 | 3357.79 | Cluster_3 | *Enterobacteriaceae family* | 7.9873E-169 |
| | | | 5.34653 | 2096.64 | Cluster_4 | *Bacillus sp.* | 5.49892E-70 |
| | | Fungi | -15.7613 | 5340.93 | Cluster_2 | *Millerozyma farinosa* | 4.52174E-20 |
| Time | Start vs End | Bacteria | 12.7611 | 36,696.4 | Cluster_1 | *Pediococcus pentosaceus* | 8.62768e-66 |
| | | | 28.3819 | 524.835 | Cluster_13 | *Tetragenococcus halophilus* | 1.27020e-49 |
| | | | 7.4128 | 6923.93 | Cluster_2 | *Staphylococcus* | 9.44810e-28 |
| | | | -1.61572 | 1490.64 | Cluster_6 | *Enterobacteriaceae family* | 8.48343E-05 |
| | | | -2.03185 | 805.354 | Cluster_10 | *Bacillus* | 0.000661883 |
| | | Fungi | 20.2496 | 7.59856 | Cluster_4 | *Zygosaccharomyces parabailii-bailii* | 1.54524e-9 |
| | | | -2.59502 | 77,498.2 | Cluster_1 | *Aspergillus oryzae* | 1.54524e-9 |
| | | | 5.07547 | 29.2362 | Cluster_18 | *Saccharomycopsis fibuligera* | 1.69443e-8 |
| | | | 8.89726 | 1660.22 | Cluster_2 | *Millerozyma farinosa* | 1.33658e-7 |
| | | | 5.58216 | 43.1051 | Cluster_6 | *[Candida] glabrata* | 6.94074E-05 |
| Treatment | Boiled vs Nixtamalised | Bacteria | -6.89315 | 5545.57 | Cluster_2 | *Staphylococcus sp.* | 6.85910e-117 |
| | | | -0.910104 | 2155.65 | Cluster_3 | *Enterobacteriaceae family* | 9.97063e-7 |
| | | | 1.96686 | 296.697 | Cluster_10 | *Bacillus sp.* | 5.60794E-06 |
| | | Fungi | -17.1183176 | 9739.468342 | Cluster_2 | *Millerozyma farinosa* | 1.87563E-71 |
| | | | 28.97424875 | 707.818516 | Cluster_4 | *Zygosaccharomyces parabailii-bailii* | 1.25307E-28 |
| | | | 5.175894499 | 254.164077 | Cluster_6 | *[Candida] glabrata* | 4.25736E-07 |
| Location | NOMA vs INUA | Bacteria | 10.8104 | 869.367 | Cluster_14 | *Enterococcus sp.* | 2.34927e-11 |
| | | | 11.0338 | 4421.53 | Cluster_5 | *Pediococcus acidilactici* | 4.83348e-9 |
| | | | 9.93118 | 64.0426 | Cluster_19 | *Weissella sp.* | 1.33806e-8 |
| | | | -4.55524 | 36,075.3 | Cluster_1 | *Pediococcus pentosaceus* | 4.38019e-8 |
| | | | 6.94622 | 6.0684 | Cluster_11 | *Lacticaseibacillus sp.* | 5.20970e-8 |
| | | | -9.99884 | 1119.82 | Cluster_13 | *Tetragenococcus halophilus* | 1.0398E-06 |
| | | Fungi | -30 | 3285.24 | Cluster_2 | *Millerozyma farinosa* | 5.50599e-84 |
| | | | 9.36037 | 46.1304 | Cluster_11 | *Candida sake* | 3.05951e-10 |
| | | | 9.82037 | 63.8507 | Cluster_8 | *Debaryomyces hansenii* | 2.24147e-9 |

Log2FoldChange negative values indicates that the ASV is more abundant in the first factor when compared to the secondone. Table is showing only most abundant ASVs and significant values for Log2FoldChange ($p < 0.001$).

**Table S4.** 'metaphan', the relative abundance of microbial communities in the 38 miso metagenomes using MetaPhlAn taxonomic assignment

| Kingdom | Phylum | Class | Order | Family | Genus | Species | F.1 | F1.3 | F2.3 | F3.3 | NF.1 | NF1.3 | NF2.3 | NF3.3 | L.1 | L1.3 | L2.3 | L3.3 | NL.1 | NL1.3 | NL2.3 | NL3.3 | P.1 | P1.3 | P2.3 | P3.3 | NP.1 | NP1.3 | NP2.3 | NP3.3 | R.1 | R1.3 | R2.3 | R3.3 | S.1 | S1.3 | S2.3 | S3.3 | HB | MS | PB | PR | SY | TS |
|---|---|---|---|---|---|---|---|---|---|---|---|---|---|---|---|---|---|---|---|---|---|---|---|---|---|---|---|---|---|---|---|---|---|---|---|---|---|---|---|---|---|---|---|---|
| k__Bacteria | p__Firmicutes | c__Bacilli | o__Lactobacillales | f__Lactobacillaceae | g__Pediococcus | s__Pediococcus_pentosaceus | 0 | 13.53 | 39.17 | 36.43 | 0.14 | 86.98 | 89.21 | 95.69 | 0 | 3.49 | 14 | 33.4 | 0 | 95.9 | 95.4 | 94.1 | 0 | 3.61 | 28.5 | 33.5 | 0 | 78 | 56.3 | 84.8 | 0 | 0.19 | 0 | 0 | 0 | 0 | 0 | 0 | 76.4 | 0.13 | 0.02 | 0.08 | 0 | 3.09 |
| k__Bacteria | p__Firmicutes | c__Bacilli | o__Bacillales | f__Staphylococcaceae | g__Staphylococcus | s__Staphylococcus_epidermidis | 0 | 6.945 | 3.77 | 2.999 | 0 | 5.296 | 4.047 | 0.752 | 0 | 8.48 | 13.9 | 14.9 | 0 | 0.11 | 0.18 | 0.27 | 0 | 8.98 | 9.63 | 7.45 | 0 | 5.66 | 14.2 | 2.83 | 0 | 0.04 | 0.13 | 0.08 | 6.82 | 89.2 | 87.5 | 84.6 | 0.42 | 15.4 | 1.76 | 51.8 | 4.14 | 8.03 |
| k__Bacteria | p__Firmicutes | c__Bacilli | o__Lactobacillales | f__Lactobacillaceae | g__Pediococcus | s__Pediococcus_acidilactici | 0 | 0 | 0 | 0 | 0 | 0 | 0.003 | 0 | 0 | 0 | 0 | 0 | 0 | 0 | 0 | 0 | 0 | 0 | 0 | 0 | 0 | 0 | 0 | 0 | 0 | 0 | 0 | 0 | 0 | 0 | 0 | 0 | 0 | 1.19 | 87.3 | 7.74 | 0.28 | 45.9 |
| k__Bacteria | p__Firmicutes | c__Bacilli | o__Bacillales | f__Staphylococcaceae | g__Staphylococcus | s__Staphylococcus_pasteuri | 0 | 75.97 | 53.63 | 58.38 | 0 | 3.497 | 2.823 | 0.41 | 0 | 86.4 | 69.2 | 40.7 | 0 | 0.23 | 0.77 | 0.94 | 0.23 | 85.3 | 58.4 | 57 | 0 | 12.3 | 23.8 | 8.66 | 0 | 0.06 | 0 | 0.03 | 1.48 | 7.57 | 11 | 10.5 | 12.1 | 3.95 | 0.28 | 0.13 | 7.89 | 5.64 |
| k__Eukaryota | p__Ascomycota | c__Eurotiomyce | o__Eurotiales | f__Aspergillaceae | g__Aspergillus | s__Aspergillus_oryzae | 43.42 | 0.524 | 0.404 | 0.247 | 17.07 | 0.174 | 0.087 | 0.092 | 59.6 | 0.66 | 0.43 | 0.34 | 27.5 | 0.09 | 0.12 | 0.17 | 21.6 | 0.21 | 0.14 | 0.12 | 44.8 | 0.61 | 0.41 | 0.29 | 41.6 | 20 | 20.1 | 17.1 | 85.7 | 0.36 | 0.3 | 0.27 | 0.17 | 0.04 | 0.12 | 31 | 0.22 | 0 |
| k__Bacteria | p__Proteobacteria | c__Gammaprote | o__Enterobacterales | f__Enterobacteriaceae | g__Cronobacter | s__Cronobacter_malonaticus | 42.01 | 0.356 | 0.248 | 0.281 | 38.66 | 0.312 | 0.333 | 0.266 | 34.9 | 0.43 | 0.31 | 0.82 | 21.5 | 0.27 | 0.32 | 0.35 | 71.9 | 0.61 | 0.88 | 0.63 | 14.8 | 0.23 | 0.2 | 0.15 | 18.8 | 26 | 29.4 | 26.3 | 0.55 | 0 | 0 | 0.01 | 0 | 0 | 0 | 0 | 0 | 0 |
| k__Bacteria | p__Firmicutes | c__Bacilli | o__Lactobacillales | f__Enterococcaceae | g__Enterococcus | s__Enterococcus_faecium | 0 | 0 | 0 | 0 | 0 | 0 | 0 | 0 | 0 | 0 | 0 | 0 | 0 | 0 | 0 | 0 | 0 | 0 | 0 | 0 | 0 | 0 | 0 | 0 | 0 | 0 | 0 | 0 | 0 | 0 | 0 | 0 | 0.03 | 0.16 | 0 | 0 | 67 | 0.49 |
| k__Bacteria | p__Firmicutes | c__Bacilli | o__Bacillales | f__Staphylococcaceae | g__Staphylococcus | s__Staphylococcus_warneri | 0.427 | 2.103 | 2.069 | 1.045 | 0 | 0 | 0.212 | 0.011 | 0 | 0.3 | 1.23 | 4.8 | 0 | 0 | 0 | 0 | 1.41 | 0.6 | 1.28 | 0.5 | 0 | 0.01 | 0 | 0 | 0 | 0.42 | 0 | 0.05 | 0 | 0.22 | 0.34 | 0.23 | 0.43 | 54 | 10.1 | 5.87 | 12.8 | 21.2 |
| k__Bacteria | p__Proteobacteria | c__Gammaprote | o__Enterobacterales | f__Enterobacteriaceae | g__Enterobacter | s__Enterobacter_cloacae_complex | 14.01 | 0.084 | 0.049 | 0.051 | 30.02 | 0.315 | 0.312 | 0.34 | 4.76 | 0.12 | 0.07 | 0.23 | 42.7 | 0.64 | 0.52 | 0.72 | 4.6 | 0.11 | 0.11 | 0.08 | 35.6 | 0.85 | 0.68 | 0.51 | 31 | 47.1 | 44.7 | 50.7 | 1.39 | 0 | 0 | 0 | 0 | 0 | 0.12 | 0 | 0 | 0 |
| k__Bacteria | p__Firmicutes | c__Bacilli | o__Lactobacillales | f__Lactobacillaceae | g__Lactobacillus | s__Lactobacillus_plantarum | 0 | 0.075 | 0.572 | 0.499 | 0 | 2.261 | 2.614 | 2.246 | 0 | 0 | 0.07 | 1.68 | 0 | 2.58 | 2.45 | 3.15 | 0 | 0.03 | 0.94 | 0.64 | 0 | 1.81 | 1.2 | 2.15 | 0 | 0 | 0 | 0 | 0 | 0 | 0 | 0 | 0.82 | 25 | 0.06 | 0 | 0.04 | 4 |
| k__Bacteria | p__Firmicutes | c__Bacilli | o__Bacillales | f__Bacillaceae | g__Bacillus | s__Bacillus_cereus_group | 0 | 0.001 | 0 | 0 | 12.69 | 0.138 | 0.062 | 0.077 | 0 | 0 | 0 | 0.02 | 7.13 | 0.06 | 0.04 | 0.1 | 0.16 | 0 | 0 | 0 | 1.55 | 0.05 | 0.04 | 0.03 | 0.58 | 0.51 | 0.42 | 0.78 | 0 | 0 | 0 | 0 | 0.01 | 0 | 0 | 0 | 0 | 0 |
| k__Bacteria | p__Firmicutes | c__Bacilli | o__Lactobacillales | f__Enterococcaceae | g__Enterococcus | s__Enterococcus_faecalis | 0 | 0 | 0 | 0 | 0 | 0 | 0 | 0 | 0 | 0 | 0.01 | 0 | 0 | 0 | 0 | 0 | 0 | 0 | 0 | 0 | 0 | 0 | 0 | 0 | 0 | 0 | 0 | 0 | 0 | 0 | 0 | 0 | 0.09 | 0.06 | 0 | 0 | 0 | 11.2 |
| k__Bacteria | p__Firmicutes | c__Bacilli | o__Lactobacillales | f__Lactobacillaceae | g__Lactobacillus | s__Lactobacillus_sanfranciscensis | 0 | 0 | 0 | 0 | 0 | 0 | 0 | 0 | 0 | 0 | 0 | 0 | 0 | 0 | 0 | 0 | 0 | 0 | 0 | 0 | 0 | 0 | 0 | 0 | 6.58 | 3.13 | 3.08 | 2.34 | 0 | 0 | 0 | 0 | 0 | 0 | 0 | 0 | 0 | 0 |
| k__Bacteria | p__Firmicutes | c__Bacilli | o__Lactobacillales | f__Lactobacillaceae | g__Lactobacillus | s__Lactobacillus_sakei | 0 | 0 | 0 | 0 | 0 | 0.121 | 0.096 | 0.064 | 0 | 0 | 0 | 0 | 0 | 0.05 | 0.04 | 0.03 | 0 | 0 | 0 | 0 | 0 | 0 | 0 | 0 | 0 | 0 | 0 | 0 | 0 | 0 | 0 | 0 | 4.61 | 0 | 0 | 0 | 0 | 0.12 |
| k__Bacteria | p__Firmicutes | c__Bacilli | o__Lactobacillales | f__Leuconostocaceae | g__Weissella | s__Weissella_confusa | 0 | 0 | 0 | 0 | 0 | 0 | 0 | 0 | 0 | 0 | 0 | 0 | 0 | 0 | 0 | 0 | 0 | 0 | 0 | 0 | 0 | 0 | 0 | 0 | 0 | 0 | 0 | 0 | 0 | 0 | 0 | 0 | 0.36 | 0 | 0.01 | 0 | 4.24 | 0.01 |
| k__Bacteria | p__Firmicutes | c__Bacilli | o__Lactobacillales | f__Enterococcaceae | g__Tetragenococcus | s__Tetragenococcus_halophilus | 0 | 0 | 0 | 0 | 0 | 0.458 | 0.004 | 0 | 0 | 0 | 0 | 0 | 0 | 0 | 0 | 0 | 0 | 0.02 | 0 | 0 | 0 | 0 | 0 | 0 | 0 | 0 | 0 | 0 | 0 | 2.22 | 0.5 | 4.14 | 0.02 | 0 | 0 | 0 | 0 | 0 |
| k__Bacteria | p__Firmicutes | c__Bacilli | o__Bacillales | f__Staphylococcaceae | g__Staphylococcus | s__Staphylococcus_aureus | 0 | 0.032 | 0.024 | 0.007 | 0 | 0 | 0.035 | 0 | 0 | 0.04 | 0.44 | 2.78 | 0 | 0 | 0 | 0 | 0.02 | 0.02 | 0.07 | 0.02 | 0 | 0 | 0 | 0 | 0 | 0 | 0 | 0 | 0.43 | 0.1 | 0.16 | 0.12 | 2.62 | 0 | 0 | 0 | 0 | 0.06 |
| k__Bacteria | p__Firmicutes | c__Bacilli | o__Bacillales | f__Staphylococcaceae | g__Staphylococcus | s__Staphylococcus_simulans | 0 | 0 | 0 | 0 | 0 | 0 | 0 | 0 | 0 | 0 | 0 | 0 | 0 | 0 | 0 | 0 | 0 | 0 | 0 | 0 | 0 | 0.39 | 2.73 | 0.5 | 0 | 0 | 0 | 0 | 0 | 0 | 0 | 0 | 0 | 0 | 0 | 0 | 0 | 0 |
| k__Bacteria | p__Proteobacteria | c__Gammaprote | o__Enterobacterales | f__Enterobacteriaceae | g__Kosakonia | s__Kosakonia_cowanii | 0.126 | 0.006 | 0.002 | 0 | 1.428 | 0.018 | 0.006 | 0.008 | 0.34 | 0 | 0 | 0 | 0.91 | 0.03 | 0.02 | 0.02 | 0.05 | 0 | 0 | 0 | 2.61 | 0.03 | 0.04 | 0.02 | 0.55 | 1.27 | 0.87 | 1.47 | 0 | 0 | 0 | 0 | 0 | 0 | 0 | 0 | 0 | 0 |
| k__Bacteria | p__Firmicutes | c__Bacilli | o__Bacillales | f__Staphylococcaceae | g__Staphylococcus | s__Staphylococcus_lugdunensis | 0 | 0.016 | 0 | 0 | 0 | 0 | 0 | 0 | 0 | 0.01 | 0.04 | 0.04 | 0 | 0 | 0 | 0 | 0 | 0.36 | 0.06 | 0.05 | 0 | 0 | 0.01 | 0 | 0 | 0 | 0 | 0 | 0 | 0 | 0 | 0 | 0 | 0 | 0 | 1.93 | 0 | 0 |
| k__Bacteria | p__Firmicutes | c__Bacilli | o__Bacillales | f__Staphylococcaceae | g__Staphylococcus | s__Staphylococcus_xylosus | 0 | 0 | 0 | 0 | 0 | 0.035 | 0.049 | 0 | 0 | 0 | 0 | 0 | 0 | 0 | 0 | 0 | 0 | 0 | 0 | 0 | 0 | 0 | 0 | 0 | 0 | 0 | 0 | 0 | 1.75 | 0.13 | 0.02 | 0.01 | 0 | 0 | 0 | 0.13 | 0 | 0 |
| k__Eukaryota | p__Ascomycota | c__Saccharomyc | o__Saccharomycetale | f__Saccharomycetaceae | g__Saccharomyces | s__Saccharomyces_cerevisiae | 0 | 0 | 0 | 0 | 0 | 0.002 | 0.018 | 0 | 0 | 0 | 0 | 0 | 0 | 0 | 0 | 0.02 | 0 | 0 | 0 | 0 | 0 | 0.01 | 0 | 0 | 0 | 0 | 0 | 0 | 0 | 0 | 0 | 0 | 0.03 | 0.03 | 0.05 | 0 | 1.7 | 0 |
| k__Bacteria | p__Firmicutes | c__Bacilli | o__Lactobacillales | f__Lactobacillaceae | g__Lactobacillus | s__Lactobacillus_mindensis | 0 | 0 | 0 | 0 | 0 | 0 | 0 | 0 | 0 | 0 | 0 | 0 | 0 | 0 | 0 | 0 | 0 | 0 | 0 | 0 | 0 | 0 | 0 | 0 | 0.89 | 1.15 | 1.05 | 0.87 | 0 | 0 | 0 | 0 | 0 | 0 | 0 | 0 | 0 | 0 |
| k__Bacteria | p__Firmicutes | c__Bacilli | o__Bacillales | f__Staphylococcaceae | g__Staphylococcus | s__Staphylococcus_saprophyticus | 0 | 0.178 | 0 | 0 | 0 | 0 | 0.02 | 0.012 | 0 | 0 | 0 | 0 | 0 | 0 | 0.07 | 0.03 | 0 | 0 | 0 | 0 | 0 | 0.03 | 0.37 | 0.06 | 0 | 0 | 0 | 0 | 0.41 | 0.01 | 0 | 0 | 0.02 | 0 | 0 | 0 | 0.79 | 0 |
| k__Bacteria | p__Firmicutes | c__Bacilli | o__Lactobacillales | f__Lactobacillaceae | g__Lactobacillus | s__Lactobacillus_curvatus | 0 | 0 | 0 | 0 | 0 | 0 | 0.009 | 0.004 | 0 | 0 | 0 | 0 | 0 | 0.02 | 0.03 | 0.04 | 0 | 0 | 0 | 0 | 0 | 0 | 0 | 0 | 0 | 0 | 0 | 0 | 0 | 0 | 0 | 0 | 0.71 | 0 | 0 | 0 | 0 | 0.01 |
| k__Bacteria | p__Firmicutes | c__Bacilli | o__Lactobacillales | f__Leuconostocaceae | g__Weissella | s__Weissella_viridescens | 0 | 0 | 0 | 0 | 0 | 0.027 | 0.031 | 0.027 | 0 | 0 | 0 | 0 | 0 | 0.02 | 0.02 | 0.02 | 0 | 0 | 0 | 0 | 0 | 0 | 0 | 0 | 0 | 0 | 0 | 0 | 0 | 0 | 0 | 0 | 0.7 | 0 | 0 | 0 | 0 | 0 |
| k__Bacteria | p__Firmicutes | c__Bacilli | o__Bacillales | f__Staphylococcaceae | g__Staphylococcus | s__Staphylococcus_capitis | 0 | 0.023 | 0.015 | 0.042 | 0 | 0.004 | 0.027 | 0 | 0.32 | 0.02 | 0.28 | 0.17 | 0 | 0 | 0 | 0 | 0 | 0.13 | 0.05 | 0.02 | 0 | 0.01 | 0.02 | 0.05 | 0 | 0 | 0 | 0 | 0.67 | 0.14 | 0.18 | 0.11 | 0 | 0 | 0 | 0 | 0 | 0 |
| k__Bacteria | p__Proteobacteria | c__Gammaprote | o__Enterobacterales | f__Erwiniaceae | g__Pantoea | s__Pantoea_agglomerans | 0 | 0 | 0 | 0 | 0 | 0 | 0 | 0 | 0 | 0 | 0 | 0 | 0 | 0 | 0 | 0.01 | 0 | 0 | 0 | 0 | 0.59 | 0 | 0 | 0 | 0 | 0 | 0.13 | 0.2 | 0 | 0 | 0 | 0 | 0 | 0 | 0 | 0 | 0 | 0 |
| k__Bacteria | p__Firmicutes | c__Bacilli | o__Bacillales | f__Staphylococcaceae | g__Staphylococcus | s__Staphylococcus_succinus | 0 | 0 | 0 | 0 | 0 | 0 | 0 | 0 | 0 | 0 | 0 | 0 | 0 | 0 | 0 | 0 | 0 | 0 | 0 | 0 | 0 | 0 | 0 | 0 | 0 | 0 | 0 | 0 | 0.38 | 0.02 | 0 | 0 | 0 | 0 | 0 | 0 | 0 | 0 |
| k__Bacteria | p__Firmicutes | c__Bacilli | o__Bacillales | f__Bacillaceae | g__Bacillus | s__Bacillus_intestinalis | 0 | 0 | 0 | 0 | 0 | 0 | 0 | 0 | 0 | 0 | 0 | 0 | 0 | 0 | 0 | 0 | 0 | 0 | 0 | 0 | 0 | 0 | 0 | 0 | 0 | 0 | 0 | 0 | 0 | 0 | 0 | 0 | 0 | 0 | 0.01 | 0.34 | 0.02 | 0 |
| k__Bacteria | p__Firmicutes | c__Bacilli | o__Bacillales | f__Staphylococcaceae | g__Staphylococcus | s__Staphylococcus_sciuri | 0 | 0 | 0 | 0 | 0 | 0 | 0 | 0 | 0 | 0 | 0 | 0 | 0 | 0 | 0 | 0 | 0 | 0 | 0 | 0 | 0 | 0 | 0 | 0 | 0 | 0 | 0 | 0 | 0 | 0 | 0 | 0 | 0 | 0 | 0 | 0.29 | 0.33 | 0 |
| k__Bacteria | p__Firmicutes | c__Bacilli | o__Bacillales | f__Bacillaceae | g__Bacillus | s__Bacillus_murimartini | 0 | 0 | 0 | 0 | 0 | 0 | 0 | 0 | 0 | 0 | 0 | 0 | 0 | 0 | 0 | 0 | 0 | 0 | 0 | 0 | 0 | 0 | 0 | 0 | 0 | 0 | 0 | 0 | 0 | 0 | 0 | 0 | 0.02 | 0.02 | 0.12 | 0.29 | 0.3 | 0.06 |
| k__Bacteria | p__Proteobacteria | c__Gammaprote | o__Pseudomonadale | f__Moraxellaceae | g__Acinetobacter | s__Acinetobacter_pittii | 0 | 0 | 0 | 0 | 0 | 0 | 0 | 0 | 0 | 0 | 0 | 0 | 0.28 | 0 | 0 | 0 | 0 | 0 | 0 | 0 | 0 | 0 | 0 | 0 | 0.01 | 0.04 | 0.06 | 0.06 | 0 | 0 | 0 | 0 | 0 | 0 | 0 | 0 | 0 | 0 |
| k__Bacteria | p__Firmicutes | c__Bacilli | o__Bacillales | f__Staphylococcaceae | g__Staphylococcus | s__Staphylococcus_haemolyticus | 0 | 0.009 | 0.03 | 0.023 | 0 | 0 | 0 | 0 | 0 | 0 | 0.05 | 0.08 | 0 | 0 | 0 | 0 | 0 | 0 | 0.01 | 0 | 0 | 0 | 0 | 0 | 0 | 0 | 0 | 0 | 0 | 0 | 0 | 0 | 0 | 0 | 0.01 | 0 | 0.23 | 0 |
| k__Bacteria | p__Firmicutes | c__Bacilli | o__Bacillales | f__Bacillaceae | g__Bacillus | s__Bacillus_circulans | 0 | 0 | 0 | 0 | 0 | 0 | 0 | 0 | 0 | 0 | 0 | 0 | 0 | 0 | 0 | 0 | 0 | 0 | 0 | 0 | 0 | 0 | 0 | 0 | 0 | 0 | 0 | 0 | 0 | 0 | 0 | 0 | 0 | 0 | 0 | 0.23 | 0 | 0 |
| k__Bacteria | p__Firmicutes | c__Bacilli | o__Bacillales | f__Staphylococcaceae | g__Staphylococcus | s__Staphylococcus_hominis | 0 | 0.148 | 0.011 | 0 | 0 | 0 | 0 | 0 | 0 | 0 | 0.01 | 0.03 | 0 | 0 | 0 | 0 | 0 | 0 | 0 | 0 | 0 | 0 | 0 | 0 | 0 | 0 | 0 | 0 | 0.23 | 0 | 0 | 0 | 0 | 0 | 0 | 0 | 0 | 0.02 |
| k__Bacteria | p__Proteobacteria | c__Gammaprote | o__Pseudomonadale | f__Moraxellaceae | g__Acinetobacter | s__Acinetobacter_towneri | 0 | 0 | 0 | 0 | 0 | 0 | 0 | 0 | 0 | 0 | 0 | 0 | 0 | 0 | 0 | 0 | 0 | 0 | 0 | 0 | 0 | 0 | 0 | 0 | 0 | 0 | 0 | 0 | 0.19 | 0 | 0 | 0 | 0 | 0 | 0 | 0 | 0 | 0 |
| k__Bacteria | p__Firmicutes | c__Bacilli | o__Lactobacillales | f__Leuconostocaceae | g__Weissella | s__Weissella_minor | 0 | 0 | 0 | 0 | 0 | 0.079 | 0 | 0 | 0 | 0 | 0 | 0 | 0 | 0 | 0 | 0 | 0 | 0 | 0 | 0 | 0 | 0 | 0 | 0 | 0 | 0 | 0 | 0 | 0 | 0 | 0 | 0 | 0.19 | 0 | 0 | 0 | 0 | 0 |
| k__Bacteria | p__Firmicutes | c__Bacilli | o__Bacillales | f__Bacillaceae | g__Bacillus | s__Bacillus_gibsonii | 0 | 0 | 0 | 0 | 0 | 0 | 0 | 0 | 0 | 0 | 0 | 0 | 0 | 0 | 0 | 0 | 0 | 0 | 0 | 0 | 0 | 0 | 0 | 0 | 0 | 0 | 0 | 0 | 0 | 0 | 0 | 0 | 0 | 0 | 0.02 | 0.14 | 0.03 | 0 |
| k__Bacteria | p__Firmicutes | c__Bacilli | o__Lactobacillales | f__Leuconostocaceae | g__Weissella | s__Weissella_cibaria | 0 | 0 | 0 | 0 | 0 | 0 | 0 | 0 | 0 | 0 | 0 | 0 | 0 | 0 | 0 | 0 | 0 | 0 | 0.01 | 0 | 0 | 0 | 0 | 0 | 0 | 0 | 0 | 0 | 0 | 0 | 0 | 0 | 0.13 | 0 | 0 | 0 | 0 | 0 |
| k__Bacteria | p__Proteobacteria | c__Gammaprote | o__Enterobacterales | f__Erwiniaceae | g__Pantoea | s__Pantoea_dispersa | 0 | 0 | 0 | 0 | 0 | 0 | 0 | 0 | 0 | 0 | 0 | 0 | 0 | 0 | 0 | 0 | 0 | 0 | 0 | 0 | 0 | 0 | 0 | 0 | 0 | 0 | 0 | 0 | 0 | 0 | 0 | 0 | 0 | 0 | 0 | 0 | 0 | 0.11 |
| k__Bacteria | p__Firmicutes | c__Bacilli | o__Lactobacillales | f__Lactobacillaceae | g__Lactobacillus | s__Lactobacillus_saniviri | 0 | 0 | 0 | 0 | 0 | 0 | 0 | 0 | 0 | 0 | 0 | 0 | 0 | 0 | 0 | 0 | 0 | 0 | 0 | 0 | 0 | 0 | 0 | 0 | 0 | 0 | 0 | 0 | 0 | 0 | 0 | 0 | 0.08 | 0 | 0 | 0 | 0 | 0 |
| k__Bacteria | p__Firmicutes | c__Bacilli | o__Lactobacillales | f__Lactobacillaceae | g__Lactobacillus | s__Lactobacillus_brevis | 0 | 0 | 0 | 0 | 0 | 0 | 0 | 0 | 0 | 0 | 0 | 0 | 0 | 0 | 0 | 0 | 0 | 0 | 0 | 0 | 0 | 0 | 0 | 0 | 0 | 0 | 0 | 0 | 0 | 0 | 0 | 0 | 0.02 | 0.06 | 0 | 0 | 0 | 0 |
| k__Bacteria | p__Firmicutes | c__Clostridia | o__Clostridiales | f__Ruminococcaceae | g__Ruminococcus | s__Ruminococcus_bicirculans | 0 | 0 | 0 | 0 | 0 | 0.043 | 0 | 0 | 0 | 0 | 0 | 0 | 0 | 0 | 0 | 0 | 0 | 0 | 0 | 0 | 0 | 0 | 0 | 0 | 0 | 0 | 0 | 0 | 0 | 0 | 0 | 0 | 0 | 0 | 0 | 0 | 0 | 0 |
| k__Bacteria | p__Firmicutes | c__Clostridia | o__Clostridiales | f__Lachnospiraceae | g__Lachnospiraceae_uncl | s__Eubacterium_rectale | 0 | 0 | 0 | 0 | 0 | 0.041 | 0 | 0 | 0 | 0 | 0 | 0 | 0 | 0 | 0 | 0 | 0 | 0 | 0 | 0 | 0 | 0 | 0 | 0 | 0 | 0 | 0 | 0 | 0 | 0 | 0 | 0 | 0 | 0 | 0 | 0 | 0 | 0 |
| k__Bacteria | p__Proteobacteria | c__Gammaprote | o__Pseudomonadale | f__Moraxellaceae | g__Acinetobacter | s__Acinetobacter_johnsonii | 0 | 0 | 0 | 0 | 0 | 0 | 0 | 0 | 0 | 0 | 0 | 0 | 0 | 0 | 0 | 0 | 0 | 0 | 0 | 0 | 0 | 0 | 0 | 0 | 0.01 | 0.03 | 0.01 | 0.04 | 0 | 0 | 0 | 0 | 0 | 0 | 0 | 0 | 0 | 0 |
| k__Bacteria | p__Proteobacteria | c__Gammaprote | o__Enterobacterales | f__Enterobacteriaceae | g__Enterobacter | s__Enterobacter_mori | 0 | 0 | 0 | 0 | 0 | 0 | 0 | 0 | 0 | 0 | 0 | 0 | 0 | 0 | 0 | 0 | 0 | 0 | 0 | 0 | 0 | 0 | 0 | 0 | 0 | 0 | 0 | 0.04 | 0 | 0 | 0 | 0 | 0 | 0.04 | 0 | 0 | 0 | 0 |
| k__Bacteria | p__Firmicutes | c__Clostridia | o__Clostridiales | f__Ruminococcaceae | g__Ruminococcus | s__Ruminococcus_bromii | 0 | 0 | 0 | 0 | 0 | 0.038 | 0 | 0 | 0 | 0 | 0 | 0 | 0 | 0 | 0 | 0 | 0 | 0 | 0 | 0 | 0 | 0 | 0 | 0 | 0 | 0 | 0 | 0 | 0 | 0 | 0 | 0 | 0 | 0 | 0 | 0 | 0 | 0 |
| k__Bacteria | p__Bacteroidetes | c__Bacteroidia | o__Bacteroidales | f__Bacteroidaceae | g__Bacteroides | s__Bacteroides_stercoris | 0 | 0 | 0 | 0 | 0 | 0.03 | 0 | 0 | 0 | 0 | 0 | 0 | 0 | 0 | 0 | 0 | 0 | 0 | 0 | 0 | 0 | 0 | 0 | 0 | 0 | 0 | 0 | 0 | 0 | 0 | 0 | 0 | 0 | 0 | 0 | 0 | 0 | 0 |
| k__Bacteria | p__Actinobacteria | c__Actinobacter | o__Bifidobacteriales | f__Bifidobacteriaceae | g__Bifidobacterium | s__Bifidobacterium_longum | 0 | 0 | 0 | 0 | 0 | 0.023 | 0 | 0 | 0 | 0 | 0 | 0 | 0 | 0 | 0 | 0 | 0 | 0 | 0 | 0 | 0 | 0 | 0 | 0 | 0 | 0 | 0 | 0 | 0 | 0 | 0 | 0 | 0 | 0 | 0 | 0 | 0 | 0 |
| k__Bacteria | p__Firmicutes | c__Bacilli | o__Lactobacillales | f__Lactobacillaceae | g__Lactobacillus | s__Lactobacillus_pontis | 0 | 0 | 0 | 0 | 0 | 0 | 0 | 0 | 0 | 0 | 0 | 0 | 0 | 0 | 0 | 0 | 0 | 0 | 0 | 0 | 0 | 0 | 0 | 0 | 0 | 0.02 | 0 | 0.02 | 0 | 0 | 0 | 0 | 0 | 0 | 0 | 0 | 0 | 0 |
| k__Bacteria | p__Proteobacteria | c__Gammaprote | o__Enterobacterales | f__Erwiniaceae | g__Mixta | s__Mixta_calida | 0 | 0 | 0 | 0 | 0 | 0 | 0 | 0 | 0 | 0 | 0 | 0 | 0 | 0 | 0 | 0 | 0 | 0 | 0 | 0 | 0 | 0 | 0 | 0 | 0 | 0 | 0 | 0 | 0 | 0 | 0 | 0 | 0 | 0 | 0.02 | 0 | 0 | 0.01 |
| k__Bacteria | p__Actinobacteria | c__Actinobacter | o__Bifidobacteriales | f__Bifidobacteriaceae | g__Bifidobacterium | s__Bifidobacterium_adolescentis | 0 | 0 | 0 | 0 | 0 | 0.021 | 0 | 0 | 0 | 0 | 0 | 0 | 0 | 0 | 0 | 0 | 0 | 0 | 0 | 0 | 0 | 0 | 0 | 0 | 0 | 0 | 0 | 0 | 0 | 0 | 0 | 0 | 0 | 0 | 0 | 0 | 0 | 0 |
| k__Bacteria | p__Bacteroidetes | c__Bacteroidia | o__Bacteroidales | f__Bacteroidaceae | g__Bacteroides | s__Bacteroides_vulgatus | 0 | 0 | 0 | 0 | 0 | 0.02 | 0 | 0 | 0 | 0 | 0 | 0 | 0 | 0 | 0 | 0 | 0 | 0 | 0 | 0 | 0 | 0 | 0 | 0 | 0 | 0 | 0 | 0 | 0 | 0 | 0 | 0 | 0 | 0 | 0 | 0 | 0 | 0 |
| k__Bacteria | p__Firmicutes | c__Bacilli | o__Bacillales | f__Bacillaceae | g__Bacillus | s__Bacillus_rhizosphaerae | 0 | 0 | 0 | 0 | 0 | 0 | 0 | 0 | 0 | 0 | 0 | 0 | 0 | 0 | 0 | 0 | 0 | 0 | 0 | 0 | 0 | 0 | 0 | 0 | 0 | 0 | 0 | 0 | 0 | 0 | 0 | 0 | 0 | 0 | 0 | 0 | 0.02 | 0 |
| k__Bacteria | p__Firmicutes | c__Clostridia | o__Clostridiales | f__Ruminococcaceae | g__Faecalibacterium | s__Faecalibacterium_prausnitzii | 0 | 0 | 0 | 0 | 0 | 0.017 | 0 | 0 | 0 | 0 | 0 | 0 | 0 | 0 | 0 | 0 | 0 | 0 | 0 | 0 | 0 | 0 | 0 | 0 | 0 | 0 | 0 | 0 | 0 | 0 | 0 | 0 | 0 | 0 | 0 | 0 | 0 | 0 |
| k__Bacteria | p__Proteobacteria | c__Gammaprote | o__Enterobacterales | f__Enterobacteriaceae | g__Klebsiella | s__Klebsiella_pneumoniae | 0 | 0 | 0 | 0 | 0 | 0 | 0 | 0 | 0 | 0 | 0 | 0 | 0 | 0 | 0 | 0 | 0 | 0 | 0 | 0 | 0 | 0 | 0 | 0 | 0 | 0 | 0 | 0 | 0 | 0 | 0 | 0 | 0 | 0 | 0 | 0 | 0 | 0.01 |
| k__Bacteria | p__Firmicutes | c__Bacilli | o__Bacillales | f__Staphylococcaceae | g__Staphylococcus | s__Staphylococcus_argenteus | 0 | 0 | 0 | 0 | 0 | 0 | 0 | 0 | 0 | 0 | 0 | 0 | 0 | 0 | 0 | 0 | 0 | 0 | 0 | 0 | 0 | 0 | 0 | 0 | 0 | 0 | 0 | 0 | 0 | 0 | 0 | 0 | 0.01 | 0 | 0 | 0 | 0 | 0 |
| k__Bacteria | p__Firmicutes | c__Bacilli | o__Bacillales | f__Planococcaceae | g__Jeotgalibacillus | s__Jeotgalibacillus_marinus | 0 | 0 | 0 | 0 | 0 | 0 | 0 | 0 | 0 | 0 | 0 | 0 | 0 | 0 | 0 | 0 | 0 | 0 | 0 | 0 | 0 | 0 | 0 | 0 | 0 | 0 | 0 | 0 | 0 | 0 | 0 | 0 | 0 | 0 | 0 | 0.01 | 0 | 0 |
| k__Bacteria | p__Firmicutes | c__Bacilli | o__Lactobacillales | f__Leuconostocaceae | g__Leuconostoc | s__Leuconostoc_gelidum | 0 | 0 | 0 | 0 | 0 | 0 | 0 | 0 | 0 | 0 | 0 | 0 | 0 | 0 | 0 | 0 | 0 | 0 | 0 | 0 | 0 | 0 | 0 | 0 | 0 | 0 | 0 | 0 | 0 | 0 | 0 | 0 | 0.01 | 0 | 0 | 0 | 0 | 0 |
| k__Bacteria | p__Firmicutes | c__Clostridia | o__Clostridiales | f__Eubacteriaceae | g__Eubacterium | s__Eubacterium_hallii | 0 | 0 | 0 | 0 | 0 | 0.01 | 0 | 0 | 0 | 0 | 0 | 0 | 0 | 0 | 0 | 0 | 0 | 0 | 0 | 0 | 0 | 0 | 0 | 0 | 0 | 0 | 0 | 0 | 0 | 0 | 0 | 0 | 0 | 0 | 0 | 0 | 0 | 0 |
| k__Bacteria | p__Firmicutes | c__Bacilli | o__Lactobacillales | f__Leuconostocaceae | g__Leuconostoc | s__Leuconostoc_mesenteroides | 0 | 0 | 0 | 0 | 0 | 0 | 0 | 0 | 0 | 0 | 0 | 0 | 0 | 0 | 0 | 0 | 0 | 0 | 0 | 0 | 0 | 0 | 0 | 0 | 0 | 0 | 0 | 0 | 0 | 0 | 0 | 0 | 0.01 | 0 | 0 | 0 | 0 | 0 |
| k__Bacteria | p__Firmicutes | c__Bacilli | o__Lactobacillales | f__Lactobacillaceae | g__Lactobacillus | s__Lactobacillus_pentosus | 0 | 0 | 0 | 0 | 0 | 0 | 0 | 0 | 0 | 0 | 0 | 0 | 0 | 0 | 0 | 0 | 0 | 0 | 0 | 0 | 0 | 0 | 0 | 0 | 0 | 0 | 0 | 0 | 0 | 0 | 0 | 0 | 0.01 | 0 | 0 | 0 | 0 | 0 |
| k__Bacteria | p__Firmicutes | c__Bacilli | o__Bacillales | f__Bacillaceae | g__Bacillus | s__Bacillus_clausii | 0 | 0 | 0 | 0 | 0 | 0 | 0 | 0 | 0 | 0 | 0 | 0 | 0 | 0 | 0 | 0 | 0 | 0 | 0 | 0 | 0 | 0 | 0 | 0 | 0 | 0 | 0 | 0 | 0 | 0 | 0 | 0 | 0 | 0 | 0 | 0 | 0.01 | 0 |
| k__Bacteria | p__Firmicutes | c__Clostridia | o__Clostridiales | f__Lachnospiraceae | g__Fusicatenibacter | s__Fusicatenibacter_saccharivorans | 0 | 0 | 0 | 0 | 0 | 0.008 | 0 | 0 | 0 | 0 | 0 | 0 | 0 | 0 | 0 | 0 | 0 | 0 | 0 | 0 | 0 | 0 | 0 | 0 | 0 | 0 | 0 | 0 | 0 | 0 | 0 | 0 | 0 | 0 | 0 | 0 | 0 | 0 |
| k__Bacteria | p__Bacteroidetes | c__Bacteroidia | o__Bacteroidales | f__Bacteroidaceae | g__Bacteroides | s__Bacteroides_uniformis | 0 | 0 | 0 | 0 | 0 | 0.007 | 0 | 0 | 0 | 0 | 0 | 0 | 0 | 0 | 0 | 0 | 0 | 0 | 0 | 0 | 0 | 0 | 0 | 0 | 0 | 0 | 0 | 0 | 0 | 0 | 0 | 0 | 0 | 0 | 0 | 0 | 0 | 0 |
| k__Bacteria | p__Firmicutes | c__Firmicutes_u | o__Firmicutes_unclas | f__Firmicutes_unclassifi | g__Firmicutes_unclassifie | s__Firmicutes_bacterium_CAG_83 | 0 | 0 | 0 | 0 | 0 | 0.007 | 0 | 0 | 0 | 0 | 0 | 0 | 0 | 0 | 0 | 0 | 0 | 0 | 0 | 0 | 0 | 0 | 0 | 0 | 0 | 0 | 0 | 0 | 0 | 0 | 0 | 0 | 0 | 0 | 0 | 0 | 0 | 0 |
| k__Bacteria | p__Firmicutes | c__Bacilli | o__Lactobacillales | f__Lactobacillaceae | g__Lactobacillus | s__Lactobacillus_paraplantarum | 0 | 0 | 0 | 0 | 0 | 0 | 0 | 0 | 0 | 0 | 0 | 0 | 0 | 0 | 0 | 0 | 0 | 0 | 0 | 0 | 0 | 0 | 0 | 0 | 0 | 0 | 0 | 0 | 0 | 0 | 0 | 0 | 0.01 | 0 | 0 | 0 | 0 | 0 |
| k__Bacteria | p__Firmicutes | c__Clostridia | o__Clostridiales | f__Lachnospiraceae | g__Blautia | s__Blautia_wexlerae | 0 | 0 | 0 | 0 | 0 | 0.005 | 0 | 0 | 0 | 0 | 0 | 0 | 0 | 0 | 0 | 0 | 0 | 0 | 0 | 0 | 0 | 0 | 0 | 0 | 0 | 0 | 0 | 0 | 0 | 0 | 0 | 0 | 0 | 0 | 0 | 0 | 0 | 0 |
| k__Bacteria | p__Firmicutes | c__Bacilli | o__Bacillales | f__Bacillaceae | g__Bacillus | s__Bacillus_cellulasensis | 0 | 0 | 0 | 0 | 0 | 0 | 0 | 0 | 0 | 0 | 0 | 0 | 0 | 0 | 0 | 0 | 0 | 0 | 0 | 0 | 0 | 0 | 0 | 0 | 0 | 0 | 0 | 0 | 0 | 0 | 0 | 0 | 0 | 0 | 0 | 0 | 0 | 0 |
| k__Bacteria | p__Actinobacteria | c__Actinobacter | o__Bifidobacteriales | f__Bifidobacteriaceae | g__Bifidobacterium | s__Bifidobacterium_pseudocatenulatum | 0 | 0 | 0 | 0 | 0 | 0.002 | 0 | 0 | 0 | 0 | 0 | 0 | 0 | 0 | 0 | 0 | 0 | 0 | 0 | 0 | 0 | 0 | 0 | 0 | 0 | 0 | 0 | 0 | 0 | 0 | 0 | 0 | 0 | 0 | 0 | 0 | 0 | 0 |
| k__Bacteria | p__Actinobacteria | c__Actinobacter | o__Micrococcales | f__Micrococcaceae | g__Kocuria | s__Kocuria_kristinae | 0 | 0 | 0 | 0 | 0 | 0 | 0 | 0 | 0 | 0 | 0 | 0 | 0 | 0 | 0 | 0 | 0 | 0 | 0 | 0 | 0 | 0 | 0 | 0 | 0 | 0 | 0 | 0 | 0 | 0 | 0 | 0 | 0 | 0 | 0 | 0 | 0 | 0 |
| k__Bacteria | p__Bacteroidetes | c__Bacteroidia | o__Bacteroidales | f__Bacteroidaceae | g__Bacteroides | s__Bacteroides_ovatus | 0 | 0 | 0 | 0 | 0 | 0.002 | 0 | 0 | 0 | 0 | 0 | 0 | 0 | 0 | 0 | 0 | 0 | 0 | 0 | 0 | 0 | 0 | 0 | 0 | 0 | 0 | 0 | 0 | 0 | 0 | 0 | 0 | 0 | 0 | 0 | 0 | 0 | 0 |
| k__Bacteria | p__Firmicutes | c__Clostridia | o__Clostridiales | f__Lachnospiraceae | g__Roseburia | s__Roseburia_faecis | 0 | 0 | 0 | 0 | 0 | 0.002 | 0 | 0 | 0 | 0 | 0 | 0 | 0 | 0 | 0 | 0 | 0 | 0 | 0 | 0 | 0 | 0 | 0 | 0 | 0 | 0 | 0 | 0 | 0 | 0 | 0 | 0 | 0 | 0 | 0 | 0 | 0 | 0 |
| k__Bacteria | p__Actinobacteria | c__Coriobacterii | o__Eggerthellales | f__Eggerthellaceae | g__Eggerthella | s__Eggerthella_lenta | 0 | 0 | 0 | 0 | 0 | 0.002 | 0 | 0 | 0 | 0 | 0 | 0 | 0 | 0 | 0 | 0 | 0 | 0 | 0 | 0 | 0 | 0 | 0 | 0 | 0 | 0 | 0 | 0 | 0 | 0 | 0 | 0 | 0 | 0 | 0 | 0 | 0 | 0 |
| k__Bacteria | p__Firmicutes | c__Clostridia | o__Clostridiales | f__Lachnospiraceae | g__Dorea | s__Dorea_longicatena | 0 | 0 | 0 | 0 | 0 | 0.002 | 0 | 0 | 0 | 0 | 0 | 0 | 0 | 0 | 0 | 0 | 0 | 0 | 0 | 0 | 0 | 0 | 0 | 0 | 0 | 0 | 0 | 0 | 0 | 0 | 0 | 0 | 0 | 0 | 0 | 0 | 0 | 0 |
| k__Bacteria | p__Actinobacteria | c__Coriobacterii | o__Coriobacteriales | f__Coriobacteriaceae | g__Collinsella | s__Collinsella_intestinalis | 0 | 0 | 0 | 0 | 0 | 0.001 | 0 | 0 | 0 | 0 | 0 | 0 | 0 | 0 | 0 | 0 | 0 | 0 | 0 | 0 | 0 | 0 | 0 | 0 | 0 | 0 | 0 | 0 | 0 | 0 | 0 | 0 | 0 | 0 | 0 | 0 | 0 | 0 |
| k__Bacteria | p__Firmicutes | c__Clostridia | o__Clostridiales | f__Ruminococcaceae | g__Ruminococcus | s__Ruminococcus_callidus | 0 | 0 | 0 | 0 | 0 | 4E-04 | 0 | 0 | 0 | 0 | 0 | 0 | 0 | 0 | 0 | 0 | 0 | 0 | 0 | 0 | 0 | 0 | 0 | 0 | 0 | 0 | 0 | 0 | 0 | 0 | 0 | 0 | 0 | 0 | 0 | 0 | 0 | 0 |
| k__Bacteria | p__Firmicutes | c__Clostridia | o__Clostridiales | f__Eubacteriaceae | g__Eubacterium | s__Eubacterium_sp_CAG_38 | 0 | 0 | 0 | 0 | 0 | 4E-04 | 0 | 0 | 0 | 0 | 0 | 0 | 0 | 0 | 0 | 0 | 0 | 0 | 0 | 0 | 0 | 0 | 0 | 0 | 0 | 0 | 0 | 0 | 0 | 0 | 0 | 0 | 0 | 0 | 0 | 0 | 0 | 0 |
| k__Bacteria | p__Bacteroidetes | c__Bacteroidia | o__Bacteroidales | f__Bacteroidaceae | g__Bacteroides | s__Bacteroides_caccae | 0 | 0 | 0 | 0 | 0 | 3E-04 | 0 | 0 | 0 | 0 | 0 | 0 | 0 | 0 | 0 | 0 | 0 | 0 | 0 | 0 | 0 | 0 | 0 | 0 | 0 | 0 | 0 | 0 | 0 | 0 | 0 | 0 | 0 | 0 | 0 | 0 | 0 | 0 |

**Table S4. 'ychF', abundance table using the ychF gene as a marker;**

| Species | F.1 | F1.3 | F2.3 | F3.3 | L.1 | L1.3 | L2.3 | L3.3 | NF.1 | NF1.3 | NF2.3 | NF3.3 | NL.1 | NL1.3 | NL2.3 | NL3.3 | P.1 | P1.3 | P2.3 | P3.3 | NP.1 | NP1.3 | NP2.3 | NP3.3 | R.1 | R1.3 | R2.3 | R3.3 | S.1 | S1.3 | S2.3 | S3.3 | HB | MS | PB | PR | SY | TS |
|---|---|---|---|---|---|---|---|---|---|---|---|---|---|---|---|---|---|---|---|---|---|---|---|---|---|---|---|---|---|---|---|---|---|---|---|---|---|---|
| *Bacillus velezensis* | 65.4 | 1.56 | 1.29 | 0.9 | 82.2 | 2.49 | 1.9 | 3.92 | 65.8 | 3.49 | 2.5 | 2.64 | 73.3 | 2.06 | 2.35 | 2.96 | 67.8 | 3.2 | 2.1 | 2.15 | 100 | 4.34 | 3.64 | 2.43 | 76.5 | 83.3 | 78.3 | 80 | 100 | 0.52 | 0.43 | 0 | 0 | 13.9 | 0 | 0 | 0 | 2.35 |
| *Pediococcus pentosaceus* | 0 | 14.1 | 40.8 | 36.2 | 0 | 3.41 | 13.9 | 35.7 | 0 | 83.8 | 89.3 | 93.6 | 0 | 94.6 | 93.9 | 92.1 | 0 | 3.45 | 28.7 | 35.1 | 0 | 75.8 | 58.2 | 84.1 | 0 | 0 | 0 | 0 | 0 | 0 | 0 | 0 | 76.5 | 0.92 | 0 | 0 | 0 | 1.62 |
| *Staphylococcus pasteuri* | 0 | 78.4 | 52.5 | 59.6 | 0 | 86.3 | 70.6 | 42.6 | 0 | 3.33 | 2.58 | 0.28 | 0 | 0 | 0.81 | 0.97 | 0 | 81.2 | 58.6 | 54.6 | 0 | 12.1 | 22.6 | 8.56 | 0 | 0 | 0 | 0 | 0 | 7.8 | 11.9 | 8.74 | 9.36 | 2.4 | 0 | 0 | 1.96 | 1.39 |
| *Staphylococcus epidermidis* | 0 | 4.69 | 2.77 | 1.77 | 0 | 6.57 | 11.8 | 11.7 | 0 | 4.29 | 2.48 | 0.79 | 0 | 0.25 | 0.31 | 0.34 | 0 | 7.36 | 6.83 | 5.71 | 0 | 4.56 | 10 | 1.91 | 0 | 0 | 0 | 0 | 0 | 79.3 | 84.2 | 72.1 | 0 | 10.2 | 1.49 | 49.7 | 0 | 1.98 |
| *Pediococcus acidilactici* | 0 | 0 | 0 | 0 | 0 | 0 | 0 | 0 | 0 | 0 | 0 | 0 | 0 | 0 | 0 | 0 | 0 | 0 | 0 | 0 | 0 | 0 | 0 | 0 | 0 | 0 | 0 | 0 | 0 | 0 | 0 | 0 | 0 | 0 | 67.9 | 7.83 | 0 | 25.1 |
| *Enterococcus faecium* | 0 | 0 | 0 | 0 | 0 | 0 | 0 | 0 | 0 | 0 | 0 | 0 | 0 | 0 | 0 | 0 | 0 | 0 | 0 | 0 | 0 | 0 | 0 | 0 | 0 | 0 | 0 | 0 | 0 | 0 | 0 | 0 | 0 | 1.18 | 0 | 0 | 67.5 | 0.97 |
| *Staphylococcus warneri* | 0 | 0.57 | 1.88 | 0.84 | 0 | 0 | 1.06 | 2.47 | 0 | 0 | 0 | 0 | 0 | 0 | 0 | 0 | 0 | 0.6 | 1.34 | 0 | 0 | 0 | 0 | 0 | 0 | 0 | 0 | 0 | 0 | 0 | 0 | 0 | 3.26 | 55.8 | 28.3 | 19.6 | 10.3 | 54.8 |
| *Cronobacter malonaticus* | 22.8 | 0.39 | 0 | 0.25 | 17.8 | 0.73 | 0 | 0.75 | 9.76 | 0.3 | 0.16 | 0.29 | 6.99 | 0.33 | 0.23 | 0.34 | 20 | 2.73 | 0.61 | 0.54 | 0 | 0.27 | 0 | 0 | 8.41 | 10.2 | 10.9 | 9.27 | 0 | 0 | 0 | 0 | 0 | 0 | 0 | 0 | 0 | 0 |
| *Tetragenococcus halophilus* | 0 | 0 | 0 | 0 | 0 | 0 | 0 | 0 | 0 | 1.71 | 0 | 0 | 0 | 0 | 0 | 0 | 0 | 0.3 | 0 | 0 | 0 | 0 | 0 | 0 | 0 | 0 | 0 | 0 | 0 | 12.4 | 3.47 | 18.8 | 0 | 0 | 0 | 0 | 0 | 0 |
| *Bacillus subtilis* | 0 | 0 | 0 | 0 | 0 | 0 | 0 | 0 | 0 | 0 | 0 | 0 | 0 | 0 | 0 | 0 | 0 | 0 | 0 | 0 | 0 | 0 | 0 | 0 | 0 | 0 | 0 | 0 | 0 | 0 | 0 | 0 | 0.55 | 0 | 1.24 | 16.4 | 1.08 | 0 |
| *Lactiplantibacillus plantarum* | 0 | 0 | 0.71 | 0.51 | 0 | 0 | 0 | 1.88 | 0 | 2.09 | 2.36 | 2.03 | 0 | 2.32 | 2.16 | 2.84 | 0 | 0 | 1.07 | 0.72 | 0 | 1.9 | 1.33 | 2.08 | 0 | 0 | 0 | 0 | 0 | 0 | 0 | 0 | 0.86 | 15.6 | 0 | 0 | 0 | 2.33 |
| *Weissella hellenica* | 0 | 0 | 0 | 0 | 0 | 0 | 0 | 0 | 0 | 0 | 0 | 0 | 0 | 0 | 0 | 0 | 0 | 0 | 0 | 0 | 0 | 0 | 0 | 0 | 0 | 0 | 0 | 0 | 0 | 0 | 0 | 0 | 0 | 0 | 0 | 0 | 12.8 | 0 |
| *Enterobacter cloacae complex* | 11.8 | 0 | 0 | 0 | 0 | 0.49 | 0.34 | 0 | 12.2 | 0.37 | 0.16 | 0.31 | 12.7 | 0.41 | 0.23 | 0.46 | 12.2 | 0.78 | 0.84 | 0.68 | 0 | 0.45 | 0.32 | 0.37 | 10.3 | 6.52 | 10.8 | 9.49 | 0 | 0 | 0 | 0 | 0.31 | 0 | 0 | 0 | 4.31 | 0 |
| *Bacillus cereus* | 0 | 0 | 0 | 0 | 0 | 0 | 0 | 0 | 12.2 | 0.3 | 0 | 0 | 6.99 | 0 | 0 | 0 | 0 | 0 | 0 | 0 | 0 | 0 | 0 | 0 | 0 | 0 | 0 | 0 | 0 | 0 | 0 | 0 | 0 | 0 | 0 | 0 | 0 | 0 |
| *Enterococcus faecalis* | 0 | 0 | 0 | 0 | 0 | 0 | 0 | 0 | 0 | 0 | 0 | 0 | 0 | 0 | 0 | 0 | 0 | 0 | 0 | 0 | 0 | 0 | 0 | 0 | 0 | 0 | 0 | 0 | 0 | 0 | 0 | 0 | 0 | 0 | 0 | 0 | 0 | 8.98 |
| *Fructilactobacillus sanfranciscensis* | 0 | 0 | 0 | 0 | 0 | 0 | 0 | 0 | 0 | 0 | 0 | 0 | 0 | 0 | 0 | 0 | 0 | 0 | 0 | 0 | 0 | 0 | 0 | 0 | 4.73 | 0 | 0 | 0 | 0 | 0 | 0 | 0 | 0 | 0 | 0 | 0 | 0 | 0 |
| *Debaryomyces hansenii* | 0 | 0 | 0 | 0 | 0 | 0 | 0 | 0 | 0 | 0 | 0 | 0 | 0 | 0 | 0 | 0 | 0 | 0 | 0 | 0 | 0 | 0 | 0 | 0 | 0 | 0 | 0 | 0 | 0 | 0 | 0 | 0 | 0 | 0 | 0 | 4.44 | 0 | 0 |
| *Latilactobacillus sakei* | 0 | 0 | 0 | 0 | 0 | 0 | 0 | 0 | 0 | 0 | 0 | 0 | 0 | 0 | 0 | 0 | 0 | 0 | 0 | 0 | 0 | 0 | 0 | 0 | 0 | 0 | 0 | 0 | 0 | 0 | 0 | 0 | 3.85 | 0 | 0 | 0 | 0 | 0 |
| *Staphylococcus simulans* | 0 | 0 | 0 | 0 | 0 | 0 | 0 | 0 | 0 | 0 | 0 | 0 | 0 | 0 | 0 | 0 | 0 | 0 | 0 | 0 | 0 | 0.42 | 2.85 | 0.55 | 0 | 0 | 0 | 0 | 0 | 0 | 0 | 0 | 0 | 0 | 0 | 0 | 0 | 0 |
| *Staphylococcus aureus* | 0 | 0 | 0 | 0 | 0 | 0 | 0.38 | 0.59 | 0 | 0 | 0 | 0 | 0 | 0 | 0 | 0 | 0 | 0 | 0 | 0 | 0 | 0 | 0 | 0 | 0 | 0 | 0 | 0 | 0 | 0 | 0 | 0 | 2.58 | 0 | 0 | 0 | 0 | 0 |
| *Staphylococcus lugdunensis* | 0 | 0 | 0 | 0 | 0 | 0 | 0 | 0 | 0 | 0 | 0 | 0 | 0 | 0 | 0 | 0 | 0 | 0.4 | 0 | 0 | 0 | 0 | 0 | 0 | 0 | 0 | 0 | 0 | 0 | 0 | 0 | 0 | 0 | 0 | 0 | 2.09 | 0 | 0 |
| *Weissella confusa* | 0 | 0 | 0 | 0 | 0 | 0 | 0 | 0 | 0 | 0 | 0 | 0 | 0 | 0 | 0 | 0 | 0 | 0 | 0 | 0 | 0 | 0 | 0 | 0 | 0 | 0 | 0 | 0 | 0 | 0 | 0 | 0 | 0.23 | 0 | 0 | 0 | 1.17 | 0 |
| *Staphylococcus taiwanensis* | 0 | 0 | 0 | 0 | 0 | 0 | 0 | 0 | 0 | 0 | 0 | 0 | 0 | 0 | 0 | 0 | 0 | 0 | 0 | 0 | 0 | 0 | 0 | 0 | 0 | 0 | 0 | 0 | 0 | 0 | 0 | 0 | 0 | 0 | 1.02 | 0 | 0.52 | 0 |
| *Weissella viridescens* | 0 | 0 | 0 | 0 | 0 | 0 | 0 | 0 | 0 | 0 | 0 | 0 | 0 | 0 | 0 | 0 | 0 | 0 | 0 | 0 | 0 | 0 | 0 | 0 | 0 | 0 | 0 | 0 | 0 | 0 | 0 | 0 | 0.96 | 0 | 0 | 0 | 0 | 0 |
| *Weissella minor* | 0 | 0 | 0 | 0 | 0 | 0 | 0 | 0 | 0 | 0.3 | 0 | 0 | 0 | 0 | 0 | 0 | 0 | 0 | 0 | 0 | 0 | 0 | 0 | 0 | 0 | 0 | 0 | 0 | 0 | 0 | 0 | 0 | 0.86 | 0 | 0 | 0 | 0 | 0 |
| *Zygosaccharomyces bailii* | 0 | 0 | 0 | 0 | 0 | 0 | 0 | 0 | 0 | 0 | 0 | 0 | 0 | 0 | 0 | 0 | 0 | 0 | 0 | 0 | 0 | 0 | 0 | 0 | 0 | 0 | 0 | 0.63 | 0 | 0 | 0 | 0 | 0 | 0 | 0 | 0 | 0 | 0 |
| *[Candida] glabrata* | 0 | 0 | 0 | 0 | 0 | 0 | 0 | 0 | 0 | 0 | 0 | 0 | 0 | 0 | 0 | 0 | 0 | 0 | 0 | 0 | 0 | 0 | 0 | 0 | 0 | 0 | 0 | 0.62 | 0 | 0 | 0 | 0 | 0 | 0 | 0 | 0 | 0 | 0 |
| *Staphylococcus saprophyticus* | 0 | 0 | 0 | 0 | 0 | 0 | 0 | 0 | 0 | 0 | 0.4 | 0 | 0 | 0 | 0 | 0 | 0 | 0 | 0 | 0 | 0 | 0.18 | 0.57 | 0 | 0 | 0 | 0 | 0 | 0 | 0 | 0 | 0 | 0 | 0 | 0 | 0 | 0 | 0.45 |
| *Latilactobacillus curvatus* | 0 | 0.29 | 0 | 0 | 0 | 0 | 0 | 0 | 0 | 0 | 0 | 0 | 0 | 0 | 0 | 0 | 0 | 0 | 0 | 0.5 | 0 | 0 | 0.38 | 0 | 0 | 0 | 0 | 0 | 0 | 0 | 0 | 0 | 0 | 0 | 0 | 0 | 0 | 0 |
| *Saccharomyces cerevisiae* | 0 | 0 | 0 | 0 | 0 | 0 | 0 | 0 | 0 | 0 | 0 | 0 | 0 | 0 | 0 | 0 | 0 | 0 | 0 | 0 | 0 | 0 | 0 | 0 | 0 | 0 | 0 | 0 | 0 | 0 | 0 | 0 | 0.47 | 0 | 0 | 0 | 0 | 0 |
| *Serratia liquefaciens* | 0 | 0 | 0 | 0 | 0 | 0 | 0 | 0 | 0 | 0 | 0 | 0 | 0 | 0 | 0 | 0 | 0 | 0 | 0 | 0 | 0 | 0 | 0 | 0 | 0 | 0 | 0 | 0 | 0 | 0 | 0 | 0 | 0 | 0 | 0 | 0 | 0.39 | 0 |
| *Staphylococcus xylosus* | 0 | 0 | 0 | 0 | 0 | 0 | 0 | 0.39 | 0 | 0 | 0 | 0 | 0 | 0 | 0 | 0 | 0 | 0 | 0 | 0 | 0 | 0 | 0 | 0 | 0 | 0 | 0 | 0 | 0 | 0 | 0 | 0 | 0 | 0 | 0 | 0 | 0 | 0 |
| *Weissella cibaria* | 0 | 0 | 0 | 0 | 0 | 0 | 0 | 0 | 0 | 0 | 0 | 0 | 0 | 0 | 0 | 0 | 0 | 0 | 0 | 0 | 0 | 0 | 0 | 0 | 0 | 0 | 0 | 0 | 0 | 0 | 0 | 0.38 | 0 | 0 | 0 | 0 | 0 | 0 |
| *Serratia marcescens* | 0 | 0 | 0 | 0 | 0 | 0 | 0 | 0 | 0 | 0 | 0 | 0 | 0 | 0 | 0 | 0 | 0 | 0 | 0 | 0 | 0 | 0 | 0 | 0 | 0 | 0 | 0 | 0 | 0 | 0 | 0 | 0 | 0.18 | 0 | 0 | 0 | 0 | 0 |

**Table S4. 'leuS', abundance table using the leuS gene as a marker. The colour code indicates relative abundance, ranging from red (low abundance) to green (high abundance).**

| Species | F.1 | F1.3 | F2.3 | F3.3 | L.1 | L1.3 | L2.3 | L3.3 | NF.1 | NF1.3 | NF2.3 | NF3.3 | NL.1 | NL1.3 | NL2.3 | NL3.3 | P.1 | P1.3 | P2.3 | P3.3 | NP.1 | NP1.3 | NP2.3 | NP3.3 | R.1 | R1.3 | R2.3 | R3.3 | S.1 | S1.3 | S2.3 | S3.3 | HB | MS | PB | PR | SY | TS |
|---|---|---|---|---|---|---|---|---|---|---|---|---|---|---|---|---|---|---|---|---|---|---|---|---|---|---|---|---|---|---|---|---|---|---|---|---|---|---|
| *Pediococcus pentosaceus* | 0 | 16.6 | 40.4 | 39.8 | 0 | 4.22 | 17 | 37.6 | 0 | 83.6 | 87.6 | 92.4 | 0 | 93.5 | 92.9 | 91.6 | 0 | 4.24 | 29.4 | 39.1 | 0 | 77.9 | 60.5 | 84.7 | 0 | 0 | 0 | 0 | 0 | 0 | 0 | 0 | 79.6 | 0 | 0 | 0 | 0 | 1.73 |
| *Staphylococcus pasteuri* | 0 | 79 | 53.9 | 55.5 | 0 | 90.8 | 69.9 | 36.8 | 0 | 2.74 | 2.5 | 0.38 | 0 | 0 | 0.71 | 0.84 | 0 | 81.3 | 56.5 | 51.5 | 0 | 10.3 | 21.3 | 7.59 | 0 | 0 | 0 | 0 | 0 | 5.73 | 8.8 | 8.04 | 6.24 | 2.09 | 0 | 0 | 1.12 | 2.6 |
| *Staphylococcus epidermidis* | 0 | 0 | 2.91 | 2.01 | 0 | 0 | 8.56 | 12 | 0 | 4.35 | 2.82 | 0.74 | 0 | 0.31 | 0 | 0.42 | 0 | 6.82 | 7.4 | 4.93 | 0 | 4.85 | 9.31 | 1.96 | 0 | 0 | 0 | 0 | 0 | 83.4 | 88.5 | 73.5 | 0.41 | 13.1 | 0.91 | 36 | 0.79 | 3.23 |
| *Bacillus velezensis* | 46.9 | 2.18 | 1.08 | 1.15 | 56.7 | 3.5 | 2.65 | 4.67 | 68.5 | 4.03 | 3.16 | 3.34 | 68.5 | 2.55 | 2.92 | 3.46 | 66.8 | 4.78 | 2.73 | 2.43 | 63.3 | 3.66 | 4.21 | 2.94 | 61.3 | 73.8 | 78.5 | 82.4 | 39.3 | 0 | 0 | 0 | 0 | 21.4 | 0.27 | 0 | 0 | 2.17 |
| *Enterococcus faecium* | 0 | 0 | 0 | 0 | 0 | 0 | 0 | 0 | 0 | 0 | 0 | 0 | 0 | 0 | 0 | 0 | 0 | 0 | 0 | 0 | 0 | 0 | 0 | 0 | 0 | 0 | 0 | 0 | 0 | 0 | 0 | 0 | 0 | 0.76 | 0 | 0 | 75.3 | 1.12 |
| *Pediococcus acidilactici* | 0 | 0 | 0 | 0 | 0 | 0 | 0 | 0 | 0 | 0 | 0 | 0 | 0 | 0 | 0 | 0 | 0 | 0 | 0 | 0 | 0 | 0 | 0 | 0 | 0 | 0 | 0 | 0 | 0 | 0 | 0 | 0 | 0 | 0 | 71.5 | 6.8 | 0 | 28.3 |
| *Aspergillus oryzae* | 23.4 | 0 | 0 | 0 | 31.8 | 0 | 0 | 0 | 0 | 0 | 0 | 0 | 10.8 | 0 | 0 | 0 | 0 | 0 | 0 | 0 | 20 | 0.61 | 0 | 0 | 21 | 7.21 | 9.12 | 7.62 | 60.7 | 0 | 0 | 0 | 0 | 0 | 0 | 27.1 | 0 | 0 |
| *Enterobacter cloacae complex* | 10 | 0 | 0 | 0 | 0 | 0.77 | 0 | 0 | 14.4 | 0.25 | 0.4 | 0.4 | 13.5 | 0.81 | 0.62 | 0.55 | 11.4 | 1.26 | 1 | 0.67 | 16.7 | 1.1 | 0 | 0.48 | 10.6 | 10.1 | 0 | 0 | 0 | 0 | 0 | 0 | 0 | 0 | 0 | 0 | 0 | 0 |
| *Staphylococcus warneri* | 0 | 1.44 | 1.05 | 0.73 | 0 | 0 | 1.05 | 2.9 | 0 | 0 | 0 | 0 | 0 | 0 | 0 | 0 | 0 | 0.62 | 1.05 | 0 | 0 | 0 | 0 | 0 | 0 | 0 | 0 | 0 | 0 | 0 | 0 | 0 | 2.77 | 40.8 | 25.8 | 10.4 | 7.1 | 50.4 |
| *Lactiplantibacillus plantarum* | 0 | 0 | 0.45 | 0.46 | 0 | 0 | 0 | 2.16 | 0 | 2.27 | 2.93 | 2.41 | 0 | 2.81 | 2.6 | 3.18 | 0 | 0 | 1.06 | 0.74 | 0 | 1.67 | 1.39 | 2.32 | 0 | 0 | 0 | 0 | 0 | 0 | 0 | 0 | 0.81 | 21.9 | 0 | 0 | 0 | 2.42 |
| *Cronobacter malonaticus* | 19.7 | 0.73 | 0.27 | 0.35 | 11.6 | 0.73 | 0 | 0.77 | 0 | 0.33 | 0 | 0.36 | 0 | 0 | 0.3 | 0 | 21.8 | 0.95 | 0.83 | 0.6 | 0 | 0 | 0 | 0 | 7.12 | 8.81 | 12.3 | 9.94 | 0 | 0 | 0 | 0 | 0 | 0 | 0 | 0 | 0 | 0 |
| *Tetragenococcus halophilus* | 0 | 0 | 0 | 0 | 0 | 0 | 0 | 0 | 0 | 2.02 | 0 | 0 | 0 | 0 | 0 | 0 | 0 | 0 | 0 | 0 | 0 | 0 | 0 | 0 | 0 | 0 | 0 | 0 | 0 | 10.5 | 2.67 | 18.5 | 0 | 0 | 0 | 0 | 0 | 0 |
| *Bacillus cereus* | 0 | 0 | 0 | 0 | 0 | 0 | 0 | 0 | 17.1 | 0 | 0 | 0 | 7.22 | 0 | 0 | 0 | 0 | 0 | 0 | 0 | 0 | 0 | 0 | 0 | 0 | 0 | 0 | 0 | 0 | 0 | 0 | 0 | 0 | 0 | 0 | 0 | 0 | 0 |
| *Bacillus subtilis* | 0 | 0 | 0 | 0 | 0 | 0 | 0 | 0 | 0 | 0 | 0 | 0 | 0 | 0 | 0 | 0 | 0 | 0 | 0 | 0 | 0 | 0 | 0 | 0 | 0 | 0 | 0 | 0 | 0 | 0 | 0 | 0 | 0 | 0 | 1.18 | 14 | 1.11 | 0 |
| *Weissella hellenica* | 0 | 0 | 0 | 0 | 0 | 0 | 0 | 0 | 0 | 0 | 0 | 0 | 0 | 0 | 0 | 0 | 0 | 0 | 0 | 0 | 0 | 0 | 0 | 0 | 0 | 0 | 0 | 0 | 0 | 0 | 0 | 0 | 0 | 0 | 0 | 0 | 13.4 | 0 |
| *Enterococcus faecalis* | 0 | 0 | 0 | 0 | 0 | 0 | 0 | 0 | 0 | 0 | 0 | 0 | 0 | 0 | 0 | 0 | 0 | 0 | 0 | 0 | 0 | 0 | 0 | 0 | 0 | 0 | 0 | 0 | 0 | 0 | 0 | 0 | 0 | 0 | 0 | 0 | 0 | 8.04 |
| *Latilactobacillus sakei* | 0 | 0 | 0 | 0 | 0 | 0 | 0 | 0 | 0 | 0 | 0 | 0 | 0 | 0 | 0 | 0 | 0 | 0 | 0 | 0 | 0 | 0 | 0 | 0 | 0 | 0 | 0 | 0 | 0 | 0 | 0 | 0 | 4.64 | 0 | 0 | 0 | 0 | 0 |
| *Debaryomyces hansenii* | 0 | 0 | 0 | 0 | 0 | 0 | 0 | 0 | 0 | 0 | 0 | 0 | 0 | 0 | 0 | 0 | 0 | 0 | 0 | 0 | 0 | 0 | 0 | 0 | 0 | 0 | 0 | 0 | 0 | 0 | 0 | 0 | 0 | 0 | 0 | 4.16 | 0 | 0 |
| *Staphylococcus aureus* | 0 | 0 | 0 | 0 | 0 | 0 | 0.83 | 3.18 | 0 | 0 | 0 | 0 | 0 | 0 | 0 | 0 | 0 | 0 | 0 | 0 | 0 | 0 | 0 | 0 | 0 | 0 | 0 | 0 | 0 | 0 | 0 | 0 | 2.69 | 0 | 0 | 0 | 0 | 0 |
| *Staphylococcus simulans* | 0 | 0 | 0 | 0 | 0 | 0 | 0 | 0 | 0 | 0 | 0 | 0 | 0 | 0 | 0 | 0 | 0 | 0 | 0 | 0 | 0 | 0 | 2.55 | 0 | 0 | 0 | 0 | 0 | 0 | 0 | 0 | 0 | 0 | 0 | 0 | 0 | 0 | 0 |
| *Staphylococcus lugdunensis* | 0 | 0 | 0 | 0 | 0 | 0 | 0 | 0 | 0 | 0 | 0 | 0 | 0 | 0 | 0 | 0 | 0 | 0 | 0 | 0 | 0 | 0 | 0 | 0 | 0 | 0 | 0 | 0 | 0 | 0 | 0 | 0 | 0 | 0 | 0 | 1.46 | 0 | 0 |
| *Weissella confusa* | 0 | 0 | 0 | 0 | 0 | 0 | 0 | 0 | 0 | 0 | 0 | 0 | 0 | 0 | 0 | 0 | 0 | 0 | 0 | 0 | 0 | 0 | 0 | 0 | 0 | 0 | 0 | 0 | 0 | 0 | 0 | 0 | 0.26 | 0 | 0 | 0 | 1.21 | 0 |
| *Weissella viridescens* | 0 | 0 | 0 | 0 | 0 | 0 | 0 | 0 | 0 | 0 | 0 | 0 | 0 | 0 | 0 | 0 | 0 | 0 | 0 | 0 | 0 | 0 | 0 | 0 | 0 | 0 | 0 | 0 | 0 | 0 | 0 | 0 | 1.18 | 0 | 0 | 0 | 0 | 0 |
| *Weissella minor* | 0 | 0 | 0 | 0 | 0 | 0 | 0 | 0 | 0 | 0.39 | 0 | 0 | 0 | 0 | 0 | 0 | 0 | 0 | 0 | 0 | 0 | 0 | 0 | 0 | 0 | 0 | 0 | 0 | 0 | 0 | 0 | 0 | 1.03 | 0 | 0 | 0 | 0 | 0 |
| *[Candida] glabrata* | 0 | 0 | 0 | 0 | 0 | 0 | 0 | 0 | 0 | 0 | 0.61 | 0 | 0 | 0 | 0 | 0 | 0 | 0 | 0 | 0 | 0 | 0 | 0.81 | 0 | 0 | 0 | 0 | 0 | 0 | 0 | 0 | 0 | 0 | 0 | 0 | 0 | 0 | 0 |
| *Cyberlindnera fabianii* | 0 | 0 | 0 | 0 | 0 | 0 | 0 | 0 | 0 | 0 | 0 | 0 | 0 | 0 | 0 | 0 | 0 | 0 | 0 | 0 | 0 | 0 | 0 | 0 | 0 | 0 | 0 | 0 | 0 | 0 | 0 | 0 | 0.37 | 0 | 0 | 0 | 0 | 0 |
| *Staphylococcus taiwanensis* | 0 | 0 | 0 | 0 | 0 | 0 | 0 | 0 | 0 | 0 | 0 | 0 | 0 | 0 | 0 | 0 | 0 | 0 | 0 | 0 | 0 | 0 | 0 | 0 | 0 | 0 | 0 | 0 | 0 | 0 | 0 | 0 | 0 | 0 | 0.34 | 0 | 0 | 0 |
| *Millerozyma farinosa* | 0 | 0 | 0 | 0 | 0 | 0 | 0 | 0 | 0 | 0 | 0 | 0 | 0 | 0 | 0 | 0 | 0 | 0 | 0 | 0 | 0 | 0 | 0 | 0 | 0 | 0 | 0 | 0 | 0 | 0.37 | 0 | 0 | 0 | 0 | 0 | 0 | 0 | 0 |

**Table S5.** Quality of prokaryotic and their ANIs with the closest reference genome. The colour code indicates the MAGs' completeness, ranging from low (red) to high (green), and contamination, ranging from low (red) to high (blue).

| MAGid | Marker Lineage | # Genomes | # Markers | # Marker Sets | 0 | 1 | 2 | 3 | 4 | 5+ | Completenes | Contamina | Closely related species | ANI |
|---|---|---|---|---|---|---|---|---|---|---|---|---|---|---|
| F.1_bin.1 | g__Bacillus | 93 | 711 | 241 | 10 | 698 | 3 | 0 | 0 | 0 | 98.76 | 0.09 | Bacillus amyloliquefaciens DSM7 | 99.6 |
| F1.3_bin.3 | g__Staphylococcus | 60 | 773 | 178 | 31 | 720 | 22 | 0 | 0 | 0 | 97.49 | 2.32 | Staphylococcus pasteuri BAB3 | 98.8 |
| F1.3_bin.4 | o__Lactobacillales | 85 | 367 | 162 | 1 | 366 | 0 | 0 | 0 | 0 | 99.38 | 0 | Pediococcus pentosaceus CGMCC 7049 | 99.7 |
| F1.3_bin.5 | g__Bacillus | 93 | 711 | 241 | 37 | 669 | 5 | 0 | 0 | 0 | 94.19 | 1.09 | Bacillus amyloliquefaciens DSM7 | 99.6 |
| F2.3_bin.2 | o__Lactobacillales | 85 | 367 | 162 | 1 | 366 | 0 | 0 | 0 | 0 | 99.38 | 0 | Pediococcus pentosaceus CGMCC 7049 | 99.6 |
| F2.3_bin.4 | g__Bacillus | 93 | 711 | 241 | 6 | 697 | 8 | 0 | 0 | 0 | 98.65 | 0.23 | Bacillus amyloliquefaciens DSM 7 | 99.7 |
| F2.3_bin.5 | g__Staphylococcus | 60 | 773 | 178 | 4 | 759 | 10 | 0 | 0 | 0 | 99.11 | 0.64 | Staphylococcus pasteuri BAB3 | 98.9 |
| F3.3_bin.1 | g__Staphylococcus | 60 | 773 | 178 | 17 | 746 | 10 | 0 | 0 | 0 | 97.67 | 0.94 | Staphylococcus pasteuri BAB3 | 98.9 |
| F3.3_bin.3 | o__Lactobacillales | 85 | 367 | 162 | 56 | 311 | 0 | 0 | 0 | 0 | 95.06 | 0 | Pediococcus pentosaceus CGMCC 7049 | 99.6 |
| F3.3_bin.5 | g__Bacillus | 93 | 711 | 241 | 27 | 673 | 10 | 1 | 0 | 0 | 97.63 | 1.48 | Bacillus amyloliquefaciens DSM 7 | 99.6 |
| HB_bin.4 | o__Lactobacillales | 85 | 367 | 162 | 10 | 356 | 1 | 0 | 0 | 0 | 95.83 | 0.21 | Pediococcus pentosaceus SL4 | 99.5 |
| L.1_bin.1 | g__Bacillus | 93 | 711 | 241 | 10 | 699 | 2 | 0 | 0 | 0 | 98.76 | 0.06 | Bacillus amyloliquefaciens DSM 7 | 99.6 |
| L1.3_bin.2 | g__Staphylococcus | 60 | 773 | 178 | 3 | 766 | 4 | 0 | 0 | 0 | 99.17 | 0.46 | Staphylococcus pasteuri BAB3 | 98.9 |
| L1.3_bin.3 | o__Lactobacillales | 85 | 367 | 162 | 1 | 366 | 0 | 0 | 0 | 0 | 99.38 | 0 | Pediococcus pentosaceus CGMCC 7049 | 99.7 |
| L1.3_bin.4 | g__Bacillus | 93 | 711 | 241 | 2 | 707 | 2 | 0 | 0 | 0 | 99.58 | 0.06 | Bacillus amyloliquefaciens DSM 7 | 99.7 |
| L2.3_bin.2 | o__Lactobacillales | 85 | 367 | 162 | 1 | 366 | 0 | 0 | 0 | 0 | 99.38 | 0 | Pediococcus pentosaceus CGMCC 7049 | 99.6 |
| L2.3_bin.3 | g__Bacillus | 93 | 711 | 241 | 1 | 708 | 2 | 0 | 0 | 0 | 99.59 | 0.06 | Bacillus amyloliquefaciens DSM 7 | 99.7 |
| L3.3_bin.1 | o__Lactobacillales | 85 | 367 | 162 | 5 | 362 | 0 | 0 | 0 | 0 | 98.15 | 0 | Pediococcus pentosaceus CGMCC 7049 | 99.6 |
| L3.3_bin.2 | o__Lactobacillales | 85 | 367 | 162 | 2 | 360 | 5 | 0 | 0 | 0 | 99.07 | 2.78 | Lactiplantibacillus plantarum subsp. plantarum ST-III | 99.1 |
| L3.3_bin.5 | g__Bacillus | 93 | 711 | 241 | 1 | 707 | 3 | 0 | 0 | 0 | 99.59 | 0.47 | Bacillus amyloliquefaciens DSM 7 | 99.6 |
| MS_bin.1 | g__Bacillus | 93 | 711 | 241 | 1 | 708 | 2 | 0 | 0 | 0 | 99.59 | 0.04 | Bacillus amyloliquefaciens DSM 7 | 97.7 |
| MS_bin.4 | o__Lactobacillales | 85 | 367 | 162 | 5 | 357 | 5 | 0 | 0 | 0 | 98.46 | 2.78 | Lactiplantibacillus plantarum B21 | 99.4 |
| MS_bin.5 | g__Staphylococcus | 60 | 773 | 178 | 42 | 717 | 14 | 0 | 0 | 0 | 98.85 | 1 | Staphylococcus epidermidis ATCC 12228 | 97.1 |
| NF.1_bin.1 | g__Bacillus | 93 | 711 | 241 | 1 | 707 | 3 | 0 | 0 | 0 | 99.59 | 0.09 | Bacillus amyloliquefaciens DSM 7 | 99.6 |
| NF1.3_bin.2 | o__Lactobacillales | 85 | 367 | 162 | 3 | 361 | 3 | 0 | 0 | 0 | 98.46 | 1.54 | Lactobacillus plantarum subsp. plantarum ST-III | 99.2 |
| NF1.3_bin.4 | g__Bacillus | 93 | 711 | 241 | 1 | 707 | 3 | 0 | 0 | 0 | 99.59 | 0.09 | Bacillus amyloliquefaciens DSM 7 | 99.7 |
| NF1.3_bin.6 | o__Lactobacillales | 85 | 367 | 162 | 29 | 337 | 1 | 0 | 0 | 0 | 93.83 | 0.62 | Pediococcus pentosaceus CGMCC 7049 | 99.6 |
| NF2.3_bin.1 | o__Lactobacillales | 85 | 367 | 162 | 1 | 353 | 13 | 0 | 0 | 0 | 99.38 | 3.09 | Pediococcus pentosaceus CGMCC 7049 | 99.6 |
| NF2.3_bin.4 | o__Lactobacillales | 85 | 367 | 162 | 5 | 359 | 3 | 0 | 0 | 0 | 97.53 | 1.85 | Lactobacillus plantarum subsp. plantarum ST-III | 99.2 |
| NF2.3_bin.5 | g__Bacillus | 93 | 711 | 241 | 1 | 708 | 2 | 0 | 0 | 0 | 99.59 | 0.06 | Bacillus amyloliquefaciens DSM 7 | 99.7 |
| NF3.3_bin.1 | g__Bacillus | 93 | 711 | 241 | 1 | 708 | 2 | 0 | 0 | 0 | 99.59 | 0.06 | Bacillus amyloliquefaciens DSM 7 | 99.7 |
| NF3.3_bin.4 | o__Lactobacillales | 85 | 367 | 162 | 3 | 359 | 5 | 0 | 0 | 0 | 98.46 | 2.78 | Lactobacillus plantarum subsp. plantarum ST-III | 99.2 |
| NL.1_bin.1 | g__Bacillus | 93 | 711 | 241 | 1 | 707 | 3 | 0 | 0 | 0 | 99.59 | 0.09 | Bacillus amyloliquefaciens DSM 7 | 99.6 |
| NL1.3_bin.2 | g__Bacillus | 93 | 711 | 241 | 1 | 705 | 5 | 0 | 0 | 0 | 99.59 | 0.25 | Bacillus amyloliquefaciens DSM 7 | 99.7 |
| NL1.3_bin.4 | o__Lactobacillales | 85 | 367 | 162 | 3 | 360 | 4 | 0 | 0 | 0 | 98.46 | 1.85 | Lactobacillus plantarum subsp. plantarum ST-III | 99.1 |
| NL2.3_bin.1 | g__Bacillus | 93 | 711 | 241 | 1 | 708 | 2 | 0 | 0 | 0 | 99.59 | 0.06 | Bacillus amyloliquefaciens DSM 7 | 99.7 |
| NL2.3_bin.2 | g__Staphylococcus | 60 | 773 | 178 | 36 | 722 | 15 | 0 | 0 | 0 | 97.32 | 1.94 | Staphylococcus pasteuri BAB3 | 98.7 |
| NL2.3_bin.3 | o__Lactobacillales | 85 | 367 | 162 | 4 | 359 | 4 | 0 | 0 | 0 | 97.84 | 2.16 | Lactobacillus plantarum subsp. plantarum ST-III | 99.3 |
| NL2.3_bin.4 | o__Lactobacillales | 85 | 367 | 162 | 1 | 366 | 0 | 0 | 0 | 0 | 99.38 | 0 | Pediococcus pentosaceus CGMCC 7049 | 99.6 |
| NL3.3_bin.2 | o__Lactobacillales | 85 | 367 | 162 | 2 | 364 | 1 | 0 | 0 | 0 | 98.77 | 0.62 | Pediococcus pentosaceus CGMCC 7049 | 99.6 |
| NL3.3_bin.3 | o__Lactobacillales | 85 | 367 | 162 | 4 | 359 | 4 | 0 | 0 | 0 | 98.15 | 2.16 | Lactobacillus plantarum subsp. plantarum ST-III | 99.2 |
| NL3.3_bin.4 | g__Bacillus | 93 | 711 | 241 | 1 | 708 | 2 | 0 | 0 | 0 | 99.59 | 0.06 | Bacillus amyloliquefaciens DSM 7 | 99.7 |
| NP.1_bin.1 | g__Bacillus | 93 | 711 | 241 | 1 | 708 | 2 | 0 | 0 | 0 | 99.59 | 0.06 | Bacillus amyloliquefaciens DSM 7 | 99.6 |
| NP1.3_bin.3 | g__Staphylococcus | 60 | 773 | 178 | 8 | 749 | 16 | 0 | 0 | 0 | 98.28 | 0.87 | Staphylococcus pasteuri BAB3 | 98.9 |
| NP1.3_bin.4 | o__Lactobacillales | 85 | 367 | 162 | 3 | 360 | 4 | 0 | 0 | 0 | 98.46 | 2.47 | Lactobacillus plantarum subsp. plantarum ST-III | 99.2 |
| NP1.3_bin.5 | g__Bacillus | 93 | 711 | 241 | 1 | 705 | 5 | 0 | 0 | 0 | 99.59 | 0.51 | Bacillus amyloliquefaciens DSM 7 | 99.7 |
| NP1.3_bin.6 | o__Lactobacillales | 85 | 367 | 162 | 1 | 366 | 0 | 0 | 0 | 0 | 99.38 | 0 | Pediococcus pentosaceus CGMCC 7049 | 99.6 |
| NP2.3_bin.2 | g__Staphylococcus | 60 | 773 | 178 | 23 | 707 | 42 | 1 | 0 | 0 | 98.37 | 4.85 | Staphylococcus pasteuri BAB3 | 98.9 |
| NP2.3_bin.4 | g__Bacillus | 93 | 711 | 241 | 1 | 704 | 6 | 0 | 0 | 0 | 99.59 | 0.21 | Bacillus amyloliquefaciens DSM 7 | 99.6 |
| NP2.3_bin.5 | o__Lactobacillales | 85 | 367 | 162 | 4 | 356 | 7 | 0 | 0 | 0 | 98.15 | 3.29 | Lactobacillus plantarum subsp. plantarum ST-III | 99.3 |
| NP2.3_bin.7 | c__Bacilli | 586 | 325 | 181 | 12 | 310 | 3 | 0 | 0 | 0 | 97.24 | 1.24 | Staphylococcus simulans CJ16 | 99.4 |
| NP3.3_bin.2 | o__Lactobacillales | 85 | 367 | 162 | 1 | 366 | 0 | 0 | 0 | 0 | 99.38 | 0 | Pediococcus pentosaceus CGMCC 7049 | 99.6 |
| NP3.3_bin.3 | g__Staphylococcus | 60 | 773 | 178 | 10 | 740 | 22 | 1 | 0 | 0 | 98.96 | 1.87 | Staphylococcus pasteuri BAB3 | 98.8 |
| NP3.3_bin.4 | g__Bacillus | 93 | 711 | 241 | 10 | 697 | 4 | 0 | 0 | 0 | 99.03 | 0.23 | Bacillus amyloliquefaciens DSM 7 | 99.7 |
| NP3.3_bin.5 | o__Lactobacillales | 85 | 367 | 162 | 5 | 360 | 2 | 0 | 0 | 0 | 97.53 | 0.93 | Lactobacillus plantarum subsp. plantarum ST-III | 99.2 |
| P.1_bin.1 | f__Enterobacteriaceae | 223 | 876 | 305 | 55 | 813 | 8 | 0 | 0 | 0 | 98.64 | 0.7 | Cronobacter malonaticus 685 | 98.6 |
| P.1_bin.2 | g__Bacillus | 93 | 711 | 241 | 4 | 705 | 2 | 0 | 0 | 0 | 99.28 | 0.06 | Bacillus amyloliquefaciens DSM 7 | 99.7 |
| P1.3_bin.1 | o__Lactobacillales | 85 | 367 | 162 | 1 | 366 | 0 | 0 | 0 | 0 | 99.38 | 0 | Pediococcus pentosaceus CGMCC 7049 | 99.7 |
| P1.3_bin.3 | g__Bacillus | 93 | 711 | 241 | 1 | 707 | 3 | 0 | 0 | 0 | 99.59 | 0.47 | Bacillus amyloliquefaciens DSM 7 | 99.7 |
| P2.3_bin.1 | g__Staphylococcus | 60 | 773 | 178 | 41 | 702 | 30 | 0 | 0 | 0 | 95.09 | 3.45 | Staphylococcus epidermidis BCM-HMP0060 | 98.9 |
| P2.3_bin.3 | o__Lactobacillales | 85 | 367 | 162 | 17 | 339 | 11 | 0 | 0 | 0 | 93.67 | 3.72 | Pediococcus pentosaceus CGMCC 7049 | 99.6 |
| P2.3_bin.5 | o__Lactobacillales | 85 | 367 | 162 | 8 | 352 | 7 | 0 | 0 | 0 | 97.66 | 3.14 | Lactobacillus plantarum subsp. plantarum ST-III | 99.1 |
| P2.3_bin.6 | g__Bacillus | 93 | 711 | 241 | 1 | 706 | 4 | 0 | 0 | 0 | 99.59 | 0.11 | Bacillus amyloliquefaciens DSM 7 | 99.7 |
| P3.3_bin.1 | g__Staphylococcus | 60 | 773 | 178 | 51 | 703 | 19 | 0 | 0 | 0 | 90.59 | 1.45 | Staphylococcus pasteuri BAB3 | 98.8 |
| P3.3_bin.2 | g__Bacillus | 93 | 711 | 241 | 1 | 708 | 2 | 0 | 0 | 0 | 99.59 | 0.06 | Bacillus amyloliquefaciens DSM 7 | 99.7 |
| P3.3_bin.3 | o__Lactobacillales | 85 | 367 | 162 | 17 | 339 | 11 | 0 | 0 | 0 | 93.67 | 3.72 | Lactobacillus plantarum subsp. plantarum ST-III | 98.9 |
| P3.3_bin.4 | g__Staphylococcus | 60 | 773 | 178 | 60 | 681 | 32 | 0 | 0 | 0 | 92.13 | 3.76 | Staphylococcus epidermidis | 98.9 |
| P3.3_bin.5 | o__Lactobacillales | 85 | 367 | 162 | 17 | 350 | 0 | 0 | 0 | 0 | 95.06 | 0 | Pediococcus pentosaceus CGMCC 7049 | 99.1 |
| PB_bin.2 | g__Staphylococcus | 60 | 773 | 178 | 2 | 766 | 5 | 0 | 0 | 0 | 99.73 | 1.31 | Staphylococcus warneri | 99.7 |
| PB_bin.3 | o__Lactobacillales | 85 | 367 | 162 | 1 | 366 | 0 | 0 | 0 | 0 | 99.38 | 0 | Pediococcus acidilactici | 98.7 |
| PB_bin.4 | g__Bacillus | 93 | 711 | 241 | 74 | 626 | 11 | 0 | 0 | 0 | 94.25 | 1.77 | Bacillus subtilis SR1 | 98.5 |
| PR_bin.1 | g__Staphylococcus | 60 | 773 | 178 | 74 | 694 | 5 | 0 | 0 | 0 | 93.72 | 0.33 | Staphylococcus epidermidis ATCC 12228 | 99.2 |
| PR_bin.2 | g__Bacillus | 93 | 711 | 241 | 8 | 703 | 0 | 0 | 0 | 0 | 98.76 | 0 | Bacillus subtilis NCIB 3610 | 98.6 |
| PR_bin.3 | o__Lactobacillales | 85 | 367 | 162 | 1 | 366 | 0 | 0 | 0 | 0 | 99.38 | 0 | Pediococcus acidilactici NGRI 0510Q | 98.7 |
| R.1_bin.1 | g__Bacillus | 93 | 711 | 241 | 1 | 708 | 2 | 0 | 0 | 0 | 99.59 | 0.06 | Bacillus amyloliquefaciens DSM 7 | 99.6 |
| R1.3_bin.2 | g__Bacillus | 93 | 711 | 241 | 1 | 707 | 3 | 0 | 0 | 0 | 99.59 | 0.09 | Bacillus amyloliquefaciens DSM 7 | 99.6 |
| R2.3_bin.2 | g__Bacillus | 93 | 711 | 241 | 42 | 668 | 1 | 0 | 0 | 0 | 95.02 | 0.04 | Bacillus amyloliquefaciens DSM 7 | 99.5 |
| R3.3_bin.1 | g__Bacillus | 93 | 711 | 241 | 1 | 708 | 2 | 0 | 0 | 0 | 99.59 | 0.06 | Bacillus amyloliquefaciens DSM 7 | 99.7 |
| S.1_bin.1 | g__Bacillus | 93 | 711 | 241 | 87 | 612 | 12 | 0 | 0 | 0 | 91.83 | 1.38 | Bacillus amyloliquefaciens DSM 7 | 99.6 |
| S1.3_bin.3 | c__Bacilli | 725 | 279 | 151 | 1 | 276 | 2 | 0 | 0 | 0 | 99.34 | 0.26 | Tetragenococcus halophilus NBRC 12172 | 95.2 |
| S2.3_bin.1 | c__Bacilli | 725 | 279 | 151 | 1 | 278 | 0 | 0 | 0 | 0 | 99.34 | 0 | Tetragenococcus halophilus NBRC 12172 | 95.2 |
| S2.3_bin.3 | g__Staphylococcus | 60 | 773 | 178 | 51 | 700 | 21 | 1 | 0 | 0 | 95.92 | 3.67 | Staphylococcus pasteuri | 98.5 |
| S3.3_bin.2 | c__Bacilli | 725 | 279 | 151 | 1 | 278 | 0 | 0 | 0 | 0 | 99.34 | 0 | Tetragenococcus halophilus NBRC 12172 | 95.2 |
| SY_bin.1 | g__Staphylococcus | 60 | 773 | 178 | 109 | 656 | 8 | 0 | 0 | 0 | 91.67 | 0.8 | Staphylococcus warneri | 99.6 |
| SY_bin.3 | o__Lactobacillales | 293 | 475 | 267 | 2 | 471 | 2 | 0 | 0 | 0 | 99.25 | 0.5 | Enterococcus faecium T110 | 97.8 |
| SY_bin.4 | o__Lactobacillales | 177 | 350 | 163 | 2 | 348 | 0 | 0 | 0 | 0 | 98.77 | 0 | Weissella hellenica | 98.8 |
| TS_bin.1 | o__Lactobacillales | 85 | 367 | 162 | 61 | 303 | 3 | 0 | 0 | 0 | 93.95 | 1.23 | Pediococcus pentosaceus SL4 | 99 |
| TS_bin.3 | o__Lactobacillales | 85 | 367 | 162 | 16 | 347 | 4 | 0 | 0 | 0 | 96.6 | 2.47 | Lactobacillus plantarum subsp. plantarum ST-III | 98.9 |
| TS_bin.5 | o__Lactobacillales | 85 | 367 | 162 | 1 | 366 | 0 | 0 | 0 | 0 | 99.38 | 0 | Pediococcus acidilactici | 98.8 |
| TS_bin.6 | g__Staphylococcus | 60 | 773 | 178 | 2 | 757 | 14 | 0 | 0 | 0 | 99.73 | 1.4 | Staphylococcus warneri | 99.7 |

**Table S5.** Quality of A. oryzae MAGs. The colour code indicates the MAGs' completeness, ranging from low (red) to high (green), and contamination, ranging from low (red) to high (blue).

| | Complete | | Complete | Fragmented | Missing |
|---|---|---|---|---|---|
| | Single-copy | Duplicated | | | |
| **F.1** | 83.90% | 0.40% | 84.30% | 10.20% | 5.50% |
| **L.1** | 91.80% | 0.40% | 92.20% | 5.10% | 2.70% |
| **NL.1** | 75.30% | 0.80% | 76.10% | 14.90% | 9.00% |
| **P.1** | 80.80% | 0.40% | 81.20% | 11.80% | 7.00% |
| **PR** | 93.30% | 0.40% | 93.70% | 3.90% | 2.40% |
| **R.1** | 94.50% | 0.40% | 94.90% | 2.70% | 2.40% |
| **R1.3** | 91.40% | 0.40% | 91.80% | 4.30% | 3.90% |
| **R2.3** | 91.40% | 0.00% | 91.40% | 4.70% | 3.90% |
| **R3.3** | 92.50% | 0.40% | 92.90% | 4.30% | 2.80% |
| **S.1** | 92.20% | 0.40% | 92.60% | 5.10% | 2.30% |

**Table S6.** ANI analyses for the recovered MAGs of S. epidermidis, P. acidilacti, L. plantarum, P. pentosaceus and S. warneri. The colour code indicates the ANI values ranging from low (red) to high (green).

**S. epidermidis**

| | MS_bin.5. | PR_bin.1.f | P2.3_bin.1 | P3.3_bin.4.fa |
|---|---|---|---|---|
| MS_bin.5.fa | * | 96.85 | 97.05 | 97.14 |
| PR_bin.1.fa | 96.48 | * | 98.73 | 98.58 |
| P2.3_bin.1.fa | 97.06 | 98.94 | * | 99.1 |
| P3.3_bin.4.fa | 97.26 | 99.08 | 99.37 | * |

**P. acidilacti**

| | PR_bin.3.f | PB_bin.3.f | TS_bin.5.fa |
|---|---|---|---|
| PR_bin.3.f | * | 99.99 | 99.58 |
| PB_bin.3.f | 99.95 | * | 99.53 |
| TS_bin.5.fa | 99.57 | 99.55 | * |

**S. warneri**

| | TS_bin.6.fa | SY_bin.1.fa | PB_bin.2.fa |
|---|---|---|---|
| TS_bin.6.fa | * | 99.55 | 99.7 |
| SY_bin.1.fa | 99.54 | * | 99.67 |
| PB_bin.2.f | 99.31 | 99.35 | * |

**L. plantarum**

| | MS_bin.4. | TS_bin.3.fa | P3.3_bin.3 | L3.3_bin.2 | P2.3_bin.5 | NL1.3_bin | NP1.3_bin | NF3.3_bin | NL3.3_bin | NF1.3_bin | NP2.3_bin | NP3.3_bin | NF2.3_bin | NL2.3_bin.3.fa |
|---|---|---|---|---|---|---|---|---|---|---|---|---|---|---|
| MS_bin.4.fa | * | 99.47 | 98.83 | 98.84 | 98.8 | 98.96 | 98.88 | 98.86 | 99.01 | 98.89 | 98.86 | 98.85 | 98.82 | 99.01 |
| TS_bin.3.fa | 99.93 | * | 98.87 | 98.89 | 98.85 | 99.05 | 98.84 | 98.92 | 99.06 | 98.93 | 98.94 | 98.95 | 98.92 | 99.14 |
| P3.3_bin.3.fa | 98.74 | 98.32 | * | 99.7 | 99.76 | 99.48 | 98.88 | 99.08 | 99.33 | 98.91 | 98.97 | 99.13 | 99.04 | 99.33 |
| L3.3_bin.2.fa | 98.98 | 98.56 | 99.91 | * | 99.95 | 99.64 | 99.11 | 99.28 | 99.51 | 99.16 | 99.2 | 99.35 | 99.25 | 99.51 |
| P2.3_bin.5.fa | 98.87 | 98.52 | 99.86 | 99.87 | * | 99.57 | 99.05 | 99.22 | 99.48 | 99.15 | 99.15 | 99.35 | 99.2 | 99.46 |
| NL1.3_bin.4.fa | 98.77 | 98.37 | 99.29 | 99.28 | 99.24 | * | 99.32 | 99.36 | 99.47 | 99.29 | 99.26 | 99.34 | 99.23 | 99.31 |
| NP1.3_bin.4.fa | 98.71 | 98.23 | 98.9 | 98.95 | 98.94 | 99.55 | * | 99.73 | 99.7 | 99.67 | 99.64 | 99.49 | 99.46 | 99.48 |
| NF3.3_bin.4.fa | 98.82 | 98.31 | 99.08 | 99.16 | 99.16 | 99.68 | 99.78 | * | 99.77 | 99.69 | 99.66 | 99.56 | 99.57 | 99.55 |
| NL3.3_bin.3.fa | 98.88 | 98.48 | 99.21 | 99.29 | 99.29 | 99.72 | 99.6 | 99.71 | * | 99.54 | 99.54 | 99.42 | 99.4 | 99.49 |
| NF1.3_bin.2.fa | 98.95 | 98.46 | 99.12 | 99.19 | 99.21 | 99.76 | 99.89 | 99.88 | 99.83 | * | 99.8 | 99.67 | 99.73 | 99.67 |
| NP2.3_bin.5.fa | 99 | 98.5 | 99.13 | 99.18 | 99.18 | 99.68 | 99.84 | 99.81 | 99.78 | 99.8 | * | 99.61 | 99.61 | 99.6 |
| NP3.3_bin.5.fa | 99.11 | 98.63 | 99.48 | 99.54 | 99.55 | 99.76 | 99.74 | 99.74 | 99.75 | 99.72 | 99.7 | * | 99.74 | 99.72 |
| NF2.3_bin.4.fa | 99.02 | 98.51 | 99.42 | 99.46 | 99.47 | 99.75 | 99.84 | 99.86 | 99.84 | 99.82 | 99.82 | 99.76 | * | 99.76 |
| NL2.3_bin.3.fa | 99.15 | 98.7 | 99.64 | 99.63 | 99.66 | 99.73 | 99.71 | 99.73 | 99.75 | 99.7 | 99.71 | 99.69 | 99.68 | * |

**P. pentosaceus**

| | L2.3_bin.2 | L1.3_bin.3 | F2.3_bin.2 | F1.3_bin.4 | L3.3_bin.1 | P1.3_bin.1 | P2.3_bin.3 | F3.3_bin.3 | NL3.3_bin | NL2.3_bin | NP3.3_bin | NP1.3_bin | NF1.3_bin | NF2.3_bin | P3.3_bin.5 | TS_bin.1.fa | HB_bin.4.fa |
|---|---|---|---|---|---|---|---|---|---|---|---|---|---|---|---|---|---|
| L2.3_bin.2.fa | * | 99.99 | 100 | 99.94 | 99.91 | 99.92 | 99.93 | 99.83 | 99.74 | 99.74 | 99.74 | 99.74 | 99.67 | 99.74 | 99.33 | 98.92 | 98.73 |
| L1.3_bin.3.fa | 100 | * | 100 | 99.96 | 99.88 | 99.9 | 99.99 | 99.85 | 99.82 | 99.83 | 99.83 | 99.82 | 99.81 | 99.82 | 99.37 | 98.98 | 98.69 |
| F2.3_bin.2.fa | 100 | 100 | * | 99.94 | 99.86 | 99.88 | 99.9 | 99.79 | 99.71 | 99.73 | 99.71 | 99.69 | 99.66 | 99.71 | 99.34 | 98.99 | 98.75 |
| F1.3_bin.4.fa | 99.99 | 99.99 | 99.99 | * | 99.89 | 99.96 | 99.97 | 99.88 | 99.85 | 99.86 | 99.85 | 99.85 | 99.79 | 99.85 | 99.48 | 99.02 | 98.81 |
| L3.3_bin.1.fa | 99.86 | 99.86 | 99.86 | 99.79 | * | 99.86 | 99.78 | 99.69 | 99.81 | 99.81 | 99.81 | 99.75 | 99.61 | 99.81 | 99.3 | 98.87 | 98.58 |
| P1.3_bin.1.fa | 99.95 | 99.94 | 99.94 | 99.94 | 99.94 | * | 99.95 | 99.87 | 99.96 | 99.96 | 99.96 | 99.96 | 99.85 | 99.96 | 99.47 | 99.09 | 98.82 |
| P2.3_bin.3.fa | 99.92 | 99.92 | 99.9 | 99.88 | 99.84 | 99.8 | * | 99.76 | 99.83 | 99.84 | 99.83 | 99.83 | 99.82 | 99.82 | 99.23 | 99.01 | 98.69 |
| F3.3_bin.3.fa | 100 | 100 | 100 | 99.98 | 99.91 | 99.93 | 99.95 | * | 99.88 | 99.88 | 99.88 | 99.88 | 99.81 | 99.88 | 99.43 | 99.01 | 98.67 |
| NL3.3_bin.2.fa | 99.6 | 99.59 | 99.58 | 99.62 | 99.74 | 99.74 | 99.61 | 99.52 | * | 99.98 | 99.9 | 99.84 | 99.71 | 99.9 | 99 | 98.82 | 98.7 |
| NL2.3_bin.4.fa | 99.48 | 99.48 | 99.48 | 99.56 | 99.76 | 99.69 | 99.55 | 99.48 | 99.98 | * | 99.94 | 99.78 | 99.65 | 99.93 | 98.92 | 98.78 | 98.58 |
| NP3.3_bin.2.fa | 99.71 | 99.71 | 99.7 | 99.75 | 99.85 | 99.83 | 99.75 | 99.66 | 99.99 | 100 | * | 99.94 | 99.81 | 99.99 | 99.12 | 98.94 | 98.78 |
| NP1.3_bin.6.fa | 99.77 | 99.77 | 99.75 | 99.82 | 99.81 | 99.86 | 99.82 | 99.69 | 100 | 99.98 | 100 | * | 99.85 | 100 | 99.22 | 99.01 | 98.81 |
| NF1.3_bin.6.fa | 99.58 | 99.58 | 99.59 | 99.66 | 99.62 | 99.67 | 99.7 | 99.5 | 99.79 | 99.8 | 99.8 | 99.79 | * | 99.79 | 98.95 | 98.9 | 98.67 |
| NF2.3_bin.1.fa | 99.72 | 99.72 | 99.72 | 99.78 | 99.85 | 99.86 | 99.76 | 99.69 | 99.99 | 99.99 | 99.99 | 99.94 | 99.84 | * | 98.87 | 98.96 | 98.77 |
| P3.3_bin.5.fa | 99.93 | 99.93 | 99.93 | 99.93 | 99.94 | 99.96 | 99.9 | 99.82 | 99.87 | 99.87 | 99.87 | 99.87 | 99.7 | 99.87 | * | 99.11 | 98.7 |
| TS_bin.1.fa | 99.05 | 99.04 | 99.05 | 99.09 | 99.11 | 99.1 | 99.1 | 98.97 | 99.13 | 99.13 | 99.13 | 99.13 | 99.09 | 99.11 | 98.68 | * | 99.14 |
| HB_bin.4.fa | 98.98 | 98.98 | 98.98 | 98.98 | 99 | 99.01 | 98.98 | 98.81 | 99.06 | 99.06 | 99.06 | 99.06 | 99.02 | 99.06 | 98.39 | 99.4 | * |

**Table S7**. Unique genes annotated with BlastKoala KEGG from the recovered MAGs of S. epidermidis from the yellow pea group and maitake-soy strain.

**Pea group strains - 63 unique genes - 25.4% annotated - 16 entries**

| **Query** | | | **KO** | **Definition** | **Score** |
|---|---|---|---|---|---|
| unique/593/1/Org1_Gene4111 | Carbohydrate metabolis | Glycolysis / Gluconeogenesis an | K00001 | E1.1.1.1, adh; alcohol dehydrogenase [EC:1.1.1.1] | 251 |
| unique/1831/1/Org1_Gene1538 | Carbohydrate metabolis | Pentose phosphate pathway/ G | K00036 | G6PD, zwf; glucose-6-phosphate 1-dehydrogenase [EC:1.1.1.49 1.1.1.363] | 10 |
| unique/725/1/Org1_Gene2432 | Genetic Information Pro | Aminoacyl-tRNA biosynthesis | K01867 | WARS, trpS; tryptophanyl-tRNA synthetase [EC:6.1.1.2] | 279 |
| unique/1013/1/Org1_Gene2684 | Genetic Information Processing | | K03570 | mreC; rod shape-determining protein MreC | 239 |
| unique/1572/1/Org1_Gene3132 | Genetic Information Processing | | K03571 | mreD; rod shape-determining protein MreD | 163 |
| unique/2028/1/Org1_Gene5263 | Genetic Information Pro | Replication and repair | K07498 | K07498; putative transposase | 15 |
| unique/242/1/Org1_Gene218 | Glycan biosynthesis and | Teichoic acid biosynthesis | K00712 | tagE; poly(glycerol-phosphate) alpha-glucosyltransferase [EC:2.4.1.52] | 221 |
| unique/482/1/Org1_Gene2218 | Glycan biosynthesis and | Exopolysaccharide biosynthesis | K11936 | pgaC, icaA; poly-beta-1,6-N-acetyl-D-glucosamine synthase [EC:2.4.1.-] | 337 |
| unique/656/1/Org1_Gene2371 | Glycan biosynthesis and | Glycan metabolism | K21462 | icaC; probable poly-beta-1,6-N-acetyl-D-glucosamine export protein | 381 |
| unique/967/1/Org1_Gene855 | Glycan biosynthesis and | biofilm adhesin polysaccharide | K21478 | icaB; poly-beta-1,6-N-acetyl-D-glucosamine N-deacetylase [EC:3.5.1.-] | 180 |
| unique/1502/1/Org1_Gene4873 | Glycan biosynthesis and metabolism | | K21453 | icaR; TetR/AcrR family transcriptional regulator, biofilm operon repressor | 131 |
| unique/1950/1/Org1_Gene1623 | Glycan biosynthesis and | Exopolysaccharide biosynthesis | K21461 | icaD; poly-beta-1,6-N-acetyl-D-glucosamine synthesis protein | 19 |
| unique/222/1/Org1_Gene200 | Environmental Informat | Protein kinases | K25212 | hptS; two-component system, sensor histidine kinase HptS [EC:2.7.13.3] | 337 |
| unique/787/1/Org1_Gene694 | Environmental Informat | Two-component system | K25211 | hptA; hexose phosphate transport regulatory protein HptA | 213 |
| unique/980/1/Org1_Gene866 | Signaling and cellular pro | Transporters | K01990 | ABC-2.A; ABC-2 type transport system ATP-binding protein | 81 |
| unique/1173/1/Org1_Gene1019 | Signaling and cellular pro | Transporters | K01992 | ABC-2.P; ABC-2 type transport system permease protein | 25 |

**Table S7**. Unique genes annotated with BlastKoala KEGG from the recovered MAGs of S. epidermidis from the yellow pea group and maitake-soy strain.

**MS strain - 445 unique genes - 40.9% annotated - 182 entries**

| Query | | | KO | Definition | Score |
|---|---|---|---|---|---|
| unique/471/2/Org2_Gene1883 (414) | Amino acid metabolism | Lysine biosynthesis | K01439 | dapE; succinyl-diaminopimelate desuccinylase [EC:3.5.1.18] | 296 |
| unique/513/2/Org2_Gene1047 (399) | Amino acid metabolism | Cysteine and methionine metabo | K00789 | metK, MAT; S-adenosylmethionine synthetase [EC:2.5.1.6] | 371 |
| unique/1108/2/Org2_Gene111 (261) | Amino acid metabolism | Alanine, aspartate and glutamate | K13566 | NIT2, yafV; omega-amidase [EC:3.5.1.3] | 230 |
| unique/1185/2/Org2_Gene1156 (247) | Amino acid metabolism | Arginine biosynthesis | K00930 | argB; acetylglutamate kinase [EC:2.7.2.8] | 164 |
| unique/1228/2/Org2_Gene1504 (238) | Amino acid metabolism | Phenylalanine, tyrosine and trypt | K03785 | aroD; 3-dehydroquinate dehydratase I [EC:4.2.1.10] | 159 |
| unique/1348/2/Org2_Gene73 (213) | Amino acid metabolism | Cysteine and methionine metabo | K00640 | cysE; serine O-acetyltransferase [EC:2.3.1.30] | 203 |
| unique/1391/2/Org2_Gene356 (206) | Amino acid metabolism | Phenylalanine, tyrosine and trypt | K01817 | trpF; phosphoribosylanthranilate isomerase [EC:5.3.1.24] | 165 |
| unique/230/2/Org2_Gene1884 (507) | Brite Hierarchies, Protein fami | protease, Peptidases and inhibito | K01401 | aur; aureolysin [EC:3.4.24.29] | 363 |
| unique/425/2/Org2_Gene1371 (428) | Brite Hierarchies, Protein fami | protease, Peptidases and inhibito | K11749 | rseP; regulator of sigma E protease [EC:3.4.24.-] | 372 |
| unique/633/2/Org2_Gene1012 (361) | Brite Hierarchies, Protein fami | Peptidases and inhibitors | K01261 | pepA; glutamyl aminopeptidase [EC:3.4.11.7] | 299 |
| unique/997/2/Org2_Gene267 (283) | Brite Hierarchies, Protein fami | Peptidases and inhibitors | K01318 | sspA; glutamyl endopeptidase [EC:3.4.21.19] | 24 |
| unique/1114/2/Org2_Gene1698 (260) | Brite Hierarchies, Protein fami | Peptidases and inhibitors | K07052 | K07052; CAAX protease family protein | 14 |
| unique/1171/2/Org2_Gene1700 (250) | Brite Hierarchies, Protein fami | Peptidases and inhibitors | K07052 | K07052; CAAX protease family protein | 62 |
| unique/1190/2/Org2_Gene109 (246) | Brite Hierarchies, Protein fami | Peptidases and inhibitors | K07052 | K07052; CAAX protease family protein | 143 |
| unique/1411/2/Org2_Gene1652 (203) | Brite Hierarchies, Protein fami | Peptidases and inhibitors, Peptid | K07284 | srtA; sortase A [EC:3.4.22.70] | 170 |
| unique/362/2/Org2_Gene578 (450) | Carbohydrate metabolism | Citrate cycle | K00382 | DLD, lpd, pdhD; dihydrolipoamide dehydrogenase [EC:1.8.1.4] | 333 |
| unique/433/2/Org2_Gene581 (425) | Carbohydrate metabolism | Pyruvate metabolism | K00627 | DLAT, aceF, pdhC; pyruvate dehydrogenase E2 component (dihydrolipoamid | 301 |
| unique/1120/2/Org2_Gene1940 (259) | Carbohydrate metabolism | Butanoate metabolism | K03366 | butA, budC; meso-butanediol dehydrogenase / (S,S)-butanediol dehydrogen | 219 |
| unique/1249/2/Org2_Gene1457 (234) | Carbohydrate metabolism | Butanoate metabolism | K01575 | alsD, budA, aldC; acetolactate decarboxylase [EC:4.1.1.5] | 274 |
| unique/1351/2/Org2_Gene280 (213) | Carbohydrate metabolism | Glyoxylate and dicarboxylate met | K01091 | gph; phosphoglycolate phosphatase [EC:3.1.3.18] | 45 |
| unique/1827/2/Org2_Gene1508 (126) | Carbohydrate metabolism | Glyoxylate and dicarboxylate met | K02437 | gcvH, GCSH; glycine cleavage system H protein | 117 |
| unique/2053/2/Org2_Gene239 (77) | Carbohydrate metabolism | Butanoate metabolism | K01653 | E2.2.1.6S, ilvH, ilvN; acetolactate synthase I/III small subunit [EC:2.2.1.6] | 27 |
| unique/2172/2/Org2_Gene266 (50) | Cellular Processes | Cellular community - prokaryotes | K20337 | psmB; phenol-soluble modulin beta | 11 |
| unique/2182/2/Org2_Gene259 (46) | Cellular Processes | Cellular community - prokaryotes | K07800 | agrD; AgrD protein | 27 |
| unique/2193/2/Org2_Gene949 (44) | Cellular Processes | Cellular community - prokaryotes | K20337 | psmB; phenol-soluble modulin beta | 19 |
| unique/2195/2/Org2_Gene951 (44) | Cellular Processes | Cellular community - prokaryotes | K20337 | psmB; phenol-soluble modulin beta | 16 |
| unique/2198/2/Org2_Gene952 (43) | Cellular Processes | Cellular community - prokaryotes | K20337 | psmB; phenol-soluble modulin beta | 13 |
| unique/24/2/Org2_Gene675 (983) | Energy metabolism | Enzymes | K00123 | fdoG, fdhF, fdwA; formate dehydrogenase major subunit [EC:1.17.1.9] | 879 |
| unique/204/2/Org2_Gene1048 (530) | Energy metabolism | Glycolysis / Gluconeogenesis, Citr | K01610 | E4.1.1.49, pckA; phosphoenolpyruvate carboxykinase (ATP) [EC:4.1.1.49] | 470 |
| unique/206/2/Org2_Gene1441 (526) | Energy metabolism | Glycolysis / Gluconeogenesis, Pyr | K01895 | ACSS1_2, acs; acetyl-CoA synthetase [EC:6.2.1.1] | 605 |
| unique/1103/2/Org2_Gene645 (262) | Energy metabolism | | K02379 | fdhD; FdhD protein | 249 |
| unique/1510/2/Org2_Gene1942 (184) | Energy metabolism | Oxidative phosphorylation | K02826 | qoxA; cytochrome aa3-600 menaquinol oxidase subunit II [EC:7.1.1.5] | 81 |
| unique/1542/2/Org2_Gene300 (179) | Energy metabolism | Oxidative phosphorylation | K02113 | ATPF1D, atpH; F-type H+-transporting ATPase subunit delta | 140 |
| unique/1583/2/Org2_Gene299 (171) | Energy metabolism | Oxidative phosphorylation | K02109 | ATPF0B, atpF; F-type H+-transporting ATPase subunit b | 148 |
| unique/1682/2/Org2_Gene1526 (154) | Energy metabolism | | K04488 | iscU, nifU; nitrogen fixation protein NifU and related proteins | 154 |
| unique/2025/2/Org2_Gene806 (82) | Energy metabolism | | K05337 | fer; ferredoxin | 77 |
| unique/2078/2/Org2_Gene298 (70) | Energy metabolism | Oxidative phosphorylation | K02110 | ATPF0C, atpE; F-type H+-transporting ATPase subunit c | 74 |
| unique/186/2/Org2_Gene510 (547) | Environmental Information Pr | Membrane transport | K15580 | oppA, mppA; oligopeptide transport system substrate-binding protein | 410 |
| unique/631/2/Org2_Gene1116 (363) | Environmental Information Pr | Two-component system | K20487 | nisK, spaK; two-component system, OmpR family, lantibiotic biosynthesis se | 112 |
| unique/703/2/Org2_Gene1511 (341) | Environmental Information Pr | Membrane transport | K02071 | metN; D-methionine transport system ATP-binding protein | 304 |
| unique/750/2/Org2_Gene521 (331) | Environmental Information Pr | Membrane transport | K09815 | znuA; zinc transport system substrate-binding protein | 198 |
| unique/776/2/Org2_Gene2108 (325) | Environmental Information Pr | Membrane transport, ABC transp | K02040 | pstS; phosphate transport system substrate-binding protein | 250 |
| unique/872/2/Org2_Gene2107 (308) | Environmental Information Pr | Membrane transport, ABC transp | K02037 | pstC; phosphate transport system permease protein | 261 |
| unique/907/2/Org2_Gene2106 (301) | Environmental Information Pr | Membrane transport, ABC transp | K02038 | pstA; phosphate transport system permease protein | 231 |
| unique/911/2/Org2_Gene1120 (301) | Environmental Information Pr | Membrane transport, ABC transp | K01990 | ABC-2.A; ABC-2 type transport system ATP-binding protein | 136 |
| unique/957/2/Org2_Gene2105 (291) | Environmental Information Pr | Membrane transport, ABC transp | K02036 | pstB; phosphate transport system ATP-binding protein [EC:7.3.2.1] | 268 |
| unique/1059/2/Org2_Gene1513 (270) | Environmental Information Pr | Membrane transport | K02073 | metQ; D-methionine transport system substrate-binding protein | 223 |
| unique/1106/2/Org2_Gene644 (261) | Environmental Information Pr | ABC transporters | K02020 | modA; molybdate transport system substrate-binding protein | 186 |
| unique/1261/2/Org2_Gene1512 (231) | Environmental Information Pr | Membrane transport, ABC transp | K02072 | metI; D-methionine transport system permease protein | 187 |
| unique/1315/2/Org2_Gene1117 (221) | Environmental Information Pr | Signal transduction, Two-compon | K20488 | nisR, spaR; two-component system, OmpR family, lantibiotic biosynthesis res | 151 |
| unique/1331/2/Org2_Gene2133 (219) | Environmental Information Pr | Signal transduction, Two-compon | K18941 | arlR; two-component system, OmpR family, response regulator ArlR | 252 |
| unique/1435/2/Org2_Gene871 (199) | Environmental Information Pr | Signal transduction | K04564 | SOD2; superoxide dismutase, Fe-Mn family [EC:1.15.1.1] | 275 |
| unique/4/2/Org2_Gene1901 (1481) | Genetic Information Processing | | K03466 | ftsK, spoIIIE; DNA segregation ATPase FtsK/SpoIIIE, S-DNA-T family | 1037 |
| unique/5/2/Org2_Gene1373 (1438) | Genetic Information Processin | DNA replication | K03763 | polC; DNA polymerase III subunit alpha, Gram-positive type [EC:2.7.7.7] | 1380 |
| unique/36/2/Org2_Gene1999 (893) | Genetic Information Processing | | K02469 | gyrA; DNA gyrase subunit A [EC:5.6.2.2] | 802 |
| unique/163/2/Org2_Gene1372 (567) | Genetic Information Processin | Aminoacyl-tRNA biosynthesis | K01881 | PARS, proS; prolyl-tRNA synthetase [EC:6.1.1.15] | 504 |
| unique/191/2/Org2_Gene107 (539) | Genetic Information Processin | RNA degradation | K04077 | groEL, HSPD1; chaperonin GroEL [EC:5.6.1.7] | 705 |
| unique/228/2/Org2_Gene324 (509) | Genetic Information Processin | RNA degradation | K05592 | deaD, cshA; ATP-dependent RNA helicase DeaD [EC:3.6.4.13] | 463 |
| unique/251/2/Org2_Gene1955 (499) | Genetic Information Processin | Replication and repair | K06919 | K06919; putative DNA primase/helicase | 178 |
| unique/284/2/Org2_Gene74 (484) | Genetic Information Processin | Aminoacyl-tRNA biosynthesis | K09698 | gltX; nondiscriminating glutamyl-tRNA synthetase [EC:6.1.1.24] | 456 |
| unique/419/2/Org2_Gene778 (430) | Genetic Information Processin | Aminoacyl-tRNA biosynthesis | K01893 | NARS, asnS; asparaginyl-tRNA synthetase [EC:6.1.1.22] | 426 |
| unique/427/2/Org2_Gene2001 (428) | Genetic Information Processin | Aminoacyl-tRNA biosynthesis | K01875 | SARS, serS; seryl-tRNA synthetase [EC:6.1.1.11] | 403 |
| unique/490/2/Org2_Gene1375 (407) | Genetic Information Processing | | K02600 | nusA; transcription termination/antitermination protein NusA | 362 |
| unique/680/2/Org2_Gene1394 (349) | Genetic Information Processin | Replication and repair | K03553 | recA; recombination protein RecA | 329 |
| unique/755/2/Org2_Gene511 (329) | Genetic Information Processin | Aminoacyl-tRNA biosynthesis | K01867 | WARS, trpS; tryptophanyl-tRNA synthetase [EC:6.1.1.2] | 313 |
| unique/933/2/Org2_Gene814 (295) | Genetic information processin | Chromosome and associated prot | K04763 | xerD; integrase/recombinase XerD | 262 |
| unique/951/2/Org2_Gene1366 (292) | Genetic information processin | Translation factors | K02357 | tsf, TSFM; elongation factor Ts | 266 |
| unique/1006/2/Org2_Gene1146 (281) | Genetic Information Processin | Replication and repair | K06223 | dam; DNA adenine methylase [EC:2.1.1.72] | 106 |
| unique/1010/2/Org2_Gene1536 (280) | Genetic Information Processin | Replication and repair | K07497 | K07497; putative transposase | 134 |
| unique/1040/2/Org2_Gene42 (273) | Genetic Information Processin | Transcription factors | K21900 | cysL; LysR family transcriptional regulator, transcriptional activator of the cy | 160 |

| | | | | | |
|---|---|---|---|---|---|
| unique/1104/2/Org2_Gene1365 (262) | Genetic Information Processin | Translation | K02967 | RP-S2, MRPS2, rpsB; small subunit ribosomal protein S2 | 258 |
| unique/1110/2/Org2_Gene1850 (261) | Genetic Information Processin | DNA replication proteins | K02315 | dnaC; DNA replication protein DnaC | 130 |
| unique/1233/2/Org2_Gene1464 (238) | Genetic Information Processin | Transfer RNA biogenesis | K09765 | queH; epoxyqueuosine reductase [EC:1.17.99.6] | 301 |
| unique/1265/2/Org2_Gene1019 (230) | Genetic Information Processin | Ribosome biogenesis | K06183 | rsuA; 16S rRNA pseudouridine516 synthase [EC:5.4.99.19] | 180 |
| unique/1470/2/Org2_Gene2216 (191) | Genetic Information Processin | Transcription factors | K02250 | comK; competence protein ComK | 158 |
| unique/1506/2/Org2_Gene1368 (184) | Genetic Information Processin | Translation factors | K02838 | frr, MRRF, RRF; ribosome recycling factor | 169 |
| unique/1511/2/Org2_Gene40 (184) | Genetic Information Processin | Replication and repair | K07486 | K07486; transposase | 52 |
| unique/1520/2/Org2_Gene1069 (182) | Genetic Information Processin | Ribosome biogenesis | K03817 | rimL; ribosomal-protein-serine acetyltransferase [EC:2.3.1.-] | 133 |
| unique/1623/2/Org2_Gene2244 (163) | Genetic Information Processin | Replication and repair | K07474 | xtmA; phage terminase small subunit | 114 |
| unique/1645/2/Org2_Gene1149 (159) | Genetic Information Processin | Ribosome biogenesis | K00783 | rlmH; 23S rRNA (pseudouridine1915-N3)-methyltransferase [EC:2.1.1.177] | 149 |
| unique/1673/2/Org2_Gene1374 (155) | Genetic Information Processin | Ribosome biogenesis | K09748 | rimP; ribosome maturation factor RimP | 136 |
| unique/1675/2/Org2_Gene1050 (155) | Genetic Information Processin | DNA repair and recombination pr | K03574 | mutT, NUDT15, MTH2; 8-oxo-dGTP diphosphatase [EC:3.6.1.55] | 122 |
| unique/1701/2/Org2_Gene39 (150) | Genetic Information Processin | Replication and repair | K07486 | K07486; transposase | 19 |
| unique/1711/2/Org2_Gene815 (149) | Genetic Information Processin | Transcription factors | K03711 | fur, zur, furB; Fur family transcriptional regulator, ferric uptake regulator | 204 |
| unique/1714/2/Org2_Gene752 (148) | Genetic Information Processin | Transcription factors | K23775 | ohrR; MarR family transcriptional regulator, organic hydroperoxide resistanc | 118 |
| unique/1717/2/Org2_Gene137 (148) | Genetic Information Processin | Chromosome and associated prot | K04047 | dps; starvation-inducible DNA-binding protein | 134 |
| unique/1741/2/Org2_Gene1395 (143) | Genetic Information Processin | RNA degradation | K18682 | rny; ribonucrease Y [EC:3.1.-.-] | 23 |
| unique/1778/2/Org2_Gene1236 (135) | Genetic Information Processin | Translation factors | K02358 | tuf, TUFM; elongation factor Tu | 45 |
| unique/1792/2/Org2_Gene481 (133) | Genetic Information Processin | Transcription | K16509 | spxA; regulatory protein spx | 127 |
| unique/1798/2/Org2_Gene1685 (132) | Genetic Information Processin | DNA repair and recombination pr | K03574 | mutT, NUDT15, MTH2; 8-oxo-dGTP diphosphatase [EC:3.6.1.55] | 102 |
| unique/1802/2/Org2_Gene2158 (131) | Genetic Information Processin | Replication and repair | K03469 | rnhA, RNASEH1; ribonuclease HI [EC:3.1.26.4] | 107 |
| unique/1808/2/Org2_Gene1575 (130) | Genetic Information Processin | Translation, Ribosome | K02996 | RP-S9, MRPS9, rpsI; small subunit ribosomal protein S9 | 128 |
| unique/1818/2/Org2_Gene1509 (128) | Genetic Information Processin | Replication and repair | K07476 | yusF; toprim domain protein | 106 |
| unique/1877/2/Org2_Gene1351 (116) | Genetic Information Processin | Translation, Ribosome | K02884 | RP-L19, MRPL19, rplS; large subunit ribosomal protein L19 | 108 |
| unique/1927/2/Org2_Gene418 (105) | Genetic Information Processin | Transcription factors | K03892 | arsR; ArsR family transcriptional regulator, arsenate/arsenite/antimonite-res | 62 |
| unique/1976/2/Org2_Gene1480 (94) | Genetic Information Processin | Translation, Ribosome | K02899 | RP-L27, MRPL27, rpmA; large subunit ribosomal protein L27 | 130 |
| unique/1977/2/Org2_Gene108 (94) | Genetic Information Processin | Chaperones and folding catalysts | K04078 | groES, HSPE1; chaperonin GroES | 116 |
| unique/2015/2/Org2_Gene118 (85) | Genetic Information Processin | Translation, Ribosome | K02909 | RP-L31, rpmE; large subunit ribosomal protein L31 | 125 |
| unique/2034/2/Org2_Gene1879 (80) | Genetic Information Processin | Translation, Ribosome | K02963 | RP-S18, MRPS18, rpsR; small subunit ribosomal protein S18 | 109 |
| unique/2180/2/Org2_Gene2175 (47) | Genetic Information Processin | Translation, Ribosome | K02913 | RP-L33, MRPL33, rpmG; large subunit ribosomal protein L33 | 35 |
| unique/2188/2/Org2_Gene1091 (45) | Genetic Information Processin | Translation, Ribosome | K02914 | RP-L34, MRPL34, rpmH; large subunit ribosomal protein L34 | 61 |
| unique/116/2/Org2_Gene2245 (642) | Glycan biosynthesis and metal | Other glycan degradation | K23989 | lytD, lytB; mannosyl-glycoprotein endo-beta-N-acetylglucosaminidase [EC:3. | 322 |
| unique/246/2/Org2_Gene415 (500) | Glycan biosynthesis and metal | Teichoic acid biosynthesis | K00712 | tagE; poly(glycerol-phosphate) alpha-glucosyltransferase [EC:2.4.1.52] | 312 |
| unique/628/2/Org2_Gene1321 (363) | Glycan biosynthesis and metal | Teichoic acid biosynthesis | K21285 | tagB, tarB; teichoic acid glycerol-phosphate primase [EC:2.7.8.44] | 258 |
| unique/1135/2/Org2_Gene1369 (256) | Glycan biosynthesis and metal | Peptidoglycan biosynthesis | K00806 | uppS; undecaprenyl diphosphate synthase [EC:2.5.1.31] | 277 |
| unique/112/2/Org2_Gene1791 (646) | Lipid metabolism | Glycerolipid metabolism, Glycan b | K19005 | ltaS; lipoteichoic acid synthase [EC:2.7.8.20] | 606 |
| unique/1113/2/Org2_Gene1370 (260) | Lipid metabolism | Glycerophospholipid metabolism | K00981 | E2.7.7.41, CDS1, CDS2, cdsA; phosphatidate cytidylyltransferase [EC:2.7.7.41 | 332 |
| unique/1137/2/Org2_Gene2233 (256) | Lipid metabolism/ Metabolism | Fatty acid biosynthesis/ Biotin me | K00208 | fabI; enoyl-[acyl-carrier protein] reductase I [EC:1.3.1.9 1.3.1.10] | 238 |
| unique/296/2/Org2_Gene1053 (474) | Metabolism of cofactors and v | Ubiquinone and other terpenoid- | K01911 | menE; o-succinylbenzoate---CoA ligase [EC:6.2.1.26] | 337 |
| unique/361/2/Org2_Gene264 (451) | Metabolism of cofactors and v | Biotin metabolism | K00833 | bioA; adenosylmethionine---8-amino-7-oxononanoate aminotransferase [EC | 435 |
| unique/566/2/Org2_Gene1786 (383) | Metabolism of cofactors and v | Folate biosynthesis | K01665 | pabB; para-aminobenzoate synthetase component I [EC:2.6.1.85] | 306 |
| unique/574/2/Org2_Gene1393 (381) | Metabolism of cofactors and v | Nicotinate and nicotinamide meta | K03742 | pncC; nicotinamide-nucleotide amidase [EC:3.5.1.42] | 138 |
| unique/734/2/Org2_Gene1052 (333) | Metabolism of cofactors and v | Ubiquinone and other terpenoid- | K02549 | menC; o-succinylbenzoate synthase [EC:4.2.1.113] | 203 |
| unique/970/2/Org2_Gene1458 (289) | Metabolism of cofactors and v | Pantothenate and CoA biosynthe | K00077 | panE, apbA; 2-dehydropantoate 2-reductase [EC:1.1.1.169] | 184 |
| unique/1041/2/Org2_Gene498 (273) | Metabolism of cofactors and v | Riboflavin metabolism | K21064 | ycsE, yitU, ywtE; 5-amino-6-(5-phospho-D-ribitylamino)uracil phosphatase [E | 238 |
| unique/1045/2/Org2_Gene1459 (272) | Metabolism of cofactors and v | Pantothenate and CoA biosynthe | K00606 | panB; 3-methyl-2-oxobutanoate hydroxymethyltransferase [EC:2.1.2.11] | 225 |
| unique/1047/2/Org2_Gene1742 (272) | Metabolism of cofactors and v | Ubiquinone and other terpenoid- | K01661 | menB; naphthoate synthase [EC:4.1.3.36] | 271 |
| unique/1081/2/Org2_Gene1796 (267) | Metabolism of cofactors and v | Ubiquinone and other terpenoid- | K08680 | menH; 2-succinyl-6-hydroxy-2,4-cyclohexadiene-1-carboxylate synthase [EC: | 199 |
| unique/1236/2/Org2_Gene1782 (237) | Metabolism of cofactors and v | Folate biosynthesis | K10026 | queE; 7-carboxy-7-deazaguanine synthase [EC:4.3.99.3] | 207 |
| unique/1303/2/Org2_Gene1784 (223) | Metabolism of cofactors and v | Folate biosynthesis | K06920 | queC; 7-cyano-7-deazaguanine synthase [EC:6.3.4.20] | 206 |
| unique/1304/2/Org2_Gene265 (223) | Metabolism of cofactors and v | Biotin metabolism | K01935 | bioD; dethiobiotin synthetase [EC:6.3.3.3] | 145 |
| unique/1422/2/Org2_Gene1787 (202) | Metabolism of cofactors and v | Folate biosynthesis | K02619 | pabC; 4-amino-4-deoxychorismate lyase [EC:4.1.3.38] | 145 |
| unique/1454/2/Org2_Gene1785 (195) | Metabolism of cofactors and v | Folate biosynthesis | K01664 | pabA; para-aminobenzoate synthetase component II [EC:2.6.1.85] | 180 |
| unique/1763/2/Org2_Gene1783 (139) | Metabolism of cofactors and v | Folate biosynthesis | K01737 | queD, ptpS, PTS; 6-pyruvoyltetrahydropterin/6-carboxytetrahydropterin synt | 143 |
| unique/475/2/Org2_Gene1525 (413) | Metabolism of other amino ac | Selenocompound metabolism | K11717 | sufS; cysteine desulfurase / selenocysteine lyase [EC:2.8.1.7 4.4.1.16] | 391 |
| unique/985/2/Org2_Gene1460 (286) | Metabolism of other amino ac | beta-Alanine metabolism | K01918 | panC; pantoate--beta-alanine ligase [EC:6.3.2.1] | 296 |
| unique/1816/2/Org2_Gene1461 (128) | Metabolism of other amino ac | beta-Alanine metabolism | K01579 | panD; aspartate 1-decarboxylase [EC:4.1.1.11] | 156 |
| unique/244/2/Org2_Gene1610 (501) | Metabolism of terpenoids and | Carotenoid biosynthesis | K10209 | crtN; 4,4'-diapophytoene desaturase [EC:1.3.8.2] | 399 |
| unique/247/2/Org2_Gene615 (500) | Metabolism of terpenoids and | Carotenoid biosynthesis | K10210 | crtP; diapolycopene oxygenase [EC:1.14.99.44] | 345 |
| unique/607/2/Org2_Gene616 (372) | Metabolism of terpenoids and | Carotenoid biosynthesis | K10211 | crtQ; 4,4'-diaponeurosporenoate glycosyltransferase [EC:2.4.1.-] | 189 |
| unique/1145/2/Org2_Gene1609 (255) | Metabolism of terpenoids and | Carotenoid biosynthesis | K10208 | crtM; 4,4'-diapophytoene synthase [EC:2.5.1.96] | 135 |
| unique/1628/2/Org2_Gene614 (161) | Metabolism of terpenoids and | Carotenoid biosynthesis | K10212 | K10212, crtO; glycosyl-4,4'-diaponeurosporenoate acyltransferase [EC:2.3.1. | 80 |
| unique/1222/2/Org2_Gene1367 (240) | Nucleotide metabolism | Pyrimidine metabolism | K09903 | pyrH; uridylate kinase [EC:2.7.4.22] | 340 |
| unique/1241/2/Org2_Gene136 (236) | Nucleotide metabolism | Purine metabolism | K03784 | deoD; purine-nucleoside phosphorylase [EC:2.4.2.1] | 272 |
| unique/1437/2/Org2_Gene117 (199) | Nucleotide metabolism | Pyrimidine metabolism | K00857 | tdk, TK; thymidine kinase [EC:2.7.1.21] | 183 |
| unique/316/2/Org2_Gene1527 (465) | Poorly characterized | | K09014 | sufB; Fe-S cluster assembly protein SufB | 469 |
| unique/457/2/Org2_Gene1524 (418) | Poorly characterized | | K09015 | sufD; Fe-S cluster assembly protein SufD | 371 |
| unique/691/2/Org2_Gene1122 (346) | Poorly characterized | Function unknown | K06871 | K06871; uncharacterized protein | 77 |
| unique/733/2/Org2_Gene281 (334) | Poorly characterized | Function unknown | K09190 | K09190; uncharacterized protein | 124 |
| unique/922/2/Org2_Gene2109 (298) | Poorly characterized | Function unknown | K00243 | K00243; uncharacterized protein | 267 |
| unique/986/2/Org2_Gene1516 (286) | Poorly characterized | Function unknown | K08974 | K08974; putative membrane protein | 254 |
| unique/1028/2/Org2_Gene1532 (275) | Poorly characterized | Function unknown | K07090 | K07090; uncharacterized protein | 208 |
| unique/1531/2/Org2_Gene287 (180) | Poorly characterized | Function unknown | K07089 | K07089; uncharacterized protein | 102 |
| unique/1569/2/Org2_Gene813 (174) | Poorly characterized | Function unknown | K09763 | K09763; uncharacterized protein | 86 |

| | | | | | |
|---|---|---|---|---|---|
| unique/1653/2/Org2_Gene1298 (158) | Poorly characterized | Function unknown | K09705 | K09705; uncharacterized protein | 150 |
| unique/1916/2/Org2_Gene1481 (106) | Poorly characterized | Function unknown | K07584 | ysxB; uncharacterized protein | 130 |
| unique/1981/2/Org2_Gene1376 (94) | Poorly characterized | Function unknown | K07742 | ylxR; uncharacterized protein | 81 |
| unique/2020/2/Org2_Gene1051 (83) | Poorly characterized | Function unknown | K08998 | K08998; uncharacterized protein | 91 |
| unique/31/2/Org2_Gene1145 (930) | Signaling and cellular processes | | K01153 | hsdR; type I restriction enzyme, R subunit [EC:3.1.21.3] | 716 |
| unique/68/2/Org2_Gene1136 (757) | Signaling and cellular processe | Prokaryotic defense system | K07016 | csm1, cas10; CRISPR-associated protein Csm1 | 641 |
| unique/213/2/Org2_Gene1143 (518) | Signaling and cellular processe | Prokaryotic defense system | K03427 | hsdM; type I restriction enzyme M protein [EC:2.1.1.72] | 452 |
| unique/301/2/Org2_Gene577 (472) | Signaling and cellular processe | Transport | K03319 | TC.DASS; divalent anion:Na+ symporter, DASS family | 403 |
| unique/346/2/Org2_Gene1110 (453) | Signaling and cellular processe | Transporter | K16211 | malY, malT; maltose/moltooligosaccharide transporter | 430 |
| unique/507/2/Org2_Gene1142 (400) | Signaling and cellular processe | Prokaryotic defense system | K01154 | hsdS; type I restriction enzyme, S subunit [EC:3.1.21.3] | 92 |
| unique/711/2/Org2_Gene1132 (340) | signaling and cellular processe | Prokaryotic defense system | K19140 | csm5; CRISPR-associated protein Csm5 | 205 |
| unique/714/2/Org2_Gene1840 (339) | Signaling and cellular processe | Prokaryotic defense system | K07451 | mcrA; 5-methylcytosine-specific restriction enzyme A [EC:3.1.21.-] | 27 |
| unique/722/2/Org2_Gene2097 (336) | Signaling and cellular processe | Transporters | K25282 | yclQ, ceuA; iron-siderophore transport system substrate-binding protein | 227 |
| unique/905/2/Org2_Gene1133 (302) | Signaling and cellular processe | Prokaryotic defense system | K19139 | csm4; CRISPR-associated protein Csm4 | 190 |
| unique/1099/2/Org2_Gene1128 (263) | Signaling and cellular processe | Prokaryotic defense system | K07451 | mcrA; 5-methylcytosine-specific restriction enzyme A [EC:3.1.21.-] | 139 |
| unique/1119/2/Org2_Gene1138 (259) | Signaling and cellular processe | Prokaryotic defense system | K15342 | cas1; CRISP-associated protein Cas1 | 195 |
| unique/1155/2/Org2_Gene1517 (253) | Signaling and cellular processe | Transporters | K09013 | sufC; Fe-S cluster assembly ATP-binding protein | 247 |
| unique/1198/2/Org2_Gene1130 (244) | Signaling and cellular processe | Prokaryotic defense system | K19091 | cas6; CRISPR-associated endoribonuclease Cas6 [EC:3.1.-.-] | 32 |
| unique/1300/2/Org2_Gene37 (224) | Signaling and cellular processe | Secretion system | K02242 | comFC; competence protein ComFC | 148 |
| unique/1319/2/Org2_Gene2152 (220) | Signaling and cellular processe | Transporters | K02003 | ABC.CD.A; putative ABC transport system ATP-binding protein | 170 |
| unique/1343/2/Org2_Gene2104 (215) | Signaling and cellular processes | | K02039 | phoU; phosphate transport system protein | 186 |
| unique/1346/2/Org2_Gene1134 (214) | Signaling and cellular processe | Prokaryotic defense system | K09002 | csm3; CRISPR-associated protein Csm3 | 161 |
| unique/1397/2/Org2_Gene1803 (205) | Signaling and cellular processe | Transporters | K06895 | lysE, argO; L-lysine exporter family protein LysE/ArgO | 141 |
| unique/1533/2/Org2_Gene807 (179) | Signaling and cellular processe | Transporters | K23675 | fmnP, ribU; riboflavin transporter | 139 |
| unique/1753/2/Org2_Gene1135 (141) | Signaling and cellular processe | Prokaryotic defense system | K19138 | csm2; CRISPR-associated protein Csm2 | 54 |
| unique/1891/2/Org2_Gene1335 (114) | Signaling and cellular processe | Transporters | K05567 | mnhC, mrpC; multicomponent Na+:H+ antiporter subunit C | 94 |
| unique/1952/2/Org2_Gene1137 (101) | Signaling and cellular processe | Prokaryotic defense system | K09951 | cas2; CRISPR-associated protein Cas2 | 53 |
| unique/553/2/Org2_Gene528 (388) | Signaling and cellular processe | ABC transporters, prokaryotic typ | K25143 | lnrM, sagH; linearmycin/streptolysin S transport system permease protein | 321 |
| unique/560/2/Org2_Gene279 (385) | Signaling and cellular processe | Drug transporters | K18567 | pbuE; MFS transporter, DHA1 family, purine base/nucleoside efflux pump | 85 |
| unique/692/2/Org2_Gene580 (346) | Unclassified: metabolism | Enzymes with EC numbers | K21417 | acoB; acetoin:2,6-dichlorophenolindophenol oxidoreductase subunit beta [E | 309 |
| unique/814/2/Org2_Gene579 (317) | Unclassified: metabolism | Enzymes with EC numbers | K21416 | acoA; acetoin:2,6-dichlorophenolindophenol oxidoreductase subunit alpha [ | 287 |
| unique/890/2/Org2_Gene522 (305) | Unclassified: metabolism | Enzymes with EC numbers | K03734 | apbE; FAD:protein FMN transferase [EC:2.7.1.180] | 234 |
| unique/1053/2/Org2_Gene2000 (271) | Unclassified: metabolism | Enzymes with EC numbers | K17758 | nnrD; ADP-dependent NAD(P)H-hydrate dehydratase [EC:4.2.1.136] | 295 |
| unique/1109/2/Org2_Gene602 (261) | Unclassified: metabolism | Enzymes with EC numbers | K00675 | nhoA; N-hydroxyarylamine O-acetyltransferase [EC:2.3.1.118] | 78 |
| unique/1618/2/Org2_Gene1651 (164) | Unclassified: metabolism | Enzymes with EC numbers | K24217 | mnaT; L-amino acid N-acyltransferase [EC:2.3.1.-] | 119 |
| unique/1754/2/Org2_Gene1503 (141) | Unclassified: metabolism | Enzymes with EC numbers | K04063 | osmC, ohr; lipoyl-dependent peroxiredoxin [EC:1.11.1.28] | 191 |
| unique/1799/2/Org2_Gene416 (132) | Unclassified: metabolism | Enzymes with EC numbers | K03741 | arsC; arsenate reductase (thioredoxin) [EC:1.20.4.4] | 142 |
| unique/1874/2/Org2_Gene1507 (117) | Unclassified: metabolism | Enzymes with EC numbers | K00537 | arsC; arsenate reductase (glutaredoxin) [EC:1.20.4.1] | 90 |
